# The Detailed 3D Multi-Loop Aggregate/Rosette Chromatin Architecture and Functional Dynamic Organization of the Human and Mouse Genomes

**DOI:** 10.1101/064642

**Authors:** Tobias A. Knoch, Malte Wachsmuth, Nick Kepper, Michael Lesnussa, Anis Abuseiris, A. M. Ali Imam, Petros Kolovos, Jessica Zuin, Christel E. M. Kockx, Rutger W. W. Brouwer, Harmen J. G. van de Werken, Wilfred F. J. van IJken, Kerstin S. Wendt, Frank G. Grosveld

**Affiliations:** Biophysical Genomics, Dept. Cell Biology & Genetics, Erasmus MC, Wytemaweg 80, 3015 CN Rotterdam, The Netherlands; Cell Biology & Biophysics Unit, European Molecular Biology Laboratory, Meyerhofstr. 1, 69117 Heidelberg, Germany; Cell Biology, Dept. Cell Biology & Genetics, Erasmus MC, Dr. Molewaterplein 50, 3015 GE Rotterdam, The Netherlands; Genome Organization & Function, BioQuant & German Cancer Research Center, Im Neuenheimer Feld 267, 69120 Heidelberg, Germany; Cohesin in Chromatin Structure and Gene Regulation, Dept. Cell Biology & Genetics, Erasmus MC, Dr. Molewaterplein 50, 3015 GE Rotterdam, The Netherlands; Center for Biomics, Dept. Cell Biology & Genetics, Erasmus MC, Dr. Molewaterplein 50, 3015 GE Rotterdam, The Netherlands

## Abstract

The dynamic three-dimensional chromatin architecture of genomes and its co-evolutionary connection to its function – the storage, expression, and replication of genetic information – is still one of the central issues in biology. Here, we describe the much debated 3D-architecture of the human and mouse genomes from the nucleosomal to the megabase pair level by a novel approach combining selective high-throughput high-resolution chromosomal interaction capture (T2C), polymer simulations, and scaling analysis of the 3D-architecture and the DNA sequence: The genome is compacted into a chromatin quasi-fibre with ∼5±1 nucleosomes/11nm, folded into stable ∼30-100 kbp loops forming stable loop aggregates/rosettes connected by similar sized linkers. Minor but significant variations in the architecture are seen between cell types/functional states. The architecture and the DNA sequence show very similar fine-structured multi-scaling behaviour confirming their co-evolution and the above. This architecture, its dynamics, and accessibility balance stability and flexibility ensuring genome integrity and variation enabling gene expression/regulation by self-organization of (in)active units already in proximity. Our results agree with the heuristics of the field and allow “architectural sequencing” at a genome mechanics level to understand the inseparable systems genomic properties.

## Introduction

The structure and function of genomes obviously co-evolved as an inseparable system allowing the physical storage, replication, and expression of genetic information (Eigen und Oswatitsch, 1981, Knoch, 1998, Cremer and Cremer, 2001). However, the dynamic three-dimensional higher-order architecture of genomes, their spatial and temporal modifications and/or relation to functional multi-dimensional interaction and regulatory networks have yet to be determined in detail (e.g. Cremer and Cremer, 2001; Knoch, 2002; Misteli, 2007; Jhunjhunwala et al., 2008; Rauch et al., 2008; Bickmore 2013; Belmont, 2014; Wachsmut et al., submitted). The DNA double helix and the nucleosome (Kornberg and Klug, 1973; Olins and Olins, 1974; Luger et al., 1997) have been determined in general structurally at the very highest level of detail including genome sequences and histone modifications. Additionally, it became apparent that genome organization and function indeed form a systems genomic entity (Cremer and Cremer, 2001; Knoch, 2003; Misteli, 2007; Knoch et al., 2009; Cremer and Cremer 2010; Bickmore 2013; Belmont, 2014; see also references within all these) responsible for gene expression (e.g. Bulger and Groudine, 2011; Kolovos et al., 2012) and forms the basis for individual differences and disease.

However, the immense size and structural complexity of genomes spanning many orders of magnitude impose huge experimental challenges and hence the higher-order architecture is still widely discussed. How nucleosomes are positioned, spaced, remodelled, and whether and how nucleosome chains fold into fibres at physiological salt concentrations have been matters of continuing debate (e.g. Maeshima et al., 2010): Finch and Klug (Finch and Klug, 1976) proposed a relatively regular solenoid and *in vivo* neutron scattering experiments revealed a compacted fibre with a diameter of 30±5nm as a dominant nuclear feature (Baudy and Bram, 1978; Baudy and Bram, 1979; Ibel, 1982; Notbohm, 1986). In contrast more recent suggestions range from no compaction at all (rev. Dubochet, 2012; Eltsov et al., 2008), to highly polymorphic nucleosome position- (Müller et al., 2014) and function-dependent compacted structures (Kepper et al., 2008; Stehr et al., 2010). The latter are essential to explain nucleosome concentration distributions (Weidemann at al., 2003; Wachsmuth et al., 2003; Capoulade et al. 2011), or chromatin dynamics (Belmont, 2001) and functional properties such as the nuclear diffusion of macromolecules (Knoch, 2002; Dross et al., 2009). Notably, the found fine-structured multi-scaling long-range correlation behaviour of the DNA sequence also predicts a compacted chromatin fibre (Knoch et al., 2002; Knoch, 2002; Knoch et al., 2009).

The higher-order chromatin architecture has been a matter of even greater debate: Pioneering light microscopy studies by Rabl (Rabl, 1885) and Boveri (Boveri, 1909) suggested a hierarchical self-similar, territory like organization. Electron microscopy suggested a more random interphase organization as in the models of Comings (Comings, 1968; Comings, 1978) and Vogel and Schroeder (Vogel and Schroeder, 1974). In the radial-loop-scaffold model of Paulson and Laemmli (Paulson and Laemmli, 1980) ∼60 kbp-sized chromatin loops attached to a nuclear matrix/scaffold should explain the condensation degree of metaphase chromosomes. According to Pienta and Coffey (1977; Pienta and Coffey, 1984), these loops persisted in interphase and formed stacked rosettes in metaphase. Micro-irradiation studies by C. Cremer and T. Cremer (Cremer at al., 1974) and fluorescence *in situ* hybridization (FISH) (Lichter et al., 1988; Cremer and Cremer, 2001) and studies thereafter finally confirmed a territorial organization of chromosomes, their arms, and stable sub-chromosomal domains during interphase, including their structural persistence during metaphase (de-)condensation (see Berezney et al., 2000; Misteli, 2007; Cremer and Cremer, 2010). The assumption since then has been that the ∼850 G, Q, R, and C ideogram bands (Franke, 1994) split into ∼2500 subchromosomal interphase domains. Chromatin rosettes explaining the (sub-)territorial folding were visualized by electron microscopy (Erenpreisa, 1989; Reznik et al., 1990) but remained unappreciated, whereas Belmont and Bruce proposed the EM-based helical hierarchy chromonema fibre (CF) model (Belmont and Bruce, 1994). Spatial distance measurements between small FISH-labelled genetic regions, led to the Random-Walk/Giant-Loop (RW/GL) model with the first analytical looped polymer description (Sachs et al., 1995; Yokota et al., 1995; Yokota et al., 1997). Here, 1 to 5 Mbp loops are attached to a non-protein backbone, following the line of Pienta and Coffey (Pienta and Coffey, 1984). Later, a combination of distance measurements by more structure preserving FISH, high-resolution microscopy, and massive parallel polymer simulations of chromosomes and entire cell nuclei, were only compatible with the rosette-like Multi-Loop-Subcompartment (MLS) model. In this model 60 to 120 kbp loops form rosettes connected by a similar sized linker (Knoch, 1998; Knoch et al., 1998; Knoch, 2002; Knoch, 2003; Jhunjhunwala et al., 2008; Rauch et al., 2008; Knoch et al., 2009). The MLS model is also in agreement with studies of transcription (e.g. Verschure, 1999) and replication (Berezney et al., 2000, and thereafter; Pope et al., 2014). *In vivo* FCS measurements of nucleosome concentration distributions and the dynamic and functional properties such as the architectural stability and dynamics of chromosomes (Knoch, 2002; Gerlich et al., 2003; Dross et al., 2009) or the diffusion of macromolecules (Knoch, 2002; Görisch, 2003; Dross et al., 2009) are essentially also in agreement with a small loop aggregate/rosette-like chromatin folding (Knoch et al., 1998; Knoch, 2002; Weidemann at al., 2003; Wachsmuth et al., 2003; Capoulade et al., 2011; Baum et al., 2014). Fine-structured multi-scaling long-range correlations of the DNA sequence again predict this (Knoch et al. 2002; Knoch, 2002; Knoch, 2003; Knoch et al., 2009).

However, to further investigate various aspects and to distinguish better between the different architecture proposals crosslinking techniques (used since the last century) were developed into a family of interaction capture techniques (Tab. S1) such as 3C (Dekker et al., 2002; Tolhuis et al., 2002), 3C-qPCR (Hagege et al., 2007), 4C (Simonis et al., 2006), 3C-seq/4C-seq (Stadhouders et al., 2013), 5C (Dostie and Dekker, 2007), and Hi-C (Liebermann-Aiden et al., 2009). They once more confirmed the existence of looping and subchromosomal domains (Dixon et al., 2012), now referred to as topologically associating domains (TADs) with a higher localization accuracy when compared to FISH. These led to a number of suggestions, such as the fractal globule model (Liebermann-Aiden et al., 2009), the loop array architecture in mitotic chromosomes (Naumova et al., 2013), and the highly dynamic loop formation based on single-cell (Giorgetti et al., 2014; compatible with a switch and binder model, Barbieri et al., 2012), or cell population experiments (Rao et al. 2014). However, these suggestions are not very well supported by the underlying experimental (raw) data and contrast many previous observations (see above). Nevertheless, whatever the suggested architectural model, these methods clearly showed, that physical interactions between functional elements proposed earlier (Müller-Storm et al., 1989; Carey et al., 1990; Hanscombe et al., 1991; see review Kolovos et al., 2012), are at the heart of genome function by regulating gene transcription. These often take place over huge genomic separations by direct contact via a preformed architecture and its modification (Jhunjhunwala et al., 2008; Rauch et al., 2008) or the formation of complexes such as in transcription factories (Iborra et al., 1996; Osborne et al., 2004; Kolovos et al., 2012; Ghamari et al., 2013). Additionally, more structural factors such as CTCF and/or cohesin play a role here (Zuin et al., 2014, and references therein), which seems obvious also from co-evolutionary considerations.

Here we use T2C, a novel selective high-throughput high-resolution chromosomal interaction capture developed by us (Knoch & Grosveld, 2013; Kolovos et al., 2014), which detects all probable physical genomic interactions (selective everything with everything) for a specific genomic region. Thus, it provides the means for efficient and cost effective “architectural genome sequencing” and allows to approach the major open questions discussed above with high quality: i) whether a chromatin fibre exists and how it is compacted, ii) how it is folded, iii) whether there is a general scaling behaviour of this architecture in agreement with the fine-structured multi-scaling long-range correlations of the DNA sequence organization, iv) whether this satisfies also the functional requirements with respect to the genomic life-cycle as well as dynamic *in vivo* properties, and v) whether all this is consistent with earlier experiments from a few to the mega base pair level. First we describe briefly the T2C design used here to investigate the human chromosome 11p 15.5-15.4 IGF/H19 locus, the mouse chromosome 7qE3-F1 β-globin region as well as 15 regions under different differentiation and function aspects basically from the base pair to the entire chromosome level. Then we show that T2C reaches the fundamental resolution limits where “genomic” statistical mechanics and uncertainty principles apply which is of fundamental importance for architectural T2C result interpretation. Thereafter, we show the high interaction frequency range, the reproducible detection of rare interaction events, and the high signal-to-noise ratio >10^5^-10^6^– all at the statistical limit. Then we further analyse these loci in terms of the 3D architecture which suggests that a chromatin quasi-fibre with ∼5±1 nucleosomes/11nm forms stable ∼30-100 kbp loops clustered into stable aggregate/rosette-like subchromosomal domains connected by a similar sized linker, with only minor but significant variations in the architecture in terms of cell types/functional states. In depth combination with supercomputer polymer simulations as well as scaling analysis of the 3D-architecture and the DNA sequence itself (where this architecture is represented by sequence specific “footprints”) results in the same conclusion and confirms the tight co-evolutionary entanglement between genome architecture and sequence. This is in excellent agreement with recent *in vivo* FCS measurements of the dynamics of the chromatin quasi-fibre and a there developed analytical polymer model (Wachsmuth et al., co-submitted). Consequently, T2C, polymer simulations, the DNA sequence organization, *in vivo* dynamic FCS measurements, and an analytical model are all agreeing. Since this is also consistent with the heuristics of the field, we finally conclude that this architecture, its dynamics, and accessibility balance stability and flexibility ensuring genome integrity and variation enabling gene expression/regulation by self-organization of (in)active units already in proximity.

## Results

### T2C a novel selective high-resolution high-throughput chromosome interaction capture

T2C is a selective high-resolution high-throughput chromosome interaction capture approach (Knoch & Grosveld, 2013; Kolovos et al., 2014) which we developed to design interaction capture studies in respect to their purpose – here efficient, high resolution/quality, and cost effective “architectural genome sequencing”. Briefly, T2C in this setup involves (Fig. 1A, details in Suppl. Methods): i) starting with ∼10^7^ cultured/prepared cells, ii) the cells are formaldehyde-fixed (i.e. all kinds of combinations of nucleic and protein crosslinks are formed), iii) permeabilized to allow intra-nuclear cutting with a first restriction enzyme, iv) extensively diluted to promote mono-molecular re-ligation reactions, before v) decrosslinking, purification, and final shortening of the DNA chimeric fragments to sizes < 500 bp by a second high-frequency restricting enzyme or by sonication. Then, vi) a region-specific DNA library of interacting fragments is produced using hybridisation to region specific arrays of DNA oligonucleotides, representing the end of each restriction fragment produced by the first restriction enzyme. With ∼10^9^ molecules of each hybridization-optimized oligonucleotide the capture is always in the linear range well below saturation relative to e.g. ∼10^7^ input cells. vii) After elution, the hybridised fragments are paired-end sequenced, and viii) each sequence pair is trimmed up to the first restriction enzyme and mapped to the whole reference genome. Only uniquely mapped sequences are used (eventually only between the two restriction enzymes). No other correction or cleaning resulting in information loss is performed due to the very nature of this method (see below). Thus, T2C has clearly several advantages in respect to study genome architecture in depths: i) It provides a choice between costs, resolution, interaction frequency range, size of the captured region, and multiplexing of samples in a study specific manner. E.g. a ∼500 bp average fragment resolution, in a 2 Mbp region, with six orders of magnitude interaction frequency range, and multiplexing of ten samples can be easily achieved sequencing 5 lanes. ii) The design of the oligonucleotide position ensures optimized data cleanness and high signal-to-noise ratio, allowing maximum interaction information with a minimum amount of sequencing (Fig. 1B-D; Tab. S1-3; Fig. S4-6). iii) Additionally, the process has been optimized for structure, and thus architectural preservation (Knoch, 1998; Knoch, 2002), minimal DNA loss during the procedure, and no use of signal amplification until sequencing when a limited number of PCR cycles could be performed (Tab. S1; Suppl. Methods).

**Figure 1:**
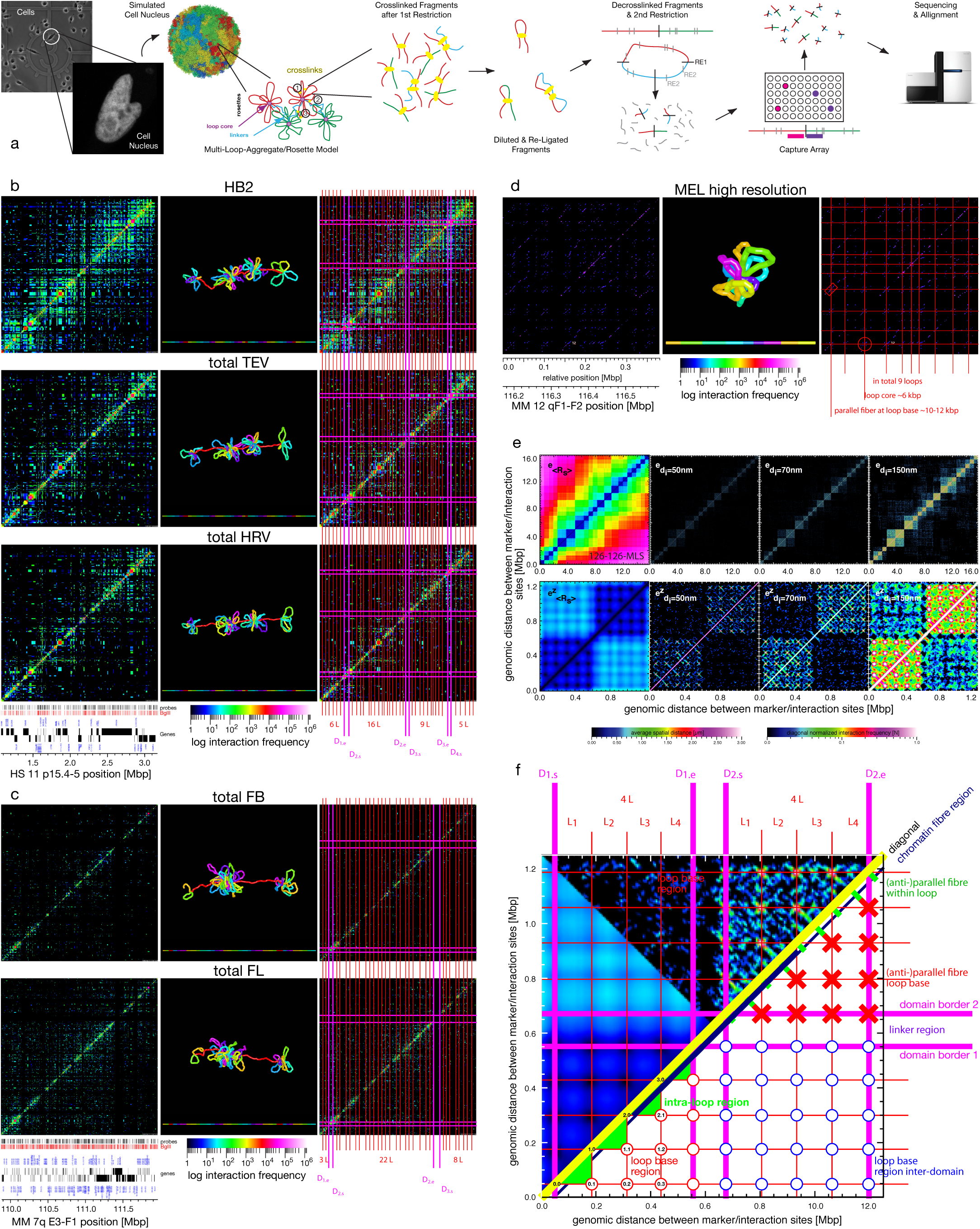
T2C description, interaction mapping, and direct determination of the chromatin quasi-fibre and the aggregated loop/rosette 3D architecture of the human and mouse genomes: **A**, Cell nuclei in a population of cells (transmission light and fluorescence microscopy, Fejes-Toth et al., 2004) have an underlying chromatin architecture (simulated cell nucleus containing 1.2 million polymer segments; resolution 5.2 kbp, i.e. ∼50 nucleosomes; Multi-Loop-Subcompartment (MLS) rosette model with 126 kbp loops and linkers; Knoch, 2002). After crosslinking the DNA is restricted within the nucleus by a 1st restriction enzyme, before the cross-linked fragments are extracted and diluted such that intra fragment re-ligation is favoured. After de-crosslinking, the re-ligated material is shortened by a 2nd restriction enzyme or sonication and purified by a capture array with oligos designed next to the 1st restriction enzyme, before paired-end-sequencing over the ligation position. After alignment to the reference genome, this results in interactions frequency matrices (B-D) and scaling curves (Fig. 2). **B, C**, Interaction matrices (logarithmic and colour coded scale; left & right) of the human IGF/H19 11p 15.5-15.4 region (B) in HB2, HEK293T TEV (intact cohesin) and HEK293T HRV (cleaved cohesin) as well as the mouse β-g 7qE3-F1 region (C) for fetal brain (inactive β-globin) and liver cells (active β-globin) show the formation of subchromosomal domains separated by a linker (borders: pink lines, right; D1s, D1e: start and end of domains), which consist of loops (red lines; 8L: number of loops), representing due to the grid-like pattern loop aggregates/rosettes. A grid-like pattern is also visible in the interactions between the domains and correspond to the interactions of loops and loop bases of interacting domains. Near the diagonal the aggregation into a chromatin quasi-fibre as well as loop internal structures are visible (zooming in- and out the images can make this clearer). Between different cell types or functional states only some local differences are visible resulting in a consensus architecture and allowing simulation of the 3D architecture (middle; resolution < ∼1 kbp). Note that the simulation is driven by the dominant consensus architecture. **D**, The interaction matrix of a 380 kbp subchromosomal domain in the mouse 12qF1-F2 region at high resolution clearly shows the regular rosette-like picture with a detailed structure of the loop base with in- and outgoing loop fibre stretches as seen in simulations (E, F). **E**, Simulated Multi-Loop-Subcompartment (MLS) model with an averaged spatial distance map for exact spatial distances 〈*R_s_*〉 (left) and on the diagonal normalized interaction maps for interaction radii 〈*d_i_*〉 of 50 nm, 70 nm, and 150 nm (right), for a MLS model with 126 kbp loops and linkers (16 Mbp upper and 1.2 Mbp zoom-in (z) lower row), showing clearly the formation of domains connected by a linker, their interaction, and the underlying loop aggregates/rosette architecture, with (anti-)parallel fibre stretches at the loop base. The dependence on the interaction radii corresponding to different crosslink probabilities is also clearly visible. **F**, Sketch of the different structures visible on different scales in the experimental and simulated interaction matrices (spatial distance matrix: left; simulated interaction matrix: upper (from E): On the smallest scale, near the diagonal the compaction of nucleosomes into the quasi-fibre (yellow line) and the fibre regime (dark blue line) can be found. On the largest scale the domains are clearly bordered (pink lines) and connected by a linker. On medium scales the loop aggregate/rosette-like structure is characterized by the loop bases (red circles: within domains, blue circles: between domains) as well as the loop interactions (green triangles). The fine-structure of the loops representing the (anti-)parallel loop stretches at the base (red crosses) and within loops (green stretches near diagonal).

**Figure 2:**
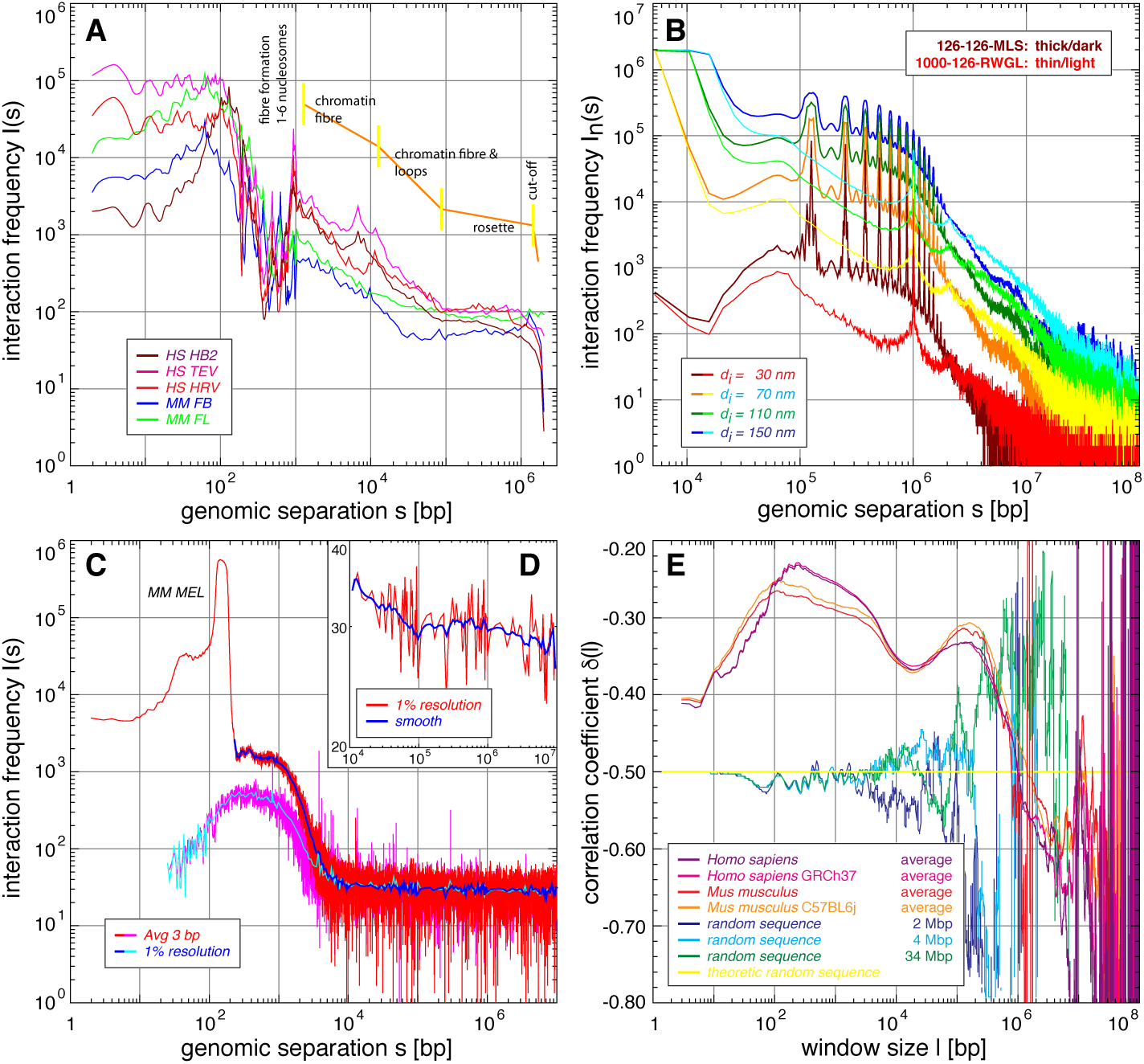
Scaling analysis of experiments, simulations, and the DNA sequence showing the formation of a chromatin quasi-fibre and the loop aggregate/rosette genome architecture: **A**, The fine-structured multi-scaling resulting from the T2C interaction frequency as a function of the genomic separation for the human IGF/H19 11p 15.5-15.4 region and the mouse β-globin locus 7qE3-F1 (3 bp average (1-200 bp) and thereafter a grouping with a 1% resolution per order of magnitude which for clarity is smoothed by a running window average for >10^3^ bp; see also Fig. S8; the values < 10 bp are due to the algorithm used and for transparency not discarded since they nevertheless show the extrapolation from values > 10 bp), shows: i) the structure of the nucleosome, ii) the formation of a plateau from 195 to ∼1000 bp, indicating the formation of a chromatin quasi-fibre with a density of 5±1 nucleosomes per 11 nm, iii) the chromatin quasi-fibre regime, iv) a mixed chromatin fibre/loop regime with a slightly higher interaction decrease, v) the plateau indicating the loop aggregate/rosette state, and vi) in principle the linker regime (not visible in A but in D). **C-D**, The fine-structured multi-scaling is even clearer for the average of 15 loci covering in total ∼99 Mbp in mouse MEL cells with subnucleosomal fragment resolution: After an initial increase a plateau is reached from ∼50 bp to ∼100 bp, followed by a sharp peak from ∼110 to 195 bp (width at plateau level ∼85 bp), followed by a second ∼10% decreasing plateau up to 1.0 to 1.2 kbp, which after a sharp decent until ∼10^4^ bp transits to the known multi-scaling behaviour (D, compare with A). With this resolution the fine-structure visible (Fig. S9), can be associated with the binding of the DNA double helix to the nucleosome, since up to ∼195 bp many of the small peaks (the most prominent at 145 bp) can be associated with the fine-structure in the fine-structured multi-scaling behaviour of DNA sequence correlations (E; Fig. S9, S21). Whereas the structure of the nucleosome vanishes using “secured” interactions (C, pink and light blue), above 195 bp the plateau and multi-scaling behaviour remains. Again the values the values < 10 bp are due to the algorithm used and for transparency not discarded since they nevertheless show the extrapolation from values > 10 bp. **B**, The interaction scaling of a simulated Multi-Loop-Subcompartment model with 126 kbp loops and linkers as well as a Random-Walk/Giant-Loop model with 1 Mbp loops and 126 kbp linkers consistently shows for different interaction radii a multi-scaling behaviour. The MLS model shows the characteristic rosette plateau, followed by the random scaling regime of the linker conducting a random-walk. The peaked fine-structure originates from the loops forming the rosettes. In contrast, the RWGL model is characterized by random-walk regime and only one major fine-structure attributable to the single loops. At greater scales the limit of the entire chromosome is seen in the cut-off. The MLS model agrees in detail with experiments (A, C-D) and the DNA sequence organization (E). **E**, The fine-structured multi-scaling long-range correlation behaviour of each of two human and mouse strains shows clearly again the architectural features: a general increase until a plateaued maximum (including the 145 bp peak), a first plateau area until ∼1200 bp, transition to a sharper decrease at ∼ 3.6 kbp (the sweet point used in the calculation of the persistence length) until a minimum ∼10-20 kbp and a second statistically significant maximum at ∼100 kbp, followed by a random regime and a final cut-off. The first maximum and plateau are characteristic for the nucleosome and formation of the quasi-fibre (C; Fig. S9, S21) which then transits to chromatin loops and their clustering into loop aggregates/rosettes which are connected by a random-walk behaving linker. Thus, due to the higher statistics here, the architectural features and their tight representation within the DNA sequence organization are even clearer.

To investigate the chromatin fibre conformation and the 3D genome architecture at the required resolution we chose the human chromosome 11p 15.5-15.4 IGF/H19 locus, and the mouse chromosome 7qE3-F1 β-globin region (Tab. S2). Both ∼2.1 Mbp regions have been well studied by FISH and other capture techniques. Using BglII or HindIII as 1st and NlaIII as 2nd restriction enzyme yields average fragment sizes of 3-6 kbp with many fragments, however, near the principle limit of the technique of a few base pairs (Fig. S3-6; average nucleosomal repeat length ∼195 bp; 3-6 kbp correspond to ∼15-30 nucleosomes). To determine the general chromatin fibre conformation at still higher resolution and to gain further insights into small scale architectural features, we also investigated 15 other regions (Tab. S2) covering in total 99.5 Mbp distributed over 10 different mouse chromosomes using ApoI as 1st restriction and sonication instead of a 2nd restriction leading to average fragment length of 549 bp (with many much smaller). This is even more at the technical limit and at nucleosomal/molecular resolution (Fig. S3-6). To investigate architectural and functional differences between species, cell lines, functional, and architectural differences, the human breast endothelial 1-7HB2 cell line (HB2), the HEK293T TEV/HRV RAD21-eGFP cell lines allowing cleavage of cohesin (Zuin et al., 2014), and mouse fetal brain and fetal liver (β-globin (in-)active) cells were used. To investigate the chromatin fibre conformation at high resolution undifferentiated murine erythroleukemia (MEL) cells were used.

### T2C reaches the fundamental resolution limits where “genomic” statistical mechanics and uncertainty principles apply

Since for “architectural sequencing” resolution is key, designing T2C using short fragment lengths down to even a few base pairs applying frequently cleaving restriction enzymes (Tab. S2; Fig. 1B-D; Fig. S3-6) not only molecular resolution (mind e.g. also the persistence length of free DNA ∼50 nm, i.e. ∼140 bp; typical protein/nucleosome binding regions are ∼100-500 bp) is reached and thus the fundamental limits of cross-linking techniques, but also the mechanism of observation is now, however, on the same scale as the observables (in *analogy* to classic and quantum mechanics). Actually due to the stochastics following the bias of the system behaviour, the observables, the observation, and thus the measured values are constrained by what we call “genomic” statistical mechanics with corresponding uncertainty principles. This originates from the individual complexity of each highly resolved interaction with a unique but coupled individual probabilistic fragment setting in each cell at a given time, e.g.: i) the cell population has a distribution of cell states and functional differences, ii) each fragment has a more or less dynamic individual DNA, RNA, protein, restriction association and length, and hence iii) a different crosslinking, restriction, re-ligation, oligonucleotide capture, sequencing, and mapping efficiency. The actual conditions and components can be determined only partially with high accuracy while with low accuracy otherwise and are eventually even entirely destroyed by the measurement. In essence, the entire T2C measurement process is highly quantitative but the local origin of this (including biases e.g. due to the oligonucleotide sequence or position), and thus its comparability, remains elusive due to its local individuality and the (at least current) incapability to determine all parameters linked in a complex network in detail and simultaneously as well as the attached biased system noise. Thus, the central limit theorem applies with an overlap of system inherent and real noise stochastics, and hence in the end only probabilistic analyses and statements can be drawn as hitherto is well known from classic mechanics, and more so from quantum (mesoscopic) systems. Consequently, population based or multiple single-cell experiments have to be interpreted and understood in a “genome” statistical mechanics manner with uncertainty principles due to the inseparability of factors/parameters also seen there. Thus, in practical terms, valid results are obtained when the statistical limit is reached, i.e. when scaling up the experiment does not narrow down the distribution any further and does not lead to fundamental (overall) changes anymore in observables. Due to the complexity involved, this has the immediate consequence that there are currently no means for adequate corrections. Even if certain biases might be known, the effect of a correction in terms of the many T2C steps remains illusive. This is the case for any interaction capture technique, although the effects of the individual complexity are partly averaged out by the lower resolutions mostly used in previous studies. This is no longer the case at the fundamental resolution limits. Nevertheless, if the statistical limit is reached and if the quality parameters like resolution, frequency range, and signal-to-noise ratio are sound, conclusions could be drawn as in the many cases of classic mechanics, and more so of quantum (mesoscopic) systems within the discussed boundaries.

### T2C reproducibly detects rare genomic interactions at the statistical limit with unprecedented signal-to-noise ratio

For the above mentioned experimental systems, with ∼10^7^ input cells, the corresponding samples (e.g. two different states) were multiplexed on the capture array to guarantee identical conditions (Tab. S3). Only sequences unique in the entire genome with a reasonably small mismatch rate (accounting for sequencing differences to and errors in the reference genome; see Suppl. Methods) and cleaned for sequences only mapping between the first and second restriction sites were analysed. Approximately ∼60-380 million paired-end sequencing reads were produced of which ∼10-65% were uniquely mappable (Tab. S3). The regional interactions (after normalisation for the total counts within the region) sorted and plotted in an upright squared interaction matrix/map with a logarithmic and rainbow colour-coded frequency range (Knoch, Lesnussa et al., 2009), including the diagonal (non- or self-ligation), shows directly the quality of the experiments and the unprecedented frequency range spanning 5-6 orders of magnitude (Fig. 1B-D; Fig. S4). Thus, also rare interactions with a frequency of 10^−4^ to 10^−6^ can be found and visualized under these conditions of region size, fragment resolution, and sequencing depth. We estimate an overall/cumulative (i.e. from cells to interaction matrix) efficiency of T2C of ∼0.1-1.0% from the ratio of cumulated counts per fragment to the number of input cells of ∼10^7^. The interaction patterns show, that the level of the stable statistical mechanical limit is reached, since data from different sequencing lanes or experiments (whether multiplexed or not) only show visually minor statistical deviations (Fig. 1B-C; Fig. S4). Quantitatively, also the statistical measures we used (e.g. frequency distributions) hardly show a change upon e.g. a two-fold increase of input cells or sequencing. At the statistical level reached, such a change leads only to an increase in novel interactions < 0.1%, mostly in the lowest interaction frequency regime. In contrast, a ten-fold sequencing decrease results in a massive interaction loss of > 25%. Most importantly, all the interaction matrices of different experiments are reproducibly mostly empty. Only ∼5-15% and 1.0-1.5% of the possible interactions show a signal for the IGF/H19 and for the β-globin, respectively (Tab. S3). Thus, there is no obvious uniform noise/background, despite the high number of sequence reads and the high number of diagonal elements showing entries of non- or self-ligated fragments. The “emptiness” is also structured, not arbitrary, and appears virtually the same in replicates, different cell types or functional states (Fig. 1B-D; Fig. S4-6). Moreover, interactions neither suddenly appear statistically nor cluster statistically somewhere near other or more prominent interactions. The signal-to-noise ratio is >10^5^-10^6^, even though noise could in principle appear at any step of the procedure, and even when assuming a highly unlikely biased distortion of a normal distributed noise signal towards e.g. interactions. A shot-noise (e.g. Poisson-like) analysis confirms this, in agreement with the change being < 0.1% during experimental scale-up (see above). Consequently, these values show that an analysis of this data in respect to genome architecture can be conducted within the limits of the above mentioned genome mechanical statistics constraints.

### The chromatin quasi-fibre forms stable loops clustered into aggregate/rosette-like subchromosomal domains connected by a linker

The interaction patterns (Fig. 1B-D; Fig. S4-6) can also be recognized clearly on all scales (within and between domains), including their re-emergence as attenuated repetition on other scales since genomes are scale-bridging systems (Knoch, 2002; Knoch, 2003). This behaviour shows once more the sensitivity of T2C allowing 3D architecture investigations despite the numerous and nonlinear parameters involved, since the probability that such repetitive patterns arise stochastically and even reproducibly is negligibly small relative to the number of those potentially formed combinatorically by hundreds of fragments. Additionally, T2C reveals agreement with other interaction techniques such as e.g. 4Cseq but with much cleaner and sharper interactions patterns for the same fragment setting (Fig. S5-6). In the following, the interaction patterns will be interpreted on the scales associated with the chromatin fibre, subchromosomal domains, and within the subchromosomal domains.

i. On the smallest genomic scale (Fig. 1B, C; Fig. S4-6), a dense and high interaction frequency pattern is observed in the region from 3 to 10 kbp (i.e. < ∼5-15, and ∼50 nucleosomes, respectively; for quantification, see scaling analysis below) along each point of the diagonal. This pattern varies independently of the local fragment size with distinct interactions and non-interacting “gaps” in-between. This is different from a homogenous random-walk or Rayleigh-like interaction “smear” decreasing uniformly and monotonously with increasing genomic separation. Additionally, the extension of the band of interactions is also smaller than a random-walk of nucleosomes would predict. A structurally uniform fibre like that seen in the (solenoid-like) helical chromatin fibre model (Finch and Klug, 1976) would result in a highly regular and defined pattern, which is also not observed. Thus, the pattern suggests, that there are defined stable interactions at the scale of DNA/nucleosomes forming an irregular yet locally defined and compacted structure. Hence, nucleosomes must form an irregular fibre, which we refer to as a “quasi-fibre” due to its inherent variation with average properties (e.g. an average linear mass density). While reading along the diagonal local interactions, compaction of nucleosomes, as well as other local properties of the chromatin quasi-fibre can be determined. The formation of such a quasi-fibre is in agreement with previous experimental results (Finch and Klug, 1976; Baudy and Bram, 1978), as well as simulations (Kepper et al., 2008; Stehr et al., 2010). This is also consistent with a variety of compacted structures described throughout the literature (see e.g. Kepper et al., 2008; Stehr et al., 2010), the absolute nucleosome concentration distributions (Weidemann at al., 2003; Wachsmuth et al., 2003), the dynamic and functional properties such as the architectural stability and movement of chromosomes (Knoch et al., 1998; Knoch, 2002; Gerlich et al., 2003; Dross et al., 2009; Baum et al., 2014), chromatin dynamics (Belmont, 2001), as well as the diffusion of molecules inside nuclei (e.g. Knoch, 2002; Dross et al., 2009; Baum et al., 2014). Moreover, recent genome wide *in vivo* FCS measurements of the chromatin quasi-fibre dynamics (Wachsmuth et al., co-submitted) also suggest such a chromatin quasi-fibre with variable, function-dependent properties. (See below for a quantification of T2C for the quasi-fibre properties).
ii. On the largest scale stable square-like domains (TADs; Dixon et al., 2012) are visible in the range of several hundred kbp to ∼1-1.5 Mbp, with clear borders and interactions with other domains (Fig. 1B-D; Fig. S4). They are more prominent e.g. in the IGF/H19 region, which shows two complete and two incomplete domains (Fig. 1B), when compared to the β-globin region with its single full domain and only two partially visible domains at the borders of the captured region (Fig. 1C). The domains feature several general properties: Firstly, the interaction frequency within domains has in general a higher average uniform height compared to interactions between domains, with a sharp drop at the edge of domains. The exact position of the border can be deducted from the folding within the domain and therefore respectively assigned exactly (see below). Thus, there is a cascade-like (average) behaviour of interactions with increasing genomic separation as predicted before (Knoch, 1998; Knoch et al., 1998; Knoch, 2002; Knoch, 2003), in contrast to the often expected general monotonous interaction decrease with growing genomic separation. Moreover, the interactions to other domains are clearly defined also in detail. Secondly, between the borders of the domains there is a clear transition or linker region, which again can be determined in respect to the folding of the chromatin quasi-fibre within the domain (see below). In and around these linker regions especially strong and complicated interactions are present depending on the specific domains. Such interactions originate from a combination of the facts that the chromatin quasi-fibre might not be shielded as is the case within the domains as well as the folding of the chromatin quasi-fibre itself (see below). A closer inspection of the interactions in the vicinity of the linker actually allows several interpretations in terms of the underlying domain architecture folding giving rise to these patterns. We favour that the genetic regions of the domains next to the linker interact more frequently compared to other domain parts due to the breaking of spatial isotropy. Two other possibilities that this is due to allelic differences (i.e. the patterns arise from two different domain architectures), or the linker being a very small linker domain consisting e.g. of a single or a few loops is much less likely (see below). A closer inspection of interactions near the linker in combination with the dynamic behaviour of subchromosomal domains (see dynamics below and Mov. S1-4) points also to a directionality along the “backbone” (the combination of several linkers of several subchromosomal domains), which is breaking the spatial isotropy of single unconnected subchromosomal domains. Consequently, these results confirm the existence of structurally stable subchromosomal domains which by (de-)condensation or (de-)looping explain the (de-)condensation of chromosomes through the cell-cycle (Pienta and Coffey, 1984; Erenpreisa, 1989; Knoch, 1998; Verschure, 1999; Berezney et al., 2000; Cremer and Cremer 2001; Knoch, 2002; Cremer and Cremer, 2010; Pope et al., 2014). The interaction pattern between subchomosomal domains and at their borders point already to a loop aggregate/rosette like architecture, since neither a free random-walk, an encaged random-walk, random or fractal globule like folding, nor a Random-Walk/Giant-Loop architecture would lead to sharp and defined borders. Instead they would lead to gradual/soft transitions instead. Constantly changing and thus very dynamic architectures with an average topology of these models or even that of a highly dynamic loop aggregate/rosette like architecture would also not result in the patterns found. This is in agreement with previous predictions on subchromosomal domains (Erenpreisa, 1989; Knoch, 1998; Berezney et al., 2000; Cremer and Cremer, 2001; Knoch, 2002; Jhunjhunwala et al., 2008; Rauch et al., 2008; Knoch et al., 2009; Cremer and Cremer, 2010; Pope et al., 2014). Moreover, these patterns are also in agreement with *in vivo* FCS measurements of the nucleosome concentration distribution (Weidemann at al., 2003; Wachsmuth et al., 2003), the dynamic and functional properties such as the architectural stability and movement of chromosomes (Knoch, 2002; Gerlich et al., 2003; Dross et al., 2009), chromatin dynamics (Belmont, 2001), as well as the diffusion of molecules inside nuclei (e.g. Knoch, 2002; Dross et al., 2009; Baum et al., 2014). Moreover, recent genome wide *in vivo* FCS measurements of the dynamics of the chromatin quasi-fibre, come to the same conclusion with characteristic functional differences (Wachsmuth et al., co-submitted). The intrinsic chromatin fibre dynamics with movements on the milli-second scale (Mov. S1-4) also points to the fact that the subchromosomal domains must have a stable architecture since otherwise they would dissolve immediately (see simulations below Wachsmuth et al., co-submitted). The break of the spatial isotropy of sequentially adjacent subchromosomal domains visible in the linker region is also linked to this stability.
iii. At intermediate scales within the subchromosomal domains, the interaction pattern is characterized by clearly distinct gaps and a crossed linear (grid-like) arrangement of interactions (Fig. 1B-D; Fig. S4-6). Interestingly, the linear pattern continues outside the subchromosomal domain and “crosses” with the linear pattern originating from the sequentially subsequent domain. Furthermore, the pattern outside is much simpler/clearer since it lacks the extra interactions originating inside the domain from e.g. the chromatin quasi-fibre, or its higher-order structure like e.g. intra-loop or loop-loop interactions (for illustration see Fig. 1E-F). This grid of interactions can also be quantified by projecting the interactions vertically and horizontally over the entire matrix, resulting in a peak-like pattern along the chromosome sequence (Fig. S7; see also Wachsmuth et al., co-submitted, for details). These peaks coincide with the grid-like pattern (Fig. S7). Projections within or outside the domains lead in essence to the same patterns with nevertheless subtle characteristic differences (see also Wachsmuth et al., co-submitted). Since interactions on scales of tens of kilo base pairs can only be due to chromatin looping, the conclusion must be that several consecutive loops have a coinciding loop base and hence form a loop aggregate/rosette like architecture. Hence, the interactions between subchromosomal domains result from the interactions of i) loops from domains next to each other, ii) loop bases of subsequent loop aggregates/rosettes when there is a relatively low density of loops, and iii) mitotic chromosomes present in the cell population. The borders of the domains seen on the medium scale (see above) are determined by the loops, and thus also the linker between subchromosomal domains is given by the end and start of loops of two subsequent subchromosomal domains. The border behaviour of domains near the linker was already discussed (see above). Determination of the loop positions and sizes (Tab. S4&5) visually as well as by projections (Fig. S7; for further details see also Wachsmuth et al., co-submitted) with an error on the level of corresponding local fragment resolution and in respect to the loop base structure of ∼3 kbp, reveals a consensus architecture independent of cell type or functional state with loop sizes of 48.6±14.5±2.4 kbp (average, StDev, StErr) and linker sizes of 46.7±15.1±8.7 kbp in the mouse β-globin region. In the human IGF/H19 locus the values are 57.8±16.2±2. kbp and 69.2±19.2±13.6, respectively. The subchromosomal domain sizes can now be calculated in detail for those subchromosomal domains which are completely covered by the T2C array: excluding the linker, the size is 1343.6±3 kbp for the single complete subchromosomal domain in the β-globin region, as well as 728.5±3 kbp and 403.4±3 kbp for the two complete subchromosomal domains of the IGF/H19 locus.

Although the ApoI T2C experiment was designed to elucidate the details of the chromatin fibre conformation only, one finds e.g. a 380 kbp subchromosmal domain region showing this pattern in greater detail (Fig. 1D). In addition to showing the same stable loop aggregate/rosette like architecture with 37.0±9.9±3.3 kbp loops (Table S6), and a subchromosomal domain size of 333.3 ± 3 kbp, part of the detailed loop base fine-structure with in and outgoing loop fibres spanning a region of ∼6 kbp can be seen (see simulations below; Fig. 1F; Fig. S11, S12).

The observation that the linear grid-like pattern outside of the domains is also not a homogeneous smear, shows that the loops and their arrangements into loop aggregates/rosettes is stable and not very variable. Once more the gaps between interactions, the grid-like pattern inside and outside the domains shows that a free random-walk, an encaged random-walk, a fractal globule like folding, or a Random-Walk/Giant-Loop architecture would lead to the patterns we find. Constantly changing and thus very dynamic architectures with an average topology of these or even that of a highly dynamic loop aggregate/rosette like architecture would not result in these patterns. Finally a non-compacted chromatin quasi-fibre, which a sea-of nucleosome organization predicts (Dubochet, 2012), would result in hugely homogeneous and very dynamic interaction possibilities, and thus patterns we do not find. Of course, the relatively simple notion of a quasi-fibre forming loop aggregates/rosettes connected by a linker becomes more complex due to the variation along the quasi-fibre, the variation of loop size and structure (e.g. super-helical topologies) and their arrangement either at the loop base or core of the loop aggregates/rosettes. Consequently, also on this architecture level the aggregate/rosette architecture also links interphase with metaphase very nicely and shows the architectural persistence during (de-)condensation within the replication process in agreement with experimental data (see Berezney et al., 2000, and thereafter). Moreover, this agrees with previous predictions on the internal structure of subchromosomal domains (Erenpreisa, 1989; Knoch, 1998; Berezney et al. 2000; Cremer and Cremer, 2001; Knoch, 2002; Jhunjhunwala et al., 2008; Rauch et al., 2008; Knoch et al., 2009; Cremer and Cremer, 2010; Pope et al., 2014). And again are also in agreement with *in vivo* FCS measurements of the nucleosome concentration (Weidemann at al., 2003; Wachsmuth et al., 2003) distribution and the dynamic and functional properties such as the architectural stability and movement of chromosomes (Knoch, 2002; Gerlich et al., 2003; Dross et al., 2009), chromatin dynamics (Belmont, 2001), as well as the diffusion of molecules inside nuclei (e.g. Knoch, 2002; Dross et al., 2009; Baum et al., 2014). Most importantly it agrees with the analysis of recent *in vivo* FCS measurements (Wachsmuth et al., co-submitted) showing similar loop sizes and loop numbers per subchromosomal domain. Thus, both T2C and the FCS *in vivo* measurements are in excellent agreement considering we investigate with T2C specific regions, in contrast to averaging over several regions in the FCS *in vivo* measurements, which suggests that this architecture occurs genome wide. We would like to stress again that the intrinsic chromatin fibre dynamics (on the milli-second scale) point to stable subchromosomal domains since the structure would otherwise dissolve immediately (see also simulations below; Mov. S1-4; and Wachsmuth et al., co-submitted).

### Comparison to the consensus 3D genome architecture shows small differences between species, cell type or functional state

To investigate how the genome architecture depends on species, cell type, functional or structural differences due to regulation or deliberate system distortion, we investigated the human IGF/H19 11p 15.5-15.4 region in human HB2, HEK293T TEV (intact cohesin), and HEK293T HRV (proteolytically cleaved cohesin) cells (Zuin et al., 2014), and the mouse β-Globin 7qE3-F1 locus in mouse fetal brain (FB; inactive β-Globin) and fetal liver (FL; active β-Globin) cells: As has been seen before (see introduction for any a 3C type assay) the subchromosomal domains are clearly very similar under different conditions (Fig. 1B, C; Fig. S4). The denser interaction pattern found in the HB2 cells when compared to the HEK293T cells may be due to differences in the level of crosslinkability. Comparing mouse FB to FL cells only shows subtle differences often belonging to single or a small group of interactions resulting from activation of the β-globin locus (Fig. 1C; Fig. S4, S6). Cleaving cohesion, which is thought to play a major constituative role in genome architecture, does not lead to dramatic changes on all scales despite some clear interaction losses and gains. Visual or quantitative determination of the loop positions also shows only minor differences (SFig. 7), which nevertheless might be functionally important. This might suggest that once formed, cohesion may not be required anymore to maintain the overall subchromosomal domain architecture. Thus, the detailed role of cohesion (as well as other factors like CTCF) in interphase chromatin remains unclear and needs to be clarified.

Consequently, these and other experiments from various labs as already mentioned show that organisms rely on a consensus architecture (overview in Cremer and Cremer, 2001; Cremer and Cremer 2010). This architecture has small functional variations on all scales from the chromatin quasi-fibre to the subchromosomal domains within the genomic regions. Between the subchromosomal domains, the architecture obviously varies more than within domains in agreement with recent FCS *in vivo* measurements (Wachsmut et al., co-submitted), where differences were found for different genomic regions or functional states such as eu- and hetero chromatin, or during massive changes by (de-)compacting the chromatin quasi-fibre by Trichostatin A or Azide treatment. The dynamics of the chromatin quasi-fibre on the millisecond scale in comparison to the size of the differences stresses again how stable this architecture is (see also simulations below; Mov. S1-4; and Wachsmuth et al., co-submitted). Hence, this illustrates the notion of the variation of a theme and points to the evolutionary balance between flexibility and stability of genome architecture in agreement with other findings/predictions (Erenpreisa, 1989; Verschure, 1999; Berezney et al., 2000; Cremer and Cremer 2001; Knoch, 2002; Knoch, 2003; Misteli, 2007; Jhunjhunwala et al., 2008; Rauch et al., 2008; Knoch et al., 2009; Cremer and Cremer, 2010; Bickmore 2013; Belmont, 2014; Pope et al., 2014). The biological implications of this are discussed below.

### Simulated polymer models *in silico* predict and confirm the genome organization in detail found by T2C

To better understand the above results and to evaluate hypotheses and plan future experiments concerning the 3D genome organization in eukaryotes we developed polymer models with pre-set conditions (i.e. without attempting to fit data; Knoch, 1998; Knoch et al., 1998; Knoch, 2002; Knoch, 2003; Jhunjhunwhala et. al., 2008; Rauch et al., 2008; Knoch, submitted; Knoch, submitted). The simulations (see Suppl. Methods) use a stretchable, bendable, and volume excluded polymer (hydrodynamic) approximation of the 30 nm chromatin fibre consisting of individual homogenous segments with a resolution of ∼1-2.5 kbp while combining Monte Carlo and Brownian Dynamics approaches. The simulations cover the Random-Walk/Giant-Loop model (Fig. S10) in which large individual loops (0.5–5.0 Mbp) are connected by a linker resembling a flexible backbone, as well as the Multi-Loop Subcompartment (MLS) model (Fig. S10) with rosette-like aggregates (0.5–2 Mbp) with smaller loops (60–250 kbp) connected by linkers (60–250 kbp). These models also contain enough information/aspects to cover other architectures such as free random-walks, random or fractal globules. They also cover their stability and dynamics. Two-dimensional spatial distance and interaction maps (for different crosslink conditions) were calculated with high statistical validity (Fig. 1E, F; Fig. S11, S12). Even in the absence of simulated nucleosomes, comparison of the simulations to our experimental results shows, that only an MLS- and thus loop aggregate/rosette-like genome architecture could explain all the above observations. Thus, also these simulations confirm previous predictions (see introduction; Erenpreisa, 1989; Pienta and Coffey, 1984; Knoch, 1998; Berezney et al., 2000; Cremer and Cremer, 2001; Knoch, 2002; Knoch, 2003; Jhunjhunwhala et. al., 2008; Rauch et al., 2008; Cremer and Cremer, 2010; Bickmore 2013; Belmont, 2014; Pope et al., 2014; Knoch, submitted; Knoch, submitted). Even slight variations of the simulated topologies (see Suppl. Methods; Tab. S7) are reflected in the simulation results: i) the interaction frequency in general depends on the spatial proximity and the (cell dependent) crosslink probability (here simulated by the crosslink kinetics in different volumes and thus interaction radii), ii) the appearance of the subchromosomal domains, subchromosomal domain separation, loops, and number of loops in a rosette (and thus density per subchromosomal domain) of the MLS model or loop size and separation in the RW/GL model are proportional to their size and number. The simulations also show more subtle effects for special combinations of parameters: i) high numbers of especially small loops in an MLS rosette result due the high density in steric exclusion and thus stretched loops eventually even “shielding” inner-rosette parts, or ii) inter-domain interactions are influenced by the connecting linker, loop size and numbers, and how non-equilibrium effects would appear (note: we deliberately show here not entirely equilibrated simulations; see also Knoch, 1998; Knoch, 2002; Knoch, submitted; Knoch, submitted). This also sheds light on the behaviour of domain borders near the linker (see above). The simulations support also the large emptiness of interaction matrices and its link to the existence of a dedicated chromatin quasi-fibre. Additionally, the simulations hint to a relatively low crosslink probability, radius, and frequency in experiments comparing the clearly visible fine-structure (such as the (anti-)parallel neighbouring of the chromatin quasi-fibre at loop bases; Fig. 1D-F) and its dependence on parameter changes in simulations.

The stability of the architecture with respect to the intrinsic chromatin fibre dynamics can also be illustrated by e.g. the decondensation from a mitotic chromosome into interphase (Mov. S1). This also shows that any 3D architecture would dissolve within seconds if it would not be stable. This agrees with the analytical polymer models developed recently to describe both structure and dynamics of the chromatin quasi-fibre (Wachsmuth et al., co-submitted). The subtle excluded volume effects or the fine structure of in/out-going loops at the loop aggregate/rosette core are not yet described to such detail in the analytical polymer model.

The *in silico* simulation approach was also used to visualize the 3D organization and its dynamics using the experimental interaction matrices as input. *In vivo* chromosomes never fold from scratch, they always have a history, and transform from one state to the other continuously. Thus, genomes are adiabatic systems, and hence we used here the consensus loop and domain positions (Tab. S4-6) as input starting conditions, rather than dropping a free linear polymer chain into the interaction landscape expecting it to fold in a defined knot-free 3D architecture. Only after a first Brownian Dynamics relaxation, the entire interaction matrix is used as additional side condition to introduce individual folding characteristics such as e.g. specific loop-loop interactions, which are not *a priori* contained implicitly in the consensus architecture. The outcome (Fig. 1B-D, middle) confirms that the chromatin quasi-fibre forms rosette-like subchromosomal domains with a high degree of agreement with the experiments and the analytic model mentioned above (Wachsmuth et al., co-submitted).

### Simulations show a fine-structured multi-scaling scaling behaviour revealing general aspects of genome organization

To comprehensively investigate and quantify the general behaviour of interactions as a function of genomic separation in a unified scale-bridging manner from a few to the mega base pair level, we already used scaling analysis to understand genome organization and showed its capabilities (see Suppl. Methods; Fig. S13; Knoch, 1998; Knoch, 2002; Knoch et al., 2009). Scaling analyses condense e.g. the two-dimensional interaction or spatial distance matrices into a one-dimensional scaling plot with higher statistical relevance to find signatures/patterns in a spectroscopy-like manner. For ease of understanding, we first introduce the scaling of the interaction frequency for the different simulated models (see Suppl. Methods; Fig. 2B; Fig. S14, S15): All MLS and RWGL simulations show clear long-range power-law scaling, with a multi-scaling behaviour and a fine-structure on top which are attributable to i) the general interaction decrease upon increasing spatial distance, ii) the subchromosomal domain or giant loop like structure, iii) the loop structure within the subchromosomal domains and/or the random-walk behaviour within the loops, and iv) the random-walk behaviour of the linker between the subchromosomal domain (i.e. the “backbone” behaviour of the entire chromosome), or that of giant loops. In the MLS models (Fig. 2B; Fig. S14) the clustered loops form a plateau with an ever steeper slope, which at scales > ∼10^6^ bp is dominated by the random-walk behaviour of the linker between the rosettes in agreement with polymer physics. The RWGL models (Fig. 2B; Fig. S15) are dominated by the random-walk behaviour within the mega-base pair loops and the linker between the loops. Obviously, the visible fine-structure is due to the loops, their size, and aggregation. The interaction itself and the pronounced fine-structure is inversely proportional to the interaction radius, i.e. to the spatial dynamics and kinetics of the crosslink. Again all model parameter variations are represented in the scaling behaviour on all scales in detail. The same holds for other scaling measures such as the spatial distance dimension or exact yard-stick dimensions directly measuring the scaling of the fibre folding (Fig. S13). The simulations show that there is no uniform scaling across all scales as e.g. seen in self-similar fractals bridging large orders of magnitude and hence this deviation shows the sub-structuring into loops and subchromosomal domains. Again this is in excellent agreement with the alternative analytical model (Wachsmuth et al., co-submitted). Consequently, scaling analysis is a relatively easy and adequate measure to analyse genome topologies in a scale bridging manner with a high degree of detail and statistical relevance.

### T2C shows fine-structured multi-scaling suggesting in detail loop aggregate/rosette 3D architectures

With this background we determined the experimental scaling behaviour (see Suppl. Methods) of the IGF/H19 locus, the β-globin region (Fig. 2A; Fig. S8) and that of the average of 15 regions in MEL cells (Fig. 2C, D; Fig. S9). In contrast to the simulations, the higher resolution of T2C allows a study of nucleosome compaction (see below), and whereas for the ∼2.1 Mbp regions the scaling cut-off is already reached at ∼10^6^ bp, the T2C design for the 15 regions reaches the cut-off later with a smaller frequency range for medium and large scales. For scales >10^4^ bp, all interactions clearly show fine-structured multi-scaling long-range power-law behaviour (Fig. 2A; Fig. S8), the details of which are only in agreement with the multi-loop aggregate/rosette like architecture (Fig. 2B; Fig. S14, S15) as predicted by us (Knoch, 1998; Knoch, 2002; Knoch, 2003; Knoch et al., 2009). The behaviour represents i) the general interaction decrease, i.e. spatial distance increase, of the chromatin quasi-fibre up to ∼3x10^4^ to 10^5^ bp, ii) the stable loop and aggregated-loop/rosette-like structure in the subchromosomal domains from ∼3x10^4^ up to 10^5^-10^6^ bp, iii) the subchromosomal-domain-like structure from ∼10^5^-10^6^ bp, and iv) the random-walk behaviour of the domain linkers above ∼0.8x10^6^ bp (until the cut-offs). As before the differences between species, cell type, or functional states are again small and the behaviour again shows the stability and functional variability of the system. We also found this scaling behaviour for Hi-C experiments of others (e.g. Liebermann-Aiden et al., 2009; Naumova et al., 2013; Giorgetti et al., 2014; Rao et al., 2014), suggesting the same 3D architecture (Imam et al., in prep.).

### High-resolution T2C scaling analysis reveals the detailed nucleosome structure and proves the formation of a chromatin quasi-fibre

Interestingly, we also get a dedicated fine-structured multi-scaling behaviour on scales from the base pair level up to 10^4^ bp (Knoch, 2002; Knoch, 2003; Knoch et al., 2009). This is especially true for the average of the scaling curves of the 15 regions totalling ∼99 Mbp regarding the high resolution of a few base pairs and the high statistical validity (Fig. 2C, D; Fig. S8, S9). Note that at such molecular scales i) the cleavage probability by the first T2C restriction enzyme depends critically on the local structure(s), ii) the crosslinked and (un)cleaved DNA would still be linked to the nucleosomal core and/or other factors, and iii) that this and the DNA fragment size (below or near the free DNA optimal end-to-end interaction probability), all influence the religation probability as well. All these factors alone or in combination may lead to the exploration of the entire possible spectrum of results. Nevertheless, distinguishing between all and “secured” interactions, i.e. including only sequentially non-adjacent neighbours and leaving out interactions based on uncleaved, i.e. non-religation events, confirms the validity of the results (Fig. 2C, D; Fig. S8, S9).

From a few base pairs onwards, the interaction frequency increases towards a plateau between ∼50 bp and ∼100 bp, followed by a sharp peak around 145 bp which is ∼1.5 orders of magnitude higher and ranges from ∼110 to 195 bp (width ∼85 bp). This is followed by a slightly decreasing plateau from ∼230 bp up to the transit to a new descent at ∼10^3^ bp which then obviously changes to a decay and at ∼10^4^ bp to the known multi-scaling behaviour for bigger scales (Fig. 2D). In the case of the lower-resolved IGF/H19 locus and the β-globin region, the plateau between ∼230 and ∼10^3^ bp is more a plateaued valley (Fig. 2A; Fig. S8) and again independent of species, cell type, or functional state. Taking “secured” interactions leaves out the behaviour below ∼195 bp but thereafter results in an increase to the plateau or plateaued valley at ∼230 bp up to the transit to the known descent at ∼10^3^ bp, with a hardly different behaviour thereafter (Fig. 2C; Fig. S8).

On top there is a dedicated fine-structure (Fig. S9), which up to ∼195 bp can be associated with the nucleosome and with the polymer behaviour of the nucleosomal chain thereafter: The most prominent feature is a small peak on the major peak at 145.5 bp (Fig. S9C), representing the winding of the DNA around the nucleosome, but other peaks are also where one would expect structural nucleosomal features (see below and Luger et al., 1997) and as we found them already by DNA sequence pattern analysis (see below; Knoch, 2002; Knoch et al., 2009). Actually, the T2C peaks from scales of 10 to 195 bp (the values < 10 bp in Fig. 1A, C, D; Fig. S8 are due to the algorithm used and for transparency not discarded since they nevertheless show the extrapolation from values > 10 bp) are equal to those found in nucleosomal binding sequences, the first plateau is associated with the nucleosomal linker sequence and the DNA only bound once to the nucleosome, the width of the main peak associates with the DNA sequence double wound around the nucleosome, and its upper border corresponds to the nucleosomal repeat length at 195 bp. On the second plateau from ∼230 to ∼10^3^ bp there might be a fine-structure as multiples of the 145.5 bp and the 195 bp nucleosomal repeat-length, e.g. at 290 bp as well as at 385 bp the peaks are exactly where dinucleosomal features are expected (Fig. 2C; Fig. S9B). Restriction enzyme dependencies might influence this as well although they seem to play only a minor or no role at all.

The plateaued valley at lower resolution and the plateau at high-resolution with a slight decrease of ∼10% for the interaction probabilities from nucleosome N1 to nucleosomes N4-N6, suggest that nucleosomes N4-N6 see the first nucleosome with nearly the exact same probability. Furthermore, for N7 the interaction decreases dramatically. Thus, each individual nucleosome has on average 4-6 clearly distinct nearest neighbour nucleosomes. This strongly suggests the formation of a chromatin quasi-fibre with an average density of 5±1 nucleosomes per 11nm, since the maximum distance between N1 and N2 can only be a stretched out DNA linker of ∼50 bp equalling ∼14 nm and for a crosslink this difference has to come down to a few Ås to form the covalent bond (neglecting the nucleosomal tails and the probability of crosslinking with a “stretched” DNA linker). Moreover, the genome wide *in vivo* FCS measurements of the dynamics of the chromatin quasi-fibre (Wachsmuth et al., co-submitted) show similar average quasi-fibre densities. We would like to stress once more that the interaction matrices show that this is only an average and thus for calling this a “quasi”-fibre.

### Apparent and average persistence length L_p_ of the chromatin quasi-fibre

To gain insight into the average mechanical properties of the chromatin quasi-fibre we calculated the average apparent persistence length L_p_ from the interaction scaling behaviour between 10^3^ and 10^4^ bp. For the calculation we assume the analytical model of an average quasi-fibre performing a (self-avoiding) random-walk (see theory in e.g. Knoch, 1998; Knoch, 2002; Wachsmuth et al., co-submitted) and importantly equal crosslinking kinetics for chromatin, and fit this to this region. This results in L_p_ between ∼1000 nm at ∼10^3^ bp down to 5 nm at ∼10^4^ bp. To gain a realistic value, one has additionally to take into account the notion that L_p_ is only defined above ∼2-3 times the basic fibre unit of 5±1 nucleosomes and for a free unlooped fibre, and thus well below the chromatin loop size, which in our case with an 10x oversampling means < 4 kbp. From the DNA sequence correlations (see below) this “sweet” point is at ∼3.6 kbp (Fig. 2E). Thus, at 3.6 kbp for 4-6 nucleosomes per 11nm, L_p_ ranges from ∼80 to 120 nm, respectively. This is in agreement with earlier values (see introduction; Kepper et al., 2008; Stehr et al., 2010) and with values derivable from spatial distance measurements between genetic markers (Knoch, 2002; Jhunjhunwala et al., 2008; Rauch et al., 2008; Knoch submitted). This average stiffness predicts that the average loop sizes will have to be on the scale seen above to ensure e.g. their stability, although, we would like to stress that this is only an average L_p_, and that L_p_ also varies with the quasi-fibre. Consequently, the notion of an apparent persistence length is only applicable in the context of a local L_p_, with a local meaning while taking the limit of the “genomic” statistical mechanics and uncertainty into account. This too agrees with values for L_p_ extractable from recent genome wide *in vivo* FCS measurements (Wachsmuth et al., co-submitted).

### The DNA sequence organization shows fine-structured multi-scaling long-range correlations tightly entangled with the 3D architecture

Since what is near in physical space should also be near (i.e. in terms of similarity) in DNA sequence space and this presumably genome wide (Knoch, 2002; Knoch, 2003; Knoch et al., 2009), and because evolutionary surviving mutations of all sorts will be biased by the genome architecture itself and vice versa, we also investigated the correlation behaviour of the DNA sequence (see Suppl. Methods; Knoch et al., 2002; Knoch, 2002; Knoch, 2003; Knoch et al., 2009; and references therein). To this end, we used the most likely simplest correlation analysis possible (to avoid information loss or being biased) and calculated the mean square deviation of the base pair composition (purines/pyrimidines) within windows of different sizes. In other words we calculated the function C(l) and its local slope δ(l), which is a measure for the correlation degree, or in more practical lay-men terms similar to a spectral measure (see Suppl. Methods; Knoch et al., 2002; Knoch, 2002; Knoch et al., 2009) for two different human and mouse strains (Fig. 1E; Fig. S16-21): i) long-range power-law correlations were found on almost the entire observable scale, ii) with the local correlation coefficients showing a species specific multi-scaling behaviour with close to random correlations on the scale of a few base pairs, a first maximum from 40 to 3.6 kbp, and a second maximum from 8×10^4^ to 3×10^5^ bp, and iii) an additional fine-structure in the first and second maxima is present. The correlation degree and behaviour is stronger in human compared to mouse, but within the species nearly identical comparing different chromosomes (with larger differences for the X and Y chromosome). The behaviour on all scales is equivalent concerning the different measures used (Fig. 2B; Fig. S13-15) to the long-range multi-scaling of the genome architecture with the transitions of behaviours even at similar scaling positions. Consequently, we can associate the behaviour with i) the nucleosome, ii) the compaction into a quasi-fibre, iii) the chromatin fibre regime, iv) the formation of loops, v) subchromosomal domains, and vi) their connection by a linker. Especially the transition from the basic nucleosomal compaction into the quasi-fibre regime at ∼3.6 kbp (what we called the sweet point in the calculation of the persistence length) is clearly visible. Additionally, on the fine-structural level, the already previously proven association to nucleosomal binding at the first general maximum (Knoch et al., 2002; Knoch, 2002; Knoch et al., 2009) is not only found again (Fig. S21), but also is in agreement with the fine-structure found in the interaction scaling (Fig. S9). This in principle holds now also for the fine-structure visible in the second general maximum as well and now even in the entire chromosome set and in two versions of two species and is associated with the loop aggregated/rosette structure there, predicting loop sizes from ∼30 to 100 kbp, and subchromosomal domain size from ∼300 kbp to ∼1.3 Mbp (see also Knoch, 2002; Knoch et al., 2009). We have, however, to acknowledge, that the experimental T2C data have only encompassed a small fraction of the genome yet, whereas the sequence correlations encompass the entire genome, which hints clearly that this is a genome wide phenomenon. Moreover, the existence and details of this behaviour show the stability and persistence of the architecture since sequence reshuffling or other destructive measures would result in a loss of this pattern. This would also be the case for an unstable architecture, which would not leave a defined footprint within the sequence. This is again in agreement with our simulations of the dynamics or the genome wide *in vivo* FCS measurements (Wachsmuth et al., co-submitted). Consequently, we show not only by two analysis of completely independent “targets” (the T2C interaction experiments and the analysis of the DNA sequence) the compaction into a chromatin quasi-fibre and a loop aggregate/rosette like genome architecture again, but prove here also the long discussed notion, that what is near in physical space is also near, i.e. more similar, in sequence space. Since all chromosome sequences show a highly similar behaviour this clearly shows the genome wide validity. Hence the 3D architecture and DNA sequence organization are coevolutionary tightly entangled (review of previous notions in Knoch, 2002 or Knoch, 2009). Thus, in the future from the DNA sequence and other higher-order codes (e.g. the epigenetic code) most architectural genome features can be determined, since also most structural/architectural features and vice versa left a footprint on the DNA sequence and other code levels as one would expect from a stable scale bridging systems genomic entity.

## Conclusions

Here we present the much debated 3D genome architecture and its entanglement with the DNA sequence from the single to the mega base pair level of the eukaryotic human and mouse genomes based on combining a novel selective high-throughput high-resolution chromosomal interaction capture (T2C), with a scaling analysis of the architecture as well as the DNA sequence organization, and polymer simulations. T2C has many a significant advantage, ranging from cost effectiveness, via a huge signal-to-noise ratio, to reaching the level of the “genomic” statistical mechanics with uncertainty principles. The latter is of major importance since here fundamental limits are reached with consequences for the setup and interpretation of experiments involving the architecture and dynamics of genomes. Actually, we face a situation very similar to the revolution in quantum mechanics brought about at the beginning of the 20th century. Thus, an entirely new way of thinking will be needed to further determine and understand the organization and function of genomes.

With this background, we show here i) the association of the DNA to the structure of the nucleosome core in detail and the existence of a chromatin quasi-fibre with an average of 5±1 nucleosomes per 11 nm and an average persistence length L_p_ from ∼80 to 120 nm, ii) the existence of stable chromatin loop aggregates/rosettes connected by a linker with loops and linkers ranging from ∼30 to 100 kbp (with details of the fibre folding at loop bases), iii) the existence of a consensus architecture with only small differences between species, cell type, or functional states likely to persist through the cell cycle, iv) the existence of fine-structured multi-scaling behaviour of the architecture, and last but not least that, v) the genome architecture is closely linked to the fine-structured multi-scaling long-range behaviour of the DNA sequence. This is a consistent scale-bridging systems picture of the 3D architecture, its dynamics, and functional variation of two mammalian genomes from the single base pair to the mega base pair level. All this is in agreement with many observations about the architecture, its dynamics, the diffusion of molecules, as well as the replication, storage, and expression of genetic information which have been made in the field (see introduction). Most interestingly, this is in agreement with novel genome wide *in vivo* FCS measurements of the chromatin quasi-fibre dynamics (Wachsmuth et al., co-submitted). Inevitably, there are still many an open question, such as the identification of the molecules (proteins, RNA, etc.) involved in looping, their dynamics, the inherent variability in the system, but what we present here provides now a framework for “architectural and dynamic sequencing” and the detailed analysis after all major architectural components in the human and mouse genome have been determined.

The implications of the architecture presented here, are many-fold of which we would like to mention a few: i) The balance between stability and flexibility of the whole system ensures that the overall genome integrity is maintained when local disturbance/damage takes place due to its modular build, while at the same time it allows fine adjustment of the architecture to enable the development of different gene expression programs/cell types. ii) The signals due to functional interactions do not stand out above those due to proximity, which is an intrinsic property of the loop aggregate/rosette-like folding of the genome. This suggests that the interaction of functional elements (both in respect to transcription as well as to replication) is achieved between fragments that are already in close proximity before their function is required. This proximity and being “tethered” in a subchromosomal domain increases the probability of interaction. iii) This architecture is open enough to allow the rapid diffusion of molecules such as transcription factors and also allow the movement of sequences to self-organize and form active and inactive units of the genome. These (and other) aspects together form an inseparable system giving rise to a functional genome.

## Acknowledgements

We would like to thank all the people who supported and influenced this work of T.A.K especially T. Weidemann, K. Fejes-Toth, M. Göker, R. Lohner, M. Stör, E. Spiess, K. Rippe, W. Waldeck, J. Langowski, C. Cremer, T. Cremer, K. Erenpreisa, A. Ollins, D. Ollins, A. Papantonis, P. R. Cook, C. C. Murre, J. Skok, K. Egger, and L. A. Knoch. We also thank the EpiGenSys virtual consortium lab for its input. This work was supported by ERASysBio+/FP7 and the nationals funding organizations (the Dutch Ministry for Science and Education, the Netherlands Science Organization, the UK Biotechnology and Biological Sciences Research Council, and the Bundesministerium für Bildung und Forschung (BMBF)). FGG thanks the SyBoSS initiative. We also thank the BMBF for its support under grant #01 KW 9602/2 (3D Human Genome Study Group, German Human Genome Project). JZ and KSW were supported by NWO ALW grant 821.02.014 and a TRR81 grant. For computational resources we thank the High-Performance Computing Center Stuttgart (HLRS; grant HumNuc), the Supercomputing Center Karlsruhe (SCC; grant ChromDyn), the Computing Facility of the German Cancer Research Center (DKFZ), the BMBF under grant #01AK803A (German MediGRID), and #01IG07015G (Services@MediGRID). We also thank all those institutions, universities, and companies providing us with ∼500.000 CPUh per day via computational grid resources: the German D-Grid, the European Grid Initiative EGEE, the Erasmus Computing Grid, the Almere Grid, EDGeS Grid, and all the unnamed computing grids we have access through via these. Especially, we would like to thank all the world-wide distributed unnamed donors of desktop computer power of our world-wide Correlizer@home BOINC grid!

## Author Contribution

TAK and FG have conceived the T2C method for the detailed analysis of genome interactions as early as 2004 and giving rise to the ERASysBio+/FP7 consortium. TAK conceived the systems genomic approach, the integrated analysis and coordinated it, made all figures, designs, carried out the systems genomic data integration, and wrote the manuscript. MW, AMAI, and FGG made contributions to the manuscript. TAK coordinated the EpiGenSys ERASysBio+/FP7 consortium, which is a major pillar of this project. FGG, KSW, and TAK directed the laboratory experiments: ML and TAK designed the oligonucleotides and capture arrays, PK and JZ did the experiments, CEK, RWWB, and WFJE, did the capture and sequencing, and ML, HJGW, RWWB, and AMAI mapped the sequencing reads. TAK made the initial simulations of the polymer models, and NK made the simulated interaction matrices and the T2C constrained polymer simulations. NK, ML, AMAI, and TAK made the interaction matrices, the visualization of results in the GLOBE 3D Genome Platform (created by ML, NK, TAK in cooperation with B. Eussen) and calculated the scaling of interactions both of experiments as well as simulations. AA and TAK calculated the sequence correlation data and were responsible for the creation of grids or their usage of the acknowledged grid infrastructures. MW and TAK determined the loop positions and matrix projections as well as analytical polymer analysis.

## Supplemental Methods

### HB2 Cell Line and Cell Culture

HB2 cells (1-7HB2, a clonal derivative of the human mammary luminal epithelial cell line MTSV1-7 (Bartek et al., 1991) were cultured in DMEM supplemented with 0.2 mM L-glutamine, 100 units/ml penicillin, 100 mg/ml streptomycin, 10% FCS, 5 µg/ml hydroxycortisone, and 10 µg/ml human insulin. In a previous 3C study we confirmed the karyotype and the DNA methylation of several regions (Nativio et al., 2009).

### Cohsin Cleavage HEK293T TEV/HRV Cell Line System and Cell Culture

The cleavable HEK293T TEV/HRV RAD21-eGFP stable cell line system (Schockel et al., 2011) is a HEK293T cell line system, which was transfected with a pRTS-1 vector (Bornkamm et al., 2005) encoding for a cleavable RAD21-eGFP fusion protein and an siRNA for endogenous RAD21 knock-down (Schockel et al., 2011; Zuin et al., 2014, and references in there). Both are expressed by doxycycline induced activation of a bidirectional promoter in-between and thus simultaneously. For the RAD2-eGFP fusion-protein, a cleavable RAD21, where the first RAD21-separase cleavage site is replaced by that of the 3C protease of the human rhinovirus (HRV protease) using a PCR-based mutagenesis (the second cleavage site remained unchanged to ensure less cell cytotoxicity) was inserted before eGFP. The tobacco etch virus protease (TEV protease), does not recognize the HRV cleavage site and thus can act as a control. The endogenous RAD21 knock-down sequence allows knock-down with the following 3’UTR-directed siRNA’s:

5′-ACUCAGACUUCAGUGUAUA-3′ (Scc1-1),

5′-AGGACAGACUGAUGGGAAA-3′ (Scc1-2).

For generating the HEK293T TEV/HRV RAD21-eGFP cell line system the original HEK293T cell line was cultured in DMEM supplemented with 0.2mM L-glutamine, 100 units/ml penicillin, 100 mg/ml streptomycin, 10% FCS, and was grown at 37°C and 5% CO_2_. For the transfection Lipofectamine 2000 (Invitrogen) according to the instructions of the manufacturer was used. Cells carrying the vector were selected by growth in 150 µg/mL hygromycin containing medium. Single clones were picked and analysed for expression of RAD21cv and RAD21wt constructs and depletion of the endogenous RAD21 three days after induction with 2 µg/ml doxycycline. The resulting HEK293T TEV/HRV RAD21-eGFP cell line was as well cultured in DMEM supplemented with 0.2mM L-glutamine at 37°C and 5% CO_2_.

To activate transgene expression with HRV (or TEV which serves as a control, thus a transfection takes place, but no cleavage) the cells were cultured for 3 days in the presence of 2 µg/ml of doxycycline. Thereafter, cells were split, reseeded until 50% confluency and transfected with HRV or TEV vectors using Lipofectamine 2000 (Invitrogen) again according to the instructions of the manufacturer. 24 hours after protease transfection the cells were used for the experiment.

### Cell Preparation from Mice

For mouse fetal liver and fetal brain cells, ∼10 embryos on day 12.5 of pregnancy from one to two transgenic FVB/N mice were used for the ∼10 million cells required by the experiment to have a complex enough cell population and enough DNA in the end to be sequenced: Mice were cleaned with 70% EtOH and the abdomen was opened to remove the cervix containing the embryos, before cutting them lose and removing them from the yolk sac and placenta. Small and underdeveloped embryos were discarded. The embryos are collected in petri dishes on ice with 0.5 ml 10% FCS/PBS. Then the fetal liver and brain were dissected from the embryos and collected in tubes (1 ml) on ice containing again 500 µl 10% FCS/PBS. The cells were then resuspended with a P1000/1ml plastic pipette tip and connective tissue was digested by adding 25 μl of a 2.5% Collagenase stock (0.125% end concentration) and incubated for ∼ 45 min at 37°C. Thereafter, the cell suspension was transferred to falcon tubes with 12 ml 10% FCS/PBS at room temperature and then was gently squeezed through a scraper mesh which was placed inside a 6-well dish using again a P1000/1ml plastic pipette tip. The mesh was washed with 2ml 10% FCS/PBS at room temperature to get all the cells from the mesh. The resulting single cell suspension was again collected in falcon tubes with an end volume of 12 ml 10%FCS/PBS at room temperature. Notably, we tried to keep the stress of the cells to the minimum, to avoid any damage to the cell nuclei. Both after the resuspension, the Collagenase treatment, and/or after the scraping of fetal liver and brain material/cells were spotted on glass slides to check for cellular and especially nuclear integrity by microscopy, with or without staining of the nucleus with DAPI (or eventually any other immunofluorescence or fluorescence *in situ* hybridization).

### T2C Crosslinking/Fixation of Cells

For crosslinking/fixation of the genome and the entire cells, the cells were first counted and their concentration adjusted to 10 million in 12 ml 10% FCS/PBS at room temperature and put into 15ml polypropylene tubes (used for cell culture and thus not excessively absorbing/fixing cells to the tube wall, Greiner Bio One). Then 650 µl of a 37 % formaldehyde PBS solution were added, i.e. an end-concentration of 1.9 % of formaldehyde was used for crosslinking/fixation, at room temperature for 10 min while gentle shaking to avoid cell aggregation. Note: the concentration of formaldehyde for crosslinking/fixation at this stage is ideal for the following steps and in respect to cell/nuclear integrity for the human and mouse cells we used here; although this might hold in general, cases are known where other concentrations and incubation times achieve better results. Thus, the tube was put on ice (from now on we kept everything on ice up to the 1st restriction of the DNA (see below) to avoid any damage of the material) and 1.6 ml of cold 1M Glycine in PBS were added to quench the crosslink/fixation. Thereafter, the cells were spun down for 8 min at 1300 rpm at 4°C, the resulting pellet was washed in ice-cold PBS, and taken up first in 1 ml before adding up to 14 ml of ice-cold PBS, followed again by spinning down for 8 min at 1300 rpm at 4°C. After discarding the supernatant, the pellet could now also be frozen for storage, but we advice to continue straight away with lysis and the 1st restriction. Again cells were spotted on glass slides to check for cellular and especially nuclear integrity by microscopy, with/without staining of the nucleus with DAPI (Note: again the cells could now also be used eventually for any other immunofluorescence or fluorescence in situ hybridization experiment).

### T2C Preparation of Cell Nuclei and 1st Nuclear Genomic DNA Restriction

For cell lysis and preparation of cell nuclei, 5 ml of an always freshly prepared (for full activity) lysis buffer on ice (!) consisting of 10 mM Tris pH 8.0 (50 μl 1M), 10 mM NaCl (10 μl 5M), 0,2% NP-40 (100 μl 10%), 100 μl 50x complete Prot. Inhib. mix (50X = 1 tablet in 1 ml PBS), and filled up with up to 5 ml MilliQ (4.74 ml) was used. The pellet prepared in the final step of crosslinking/fixation was taken up in 1 ml of this lysis buffer, resuspended and filled up with another 4 ml to a total of 5 ml and incubated for 10 min on ice. The now free cell nuclei were spun down for 5 min at 1800 rpm at 4°C, the pellet was taken up in 0.5 ml of ice-cold PBS in a safe-lock tube, and spun for 1 min at 2600 rpm a at 4°C. Again it is possible after removal of the supernatant to snap-freeze, and store the nuclei at −80°C. For a check we always spotted nuclei on glass slides to check for nuclear integrity by microscopy and/or staining of the nucleus with DAPI.

For the 1st restriction the nuclei were now resuspended in 0.5 ml/tubes with 1.2x restriction buffer (60 μl restriction buffer, 440 μl MilliQ and adjusted for BSA if necessary) and transferred to a 1.5 ml safe-lock tube. Then to gently permeabilize the nuclear lamina the tubes were put at 37°C and 7.5 μl of 20% SDS (0.3% endconcentration) were added, and incubated at 37°C for 1 h, while shaking at 900 rpm. After adding 50 μl of Triton-X-100 (2% endconcentration) for further gentle permeabilzing of the nuclear lamina, the nuclei were again incubated at 37°C for 1 h, while shaking at 900 rpm. Note: both the SDS and Triton-X-100 step need to be carried out with great care to avoid any decrosslinking – again we checked that by checking the nuclei microscopically with and/or without DAPI staining. For future controls of the undigested material (the so called 1st unrestricted control) now a 5 μl aliquot was taken and stored at −20°C. Then 400 units of the selected restriction enzyme was added and incubated over night (∼20 h) at 37°C. For the human cells in all cases the restriction enzyme BglII (Roche) was used. For the mouse cells we used either HindIII (Roche) or ApoI (New England Biolabs) was used. Note: even though its optimal temperature is 50°C, for ApoI 37°C should be used to prevent partial decrosslinking of the sample ;-). And again for future controls of the restriction now a 5 μl aliquot was taken and stored at −20°C (the so called 1st restriction control). After the 1st restriction 40 μl of 20% SDS (enconcentration 1.6%) was added to the remaining sample to stop the restriction and for further breakdown of the nuclear lamina by incubation at 65°C for 20-25 min, while being shaken at 900 rpm.

### T2C Dilution, Re-ligation and De-Crosslinking of Restricted Genomic DNA

Thereafter, the fully digested nuclear material was diluted by transferal to a 50 ml falcon tube and addition of 6.125 ml 1.15x ligation buffer (6.125ml: 5.421 ml MilliQ water + 704 µl ligation buffer). Then 375 μl of 20% Triton-X-100 (endconcentration 1.0%) was added and incubated in a 37°C water bath for 1 h, while shaking every 10 min by hand. Then 20 μl Ligase HC 5U/μl (100 U in total, Roche) was added and incubated at 16°C over night (∼20 h) followed by an additional 30 min of incubation at room temperature. To de-crosslink the nonligated and ligated DNA 30 μl 10 mg/ml Proteinase K (in total 300 μg) was added and incubated at 65°C in a water bath over night (∼20 h). Again for future controls of the relegation and de-crosslinking now a 5 μl aliquot was taken and stored at −20°C (so called religation/de-crosslink control).

### T2C DNA Purification and 2nd (Re-ligated-)DNA Restriction/Sonication

For further treatment of the sample, first the DNA was purified by adding 30 ml 10 mg/ml RNAse (300 µg in total) and incubating at 37°C for 30-45 min, followed by brief cooling to room temperature and addition of 7 ml phenol-chloroform and vigorous shaking. Then the sample was centrifuged at 4,000 rmp (2200xg) for 15 min, before the upper phase was put in a new 50 ml tube and 7 ml of MilliQ was added as well as 1 ml of glycogen per ml, 1.5 ml of 2M Sodium Acetate pH 5.6, and add 35 ml of 100% ethanol to enhance the purification, gently but thoroughly mixed and thereafter put at −80°C for 1.5-3 h. This was followed by direct centrifugation at 4,000 rmp (2200xg) for 15 min, supernatant removal, addition of 10 ml of 70% EtOH, resuspension, and again centrifugation at 4,000 rmp (2200xg) at 4°C for 15 min. After supernatant removal, the pellet was dried for 20 min and was dissoluted in 150 ml of 10 mM Tris pH 7.5 at 37°C for 30 min. Again for future controls of the relegation and decrosslinking now a 5 μl aliquot was taken and stored at −20°C (so called 1st purification control).

Thereafter, the resulting re-ligated and de-crosslinked purified material was shortened by a 2nd restriction: First, to control the amount of DNA at this stage an aliquot of 1 μl was run alongside a reference sample of species-matched genomic DNA of known concentration on a 2% agarose gel. Then the DNA was adjusted in 0.5 ml/tubes to a 100 ng/µl concentration and restricted with the 2nd restriction enzyme by adding 1 U per µg of DNA of the selected restriction enzyme and incubated over night (∼20 h) at 37°C. For the human cells in all cases the restriction enzyme NlaII (New England Biolabs) was used. For the mouse cells we used either if HindIII was used as 1st restriction enzyme DpnII (New England Biolabs) or if ApoI was used as 1st restriction enzyme sonication with 10 cycles of 15 sec on and 45 sec off. Again for future controls now a 5 μl aliquot was taken and stored at −20°C (so called 2nd restriction control – this can be combined with the purification control from above).

### T2C Treatment of the Various DNA Controls

For controls of the integrity of the DNA at the different stages the following controls were used: i) 1st unrestricted control, ii) 1st restricted control, iii) re-ligation/de-crosslink control, iv) 1st purification control, and v) 2nd restriction/final purification control. These samples were controlled on a 2% agarose gel with corresponding plasmid DNA, which was restricted alongside, re-ligated and purified as external restriction control. For controls i)-iii) the aliquots were incubated with 10 µl Proteinase K (10 mg/ml) in 90 ml 10 mM Tris pH 7.5 at 65°C for at least 1 h. The DNA was purified by adding 3 µl 10 mg/ml RNAse and incubation for at 37°C for 30-45 min, followed by brief cooling to room temperature and addition of MilliQ up to 500 µl (∼400 ml) as well as 500 µl phenol-chloroform and vigorous shaking. Then the controls were centrifuged at 13,200 rmp for 15 min, 2 µl of glycogen per ml, 50 µl of 2M Sodium Acetate pH 5.6, and add 850 µl of 100% EtOH were added, gently but thoroughly mixed and snap-frozen before direct procession to centrifugation at 13,200 rmp for 20 min, followed by supernatant removal, addition of 1 ml 70 EtOH, centrifugation at 13,200 rpm at 4°C, renewed supernatant removal, pellet drying for 20 min and dissolution in 20 µl of 10 mM Tris pH 7.5 at 37°C for 30 min.

### T2C General DNA Whole Genome Sequencing Library Preparation

In general the *T2C* DNA fragment library was prepared for sequencing analysis on the Illumina Cluster Station and HiSeq 2000 Sequencer according to the Illumina TruSeq DNA protocol with <u>*enhancing modifications from us*</u> (www.illumina.com, TruSeq DNA sample prep LS protocol; part#15026489 Rev. C): i) purification of the DNA fragments, ii) end-repair to reach blunt end status, iii) 3’-end Adenylation to avoid chimera, iv) sequencing adapter ligation including eventual multiplexing step, and finally v) purification of the *T2C* whole genome sequencing DNA fragment library.

Therefore, first the concentration of the *T2C* DNA fragment library was measured again for fine tuning using 1 µl of material using Quant-it dsDNA broad range assay kit. Then the samples were split into 4 sets of 5 µg each of the *T2C* DNA fragment library and the following complete procedure done for each of these 4 sets of material:

i. To purify the *T2C* DNA library after the 2nd restriction AMPure XP beads (Beckman Coulter) were used by adding 1.8 µl AMPure XP beads per 1.0 µl of digested DNA. This was incubated at room temperature for 5 minutes, placed on the magnetic stand and incubated at room temperature for 5 minutes, and the supernatant was discarded without disturbing the beads. The beads were washed 2 times with freshly prepared 70% ethanol, placed at 37°C for 5 minutes to let the beads dry. Then the beads were resuspended in 50 µl PCR grade water and incubated at room temperature for 5 minutes, placed on the magnetic stand for 5 minutes, and finally 50 µl supernatant was transferred to a new tube. One microliter was finally loaded on an Agilent Technologies 2100 bioanalyzer using a DNA 1000 assay to determine the quality of the purified digested DNA.
ii. For end-repair of the *T2C* library, DNA fragments, since they were restricted or sonicated before with overhanging ends, 4 material sets were each in 50 µl transferred to a 96 well plate. Since no in-line control reagent to avoid contamination of the material was used, 10 µl of resuspension buffer were added, followed by 40 µl of end repair mix, and mixed thoroughly but gently pipetting the entire volume up and down 10 times. Then the plate was covered with a microseal ‘B’ adhesive seal and placed on the pre-heated thermal cycler at 30°C for 30 min. After removing the adhesive seal from the plate, first the AMPure XP beads were vortexed until they were well dispersed, and 160 µl (consisting of 136 µl of AMPure XP beads mixed with 24 µl of PCR grade water) were added to the wells and the entire volume was again pipetted thoroughly but gently up and down 10 times. After 15 min of incubation, the plate was put on the magnetic stand at room temperature for another 15 min until the liquid appeared clear. Then twice 127.5 µl of the supernatant was removed, and thereafter 200 µl of freshly prepared 80 % EtOH was filled into the well of the plate without disturbing the beads, incubated at room temperature for 30 sec and discarded again without disturbing the beads. This was repeated twice before drying of the plate for 15 min. Only thereafter the plate was removed from the magnetic stand and the pellets resuspended with 17.5 µl of resuspension buffer, followed by 10 times thorough but gentle mixing by pipetting 10 times up and down. After incubation at room temperature for 2 min, the plate was put back on the magnetic stand at room temperature for 5 min again until the liquid appeared clear, and then 15 µl of the clear supernatant was removed containing the end-repaired material ready for the Adenylation of the 3’-ends in the next step.
iii. For 3’-end Adenylation of the end-repaired *T2C* DNA fragment libraries, i.e. to prevent the blunt ends from ligating to one another, and thus to ensure a low rate of chimera (concatenated template) formation during the adapter ligation reaction in step iv), Klenow exo enzyme in the presence of ATP was used. A corresponding single ‘T’ nucleotide on the 3’ end of the adapter provided a complementary overhang for ligating the adapter to the fragment. Therefore, 15 µl of the end-repaired *T2C* DNA fragment library were transferred to a new 0.3 ml PCR plate. Since the in-line control reagent to avoid contamination of the material was again not used 2.5 µl of the resuspension buffer was added, followed by 12.5 µl of thawed A-tailing mix, pipetted thoroughly but gently up and down 10 times. Then the plate was sealed with a microseal ‘B’ adhesive seal, and the plate was placed on a pre-heated thermal cycler at 37°C for 30 min. Immediately after removal of the plate from the thermal cycler, the adapter ligation took place.
iv. To ligate the sequencing adaptors using the Illumina provided indexed adapters #6 and #12, DNA adapter tubes and stop ligation buffer tubes were used, and centrifuged to 600 xg for 5 seconds. Immediately before use, the ligation mix containing tube was removed from the −25°C storage as recommended by Illumina. Since the in-line control reagent to avoid contamination of the material was again not used 2.5 µl of the resuspension buffer was added to the wells of another PCR plate, and 2.5 µl of the ligation mix was added as well. Then 2.5 µl from the appropriate adaptor tubes was added and thoroughly but gently pipetted up and down 10 times. Then the plate was sealed again with a microseal ‘B’ adhesive seal and the plate centrifuged to 280xg for 1 min. Thereafter, the plate was incubated on a pre-heated thermal cycler at 30°C for 10 min, the plate was taken down from the cycler, the adhesive seal removed, 5 µl of the stop ligation buffer was added, and thoroughly but gently pipetted up and down 10 times.
v. To purify the sequencer adapted *T2C* DNA fragment libraries again AMPure XP beads were used. Therefore, AMPure XP Beads were centrifuged until they were well dispersed and 42.5 µl of mixed AMPure XP Beads were added to the wells and thoroughly but gently pipetted up and down 10 times, before incubation at room temperature for 15 min. Then the plate was placed on the magnetic stand at room temperature for minimum 5 min or longer until the liquid appeared clear. Then 80 µl of the supernatant were removed from each well of the plate and while the plate remained on the magnetic stand, 200 µl of freshly prepared 80% EtOH were added without disturbing the beads, and incubated at room temperature for 30 sec. The complete supernatant was then removed. This EtOH wash was done twice, before the still on the magnetic stand resting plate was air-dried at room temperature for 15 min. After removal from the magnetic stand, the dried pellet was resuspended using 52.5 µl of resuspension buffer, and thoroughly but gently pipetted up and down 10 times. After incubation for 2 min, the plate was put back to the magnetic stand at room temperature for minimum 5 min or longer until the liquid appeared clear. Then 50 µl of the clear supernatant was transferred to a new 0.3. PCR plate for a second cleanup, and 50 µl of vortexed AMPure XP beads added, and thoroughly but gently pipetted up and down 10 times. Then the plate was again incubated at room temperature for 15 min, the plate was placed again on the magnetic stand at room temperature for minimum 5 min or longer until the liquid appeared clear. 95 µl of the supernatant were removed, while the plate still remained on the magnetic stand, 200 µl of freshly prepared 80% EtOH was added to each well without disturbing the beads, incubated at room temperature for 30 sec. The complete supernatant was then removed. This EtOH wash was done again twice, before the still on the magnetic stand resting plate was air-dried at room temperature for 15 min. After removal from the magnetic stand, the dried pellet was resuspended using 22.5 µl of resuspension buffer, and thoroughly but gently pipetted up and down 10 times. Again after incubation for 2 min the plate was put back to the magnetic stand at room temperature for minimum 5 min or longer until the liquid appeared clear. Finally 20 µl of the clear supernatant from each well of the plate were collected and the material from each of the 4 in parallel treated *T2C* DNA fragment libraries splits pooled.

### T2C Regional DNA Sequencing Capture Microarray Design

To achieve high-resolution and allow for high-throughput multiplexed sequencing and thus to achieve a highly relevant local interaction mapping, i.e. to achieve a high quality *T2C*, special capture arrays were designed to select specifically for genome regions of interest avoiding sequencing of unnecessary background, i.e. to create a region specific DNA sequencing library optimized for selection of the re-ligated DNA pieces after the 1st restriction, i.e. directly for interactions only in specific and relatively small genomic regions. Therefore, in a close cooperation with NimbleGen, we designed DNA oligos for 2.1M capture microarrays, i.e. capture microarrays capable of in principle fishing 2.1 million different genomic sequences with the same amount of different oligos. To achieve a real high quality result *T2C* only (!) one oligo was placed up- and one downstream as near as possible to the 1st restriction site used in the nuclear whole genome restriction, since the interest lies in sequencing just each side after re-ligation of this 1st restriction. The oligos were designed by NimbleGen and us for the selected regions of the human and mouse genomes using genome builts mm9 and HG19 with oligo length of 72±3 bp, unique appearance (no mismatch allowed) in the entire genome, and with respect to best and similar capturing, i.e. similar hybridization, on a microarray. Then the oligos were further selected: in the case of using a 2nd restriction enzyme to shorten the re-ligated DNA library for sequencing the oligo had to be situated between the 1st and 2nd restriction site. In the case of using sonication to shorten the re-ligated DNA only oligos within 150 bp of the 1st restriction site were chosen. If only one oligo was present, which crossed either the 1st or 2nd or even both restriction sites, only cuts at the oligo beginning or end of in total not more than 10 % were allowed, i.e. that the oligos could definitely capture DNA pieces with a minimum of 65 bp to guaranty specificity and similar hybridization efficiency. The same condition was applied in the sonication case, for the 1st restriction side. Thereafter, we mapped the oligos on the genome and controlled by hand whether the conditions were fulfilled and whether the oligos were properly placed in respect to other genome features. For production of the microarray the number of the 2.1 million possible different oligos *O_array_* was divided by the number of selected oligos *O_selected_* and then each selected oligo spotted *N_spotted_* times, with 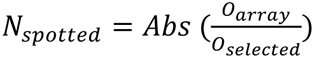, on the actual capturing array during the production process by NimbleGen. Thus, with the number of oligos used for capturing (see below) using a 1st and 2nd restriction enzyme we can be sure to have ∼10^10^ oligo molecules for each different oligo on the microarray, and thus with the 10^7^ cells we use as input, we are fare away from saturating the array by a factor of > 10^5^ to 10^6^. In the case of the experiment using sonication with ∼250 times more oligos and in total ∼50 times more genomic regions covered that is still > 10^2^, if considering the losses in the experimental procedures up to the capture array.

Concerning the experiments using a 1st and 2nd restriction enzymes, the balance between the region size chosen, the resulting size of the interaction matrix, i.e. the all possible interactions between all restriction fragments within this region, and the sequencing capabilities to achieve a high frequency range of a minimum of 4 to 5 orders of magnitude for each possible interaction (we assume an average of 2 to 3 orders of magnitude, which results in a spread of 4 to 5 orders of magnitude) was calculated. Thus, for a sequencing capability in two sequencing lanes of ∼300 and 500 million sequences, i.e. 300 and 500 million sequencings of possible interaction events, with the aim of achieving on average of 100 to 1,000 sequencing events per interaction 500 to 1,000 oligos and thus interaction fragments are optimal. The genomic region then covered depends only on the resolution, i.e. average spacing of the 1st restriction enzyme within the genome.

In the case of a 1st and 2nd restriction enzyme we chose the oligos and capture arrays as follows: In the human case this was done for the H19/IGF2 region on chromosome 11 from base pair position ∼1,110,650 to ∼3,216,350, i.e. a 2,105,700 bp sized region and 525 oligos. In the mouse case this was done for the β-globin region on chromosome 7 from base pair position ∼109,876,350 to 111,966,600, i.e. a 2,090,250 bp sized region and 800 oligos.

### T2C Regional DNA Sequencing Library Preparation - Microarray Capturing

To produce a subselected regional *T2C* DNA fragment sequencing library from the *T2C* whole genome DNA fragment sequencing library, the pooled DNA library after ligation of the sequencing adapters was subject to subselection with the above described newly and specifically developed capturing microarrays using the NimbleGen Array capture protocol and hybridization system with <u>*enhancing modifications from*</u> (www.nimblegen.com/seqcapez, NimbleGen Arrays User’s Guide, Sequence Capture Array Delivery version 3.2): The entire procedure consisted of i) microarray hybridization, ii) washing before iii) elution of the captured regional DNA library from the microarray.

i. Therefore, 3 h before the capturing, the hybridization system was set to 42°C, a first heat block was set to 95°C, and another one to 70°C, to equilibrate. Then the hybridization mixture was prepaired by adding 300 µl of 1 mg/ml Cot-1 DNA to the pooled DNA library after ligation of the sequencing adapters. In the case of using multiplexed samples not only the 4 sets of material were pooled but also the multiplexed samples were pooled. This saves microarray capacity and since the amount of DNA to be captured is far from the saturation of the microarray this leaves room for multiplexing up to 10 to 100 samples depending on the DNA amount, concentrations, and method to be used. Here multiplexing was only down by pooling 2 different materials. Then the sample was dried in a SpeedVac at 60°C for around 30 to 45 min, 11.2 µl of VWR water was added for rehydration, vortexed and centrifuged at maximum speed for 30 sec, before placement on the 70°C heat block for 10 min to fully solubilize the DNA. After a second vortexing and again centrifugation at maximum speed for 30 sec, 18.5 µl of 2X SC hybridization buffer and SC hybridization component A are added, followed again by vortexing and again centrifugation at maximum speed for 30 sec. Then to denature the DNA the sample was placed on the 95°C heat block for 10 min before another centrifugation at maximum speed for 30 sec. Thereafter, the sample was placed at 42°C and from there immediately loaded on the microarray hybridization chamber (the complete microarray system was prepared in parallel) and hybridized at 42°C for 64 h.
ii. To wash the captured regional *T2C* DNA library on the microarray first the elution chamber was assembled according to the NimbleGen array user guide. Therefore, the microarray slide was removed from the 42°C NimbleGene hybridization system and placed directly into the disassembly basin containing 100 ml of SC wash buffer II heated to 47.5°C. After ∼10 sec used for equilibration the mixer was peeled of and the slide was transferred to a second wash tube containing SC wash buffer II at 47.5°C, the sealed wash tube was inverted 10 times at a rate of 1 inversion per second. Then the slide was transferred to a new wash tube containing 32 ml of stringent wash buffer at 47.5°C, and the sealed tube was inverted 10 times at a rate of 1 inversion per second, before resting at 47.5°C for 5 min, and again inverted 10 times at a rate of 1 inversion per second. Then the slide was again transferred to a new tube containing 32 ml of stringent wash buffer at 47.5°C, and the closed tube was inverted 10 times at a rate of 1 inversion per second, before resting at 47.5°C for 5 min, and again inverted 10 times at a rate of 1 inversion per second. Then the slide was again transferred to a new tube containing 32ml of SC wash buffer I at room temperature, and the closed tube was inverted at a rate of 1 inversion per second for 2 min. Then the slide was again transferred to a new tube containing 32 ml of SC wash buffer II at room temperature, and the closed tube was inverted at a rate of 1 inversion per second for 1 min. Then the slide was again transferred to a new tube containing 32 ml of SC wash buffer III at room temperature, and the closed tube was inverted 10 times at a rate of 1 inversion per second.
iii. To elute the captured regional *T2C* DNA fragment sequencing library from the microarray the slide was transferred to the NimbleGen EL1 elution system at room temperature. Then ∼900 µl of 125 mM NaOH were added to the elution chamber until it is full, and incubated for 10 min. The eluted regional *T2C* DNA fragment sequencing library was pipetted to a 1.5 ml tube and filled up to 900 µl of 125 mM NaOH, followed by division equally in two new tubes containing 516 µl of a well mixed solution of 16 µl 20% acetic acid solution and 500 µl Qiagen Buffer PBI prepared beforehand in a 1.5ml tube. Then the mixture was transferred to a single MinElute column on a centrifuge to draw the solution through the column in several steps of 700 µl each. Then 750 µl buffer PE was put the column and centrifuged through. Then the MinElute column was put into a 2ml collection tube and centrifuged at maximum speed for 1 min. to remove any residual buffer PE. The flow-through was discarded, before placement of the MinElute column in a clean 1.5ml tube, 25 µl of buffer EB was added to the column, incubated for 1 min, and centrifuge at maximum speed for 1 min.

### T2C Amplification, Cluster Generation, and Paired-end High-Throughput Sequencing

First for paired-end sequencing the *T2C* regional DNA fragment sequencing library was enriched for sequencing first by PCR using Fusion polymerase using 30 sec at 98°C, 12 cycles of (10 sec at 98°C, 30 sec at 60°C, 30 sec at 72°C), 5 min at 72C final extension. For each 1 µg of *T2C* regional DNA fragment library 5 µl of the PCR primer cocktail and 25 µl PCR master mix was added to the PCR plate. For purification AMPure XP beads (Beckman Coulter) were used by adding 1.8 µl AMPure XP beads per 1.0 µl of DNA. This was incubated at room temperature for 5 minutes, placed on the magnetic stand and incubated at room temperature for 5 minutes, and the supernatant was discarded without disturbing the beads. The beads were washed 2 times with freshly prepared 70% ethanol, placed at 37°C for 5 minutes to let the beads dry. Then the beads were resuspended in 30 µl resuspension buffer and incubated at room temperature for 5 minutes, placed on the magnetic stand for 5 minutes, and finally 50 µl supernatant was transferred to a new tube. One microliter was finally loaded on an Agilent Technologies 2100 Bioanalyzer using a DNA 1000 assay to determine the quality of the purified digested DNA.

Cluster generation was performed according to the Illumina cBot User Guide (www.illumina.com, part#15006165 RevE). Briefly, 1 µl of a 10 nM TruSeq DNA library stock DNA was denatured with NaOH, diluted to 10 pM and hybridized onto the flowcell. The hybridized fragments are sequentially amplified, linearized and end-blocked according to the Illumina Paired-end Sequencing user guide protocol. After hybridization of the sequencing primer, sequencing-by-synthesis was performed using the HiSeq 2000 sequencer with a 101 cycle protocol according to the instructions of the manufacturer. The sequenced fragments were denaturated with NaOH using the HiSeq 2000 and the index-primer was hybridized onto the fragments. The index was sequenced with a 7-cycle protocol. The fragments were denaturated with NAOH, sequentially amplified, linearized and end-blocked. After hybridization of the sequencing primer, sequencing-by-synthesis of the third read was performed using the HiSeq 2000 sequencer with a 101-cycle protocol.

### T2C Sequence Mapping and Classification

The raw sequence reads were checked for the existence of the first restriction enzyme recognition sequence in the sequencing direction. The sequence after the first enzyme recognition site was removed. If the bases of the recognition site after the overhang were not unambiguously, the read was further trimmed by removing all the bases after the end of the overhang. These trimmed sequences were aligned using the Burrows-Wheeler Alignment (BWA) tool (Li and Durbin, 2009) to the whole human genome NCBI36/hg18 assembly and to the mouse NCBI37/mm9 assembly. Therefore the following default parameter set was used (with the value of the parameter in [] brackets): bwa aln [options] <prefix> <in.fq>

|  |  |
| --- | --- |
| -n NUM | max #diff (int) or missing prob under 0.02 err rate (float) [0.04] |
| -o INT | maximum number or fraction of gap opens [1] |
| -e INT | maximum number of gap extensions, -1 for disabling long gaps [-1] |
| -i INT | do not put an indel within INT bp towards the ends [5] |
| -d INT | maximum occurrences for extending a long deletion [10] |
| -l INT | seed length [32] |
| -k INT | maximum differences in the seed [2] |
| -m INT | maximum entries in the queue [2000000] |
| -t INT | number of threads [1] |
| -M INT | mismatch penalty [3] |
| -O INT | gap open penalty [11] |
| -E INT | gap extension penalty [4] |
| -R INT | stop searching when there are >INT equally best hits [30] |
| -q INT | quality threshold for read trimming down to 35bp [0] |
| -f FILE | file to write output to instead of stdout |
| -B INT | length of barcode |
| -L | log-scaled gap penalty for long deletions |
| -N | non-iterative mode: search for all n-difference hits (sloooow) |
| -I | the input is in the Illumina 1.3+ FASTQ-like format |
| -b | the input read file is in the BAM format |
| -0 | use single-end reads only (effective with -b) |
| -1 | use the 1st read in a pair (effective with -b) |
| -2 | use the 2nd read in a pair (effective with -b) |
| -Y | filter Casava-filtered sequences |

In case of using a second restriction enzyme (and thus not in the case of sonication) the unique sequences were aligned in a second step to a masked genome, excluding the sequence parts between second restriction enzymes that did not contain a first enzyme recognition site. Finally, only those sequences were paired using the SAMtools (Li et al., 2009) to generate paired-end Binary Alignment/Map (BAM) files, which showed in both the whole and masked genome reference sequences a unique alignment. Note: the alignments are unique, but nevertheless contain mismatches etc., which are either do to sequencing errors or hint a difference of our cells/mice to the reference genome. Unfortunately, there is also no way of distinguishing false positive or false negative alignments. Consequently, the resulting paired-end sequences then contain the interaction information with an error rate determined by the error rate of sequencing, the quality of the reference sequence, and the difference the DNA sequence of our cells/mice to this reference genome. A rough estimate of the false positive and false negative results for unique sequences without mismatches at the end of this process using known error rates indicate the error to be smaller than 10^4^ after accumulation of errors and the reduction of errors due to our procedure. This can be also deducted from the reduction of sequence pairs from the initial raw sequence throughout the entire process to the final result. Besides, omitting neighbouring interaction fragment pairs, leads to “secured” data sets where we can be sure that the result is not based on a unresctricted or restricted followed by re-ligation event.

### Computer Simulations

Using a two-step combined Monte Carlo and Brownian Dynamics method, the Random-Walk/Giant-Loop (RW/GL) model (Sachs et al., 1995) and the Multi Loop Subcompartment (MLS) model (Knoch, 1998; Knoch et al., 1998; Knoch, 2002; Knoch, 2003; Jhunjhunwhala et. al., 2008; Rauch et al., 2008; Knoch, submitted; Knoch, submitted) were simulated for human interphase chromosome 15. Assuming a flexible polymer chain, the chromatin fibre was split into ∼3,300 segments of 300 nm (∼31 kbp). To each segment a harmonic stretching potential and between two segments a harmonic bending potential were assigned. To avoid self-crossing of the polymer chain, a short ranged excluded volume potential was introduced, whose potential barrier could be changed to facilitate chain dis-entanglement to speed up simulations. *In vivo* this is mediated by Topoisomerase-II especially during chromosome de- or condensation. The simulation of single chromosomes necessitated placement into an embedding potential simulating the surrounding nuclear chromosomes. Different models were simulated (Tab. S7): In the RW/GL model loops of 5.0, 4.0, 3.0, 2.0, 1.0, 0.504 and 0.252 Mbp were used and connected by a chromatin linker whose length was adjusted such that the global territory behaviour yielded comparable results as in the MLS model. In the MLS model the loop size was 126 kbp with linker sizes of 63, 126, 189 and 252 kb. The number of loops in the rosettes varied, since the DNA content of the rosettes was assumed to be that of the metaphase ideogram banding pattern (Franke, 1994) divided by three to account for the transition into interphase ideogram bands (Yunis, 1981). The starting configuration of a chromosome had the approximate form and size of a metaphase-chromosome whose de-condensation resembles the natural process. Typically ∼400,000 Monte Carlo steps were needed to generate enough statistically independent configurations at thermodynamic equilibrium. For comparison with experimental spatial distance measurements between genetic markers as a function of their genetic separation, and the here interesting spatial distance and interaction maps, 100 to 150 statistically independent Monte Carlo configurations were taken as starting points for relaxation at higher spatial resolution by Brownian Dynamics methods using a decreased segment length of 50 nm (∼5.2 kb), corresponding to 20,000 segments for chromosome 15. 2,000 Brownian Dynamics steps were performed until equilibration was reached again. To simulate the dynamics for illustration purposes, as e.g. for the decondensation from a metaphase starting configuration into interphase, the resolution was set right away to the decreased segment length of 50 nm (∼5.2 kb; Mov. S1) or 20 nm (∼2.0 kbp; Mov. S2-4) and only Brownian Dynamics steps of 10 ns were performed.

### Simulated Spatial Distances, Spatial Distance and Interaction Maps

The simulated spatial distances were determined position dependently, i.e. the marker pairs were placed in respect to the topological folding of the simulated chromosome and position independently, i.e. the pairs of markers were placed randomly and therefore regardless of any folding topology on the chromosome (for the underlying assumption see results). All pairs of positions for one genomic separation were taken, the resulting spatial distance determined and averaged over 100 to 150 statistically independent chromosome configurations. For two-dimensional spatial distance and interaction maps the segments were divided by a factor of 10, i.e. a resolution of ∼520bp. For two-dimensional spatial distance maps all distances were calculated between every combination of subdivided segments, i.e. genetic positions, and again averaged for each distance matrix element over 100 to 150 statistically independent chromosome configurations. For two-dimensional interaction maps for each of the subdivided segments all interactions within radii of 30 nm, 40 nm, 50 nm, 60 nm, 70 nm, 80 nm, 90 nm, 100 nm, 110 nm, 120 nm, 130 nm, 140 nm, and 150 nm, respectively where calculated and cumulated into the corresponding interaction matrix, while averaging over 100 to 150 statistically independent chromosome configurations. Notably, the diagonal of the interaction matrix, due to the radii in relation to the segment resolution acts as an intrinsic normalization.

### Scaling Properties of Computer Simulated Genome Architectures

In order to describe quantitatively the different chromatin topologies and microscopic morphologies resulting from the Multi-Loop-Subcompartment (MLS) or the Random-Walk/Giant-Loop (RW/GL) models of simulated chromosomes and simulated nuclei, their scaling behaviour was investigated (Fig. S13-15; and Knoch, 1998; Knoch et al., 1998; Knoch, 2002; Knoch, 2003; Knoch et al., 2009). The scaling behaviour describes how a parameter, e.g. the length of the coast of Britain, depends e.g. on the scale of observation, i.e. the length of the ruler used to measure the coastline. Since the coastline is self-similar it follows a power-law. The exponent of the power-law is the scaling or fractal dimension of the coastline. For Britain it values ∼1.24. Mathematical lines, surfaces and volumes have fractal dimensions of 1.0, 2.0 and 3.0, which agree with their Euklidian dimension. Thus, the British coast scales like a folded and not a straight line. To determine the scaling behaviour of different chromatin topologies the exact spatial distance and exact yard stick dimensions were determined:

The scaling behaviour of the one-dimensional axis of the simulated 30 nm chromatin fibre was investigated by the exact spatial-distance dimension *D_SD_* and the exact yard-stick *D_Y_* dimension. Neglecting the different measuring process *D_SD_* <=> *D_Y_* in the limit (Mandelbrot, 1977). *D_SD_* is the inverse exponent in the scaling relation between the distance *R_SD_*(*C_SD_*) connecting two chain positions and the contour length *C_SD_* e. g. in bp

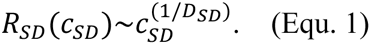

This corresponds to the measurement of spatial distances between genomic markers as function of their genomic separation *C_SD_*. Since *R_SD_* depends for large *C_SD_* on the marker position, the average over several randomly placed marker pairs need to be taken. Genomic separations ≥ 5.2 kbp (the base pair content of one chain segment) up to the whole chromosome were used. All possible maker pairs for genomic separations < 25 Mbp and 5000 pairs > 25 Mbp were taken. The average was taken over 100 to 150 chromosome configurations.

The exact yard-stick dimension *D_Y_* is the negative exponent in the scaling relation between *C_Y_*(*l_Y_*) and the yard-stick length *l_Y_*

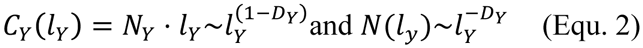

with the number of yard-sticks *N_Y_*, depending itself on *l_Y_*. The beginning coordinates of a randomly chosen chain segment defined the initial start point from which the necessary *N_Y_* to reach both chain ends was calculated by walking along the chain with *l_Y_* sized steps. First the distance to the next segments was determined until it surpassed *l_Y_* (Fig. S13). Hence, the exact chain position and the start point of the next step where the distance equals *l_Y_* is a coordinate point within the previous segment. This is given by the corresponding trigonometric vector equation. Near the chain end and/or for big *l_Y_*, the distance could be > *l_Y_*, thus *N_Y_* was determined by the fraction of *l_Y_* to reach the chain end. The resolution of *l_Y_* was 5, 25 and 50 nm up to scaling length of 10^2^, 10^3^ and 10^4^, respectively. Since, *N_Y_* depends on the initial start point the bigger *l_Y_*, *S*(*l_Y_*) = *abs*(0.05*l_Y_*) (*l_Y_* in nm) start points were averaged, thus the standard error of *D_Y_* was always <0.01. The mean was taken over 100 to 150 statistical chromosome configurations.

### Scaling Analysis Genomic Interactions

The analysis of the scaling properties of the interaction frequency *I*(*s*) as function of the genetic separation *s*maximum deviation function in base pairs is based on the sum and thus average of all possible interactions *k* whose genetic separations between two genetic chromosomal base pair locations *p_i_* and *p_j_* (whith *p_j_* > *p_i_*) are within a window/bin of size *w* centered around s and thus in the interval 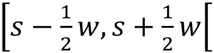 and thus:

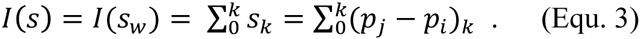

For a fractal self-similar sequence like a random-walk the interaction scaling function *I*(*s*) shows power-law behaviour:

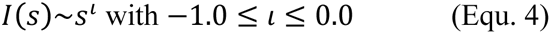

where −1.0 characterizes a negatively, −0.5 a randomly and 0.0 a positively correlated sequence.

To determine the local interaction coefficient ι(*s*) for the analysis of the general behaviour and fine-structural features of long-range interaction scaling as a function of the genetic separation *s*, the following asymmetric finite difference quotient of second order can be applied to 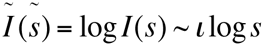 with 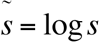:

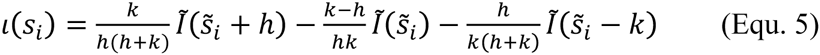

with

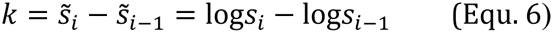

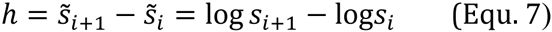

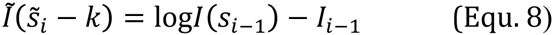

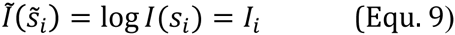

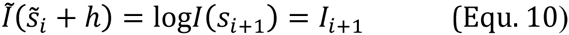

### Correlation Analysis of DNA Sequences of Genomes

The analysis of long-range power-law correlations in genetic sequences here was done as described in earlier (Knoch et al., 2002; Knoch, 2002; Knoch, 2003; Knoch et al., 2009). Briefly: It is based on the concentration profile of single nucleotides along the DNA sequence: The square root of the mean-square deviation between the concentration of nucleotides *c_l_* in a window of length *l* and the concentration 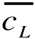 of nucleotides in the entire DNA sequence with length *L* was calculated

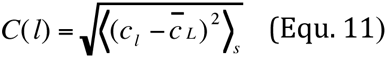

while averaging over all *s = L − l* + 1 possible window positions. Nucleotides used were adenine (A), thymine (T), guanine (G), and cytosine (C) as well as their grouping into purines (A+G) and pyrimidines (T+C). “Unknown” nucleotides were accounted for by using their general appearance probabilities. Since purines/pyrimidines are complimentary, the results are equal and their analysis as base versus non-base equals mapping the DNA sequence to the trajectory of a one-dimensional random-walk. In the following, only the results for purines vs. pyrimidines are considered.

For a fractal self-similar sequence like a random-walk the concentration fluctuation function *C*(*l*) shows power-law behaviour:

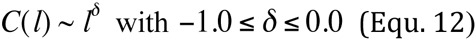

where −1.0 characterizes a negatively, −0.5 a randomly and 0.0 a positively correlated sequence. The power-law behaviour of *C(l)* is connected to the power-law behaviour of the minimum and maximum deviation function *F(l)* ∼ *l*^α^ (Peng et al., 1992), the common autocorrelation function *A(l)* ∼ *l*^γ^, and the power spectrum *S*(*f*) ∼ (1/*f*)^β^ with frequency *f* via

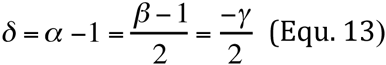

(Prabhu and Claverie, 1992; Chatzidimitriou-Dreismann and Larhammar, 1993; Borovik et al., 1994; Stanley et al., 1994). *C*(*l*) is related to the common autocorrelation function by double summation

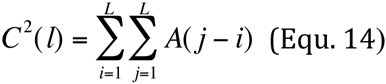

Thus, local random fluctuations are substantially reduced and the analysis leads to a more reliable characterization of the DNA sequence compared to e.g. *A*(*l*) (Peng et al., 1992; Li et al., 1994; Li 1997). Numerical calculation of *C*(*l*) by using Eq. 1 in this sequence of operations

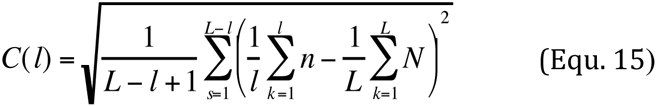

by means of the probabilities for a nucleotide at a certain position *n* = *P*(*s + k*), *N = P*(*k*) and e.g. *P* = 1 for purines and *P* = 0 elsewhere, leads to extreme numerical instabilities (Knoch, 2002; Knoch, 2003; Knoch et al., 2009). These instabilities were avoided by expansion to

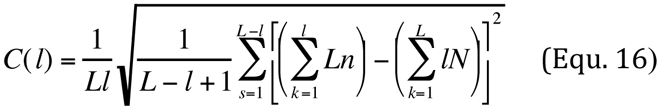

and by exact calculation provided by the GNU multiple precision package GMP. The greater stability is due to the start of deviations from the exact result and becomes especially important for sequences longer than 10^5^ base pairs. To save computer power, the program adjusted automatically the precision (guaranteeing > 8 digits) depending on the sequence length.

To determine the local correlation coefficient *δ*(*l*) for the analysis of the general behaviour and fine-structural features of long-range correlations as a function of the window size *l*, the following asymmetric finite difference quotient of second order was applied to 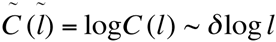 with 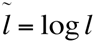:

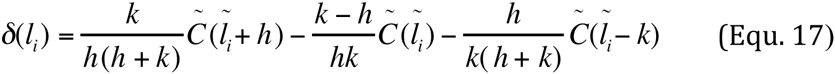

with

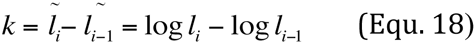

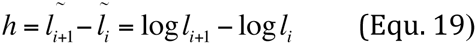

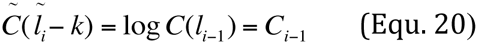

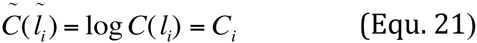

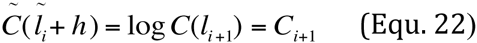

To reduce the enormous computer power needed to calculate *C*(*l*) and *δ*(*l*) for every possible *l*, every *l* from 1 to 10^4^ bp and only 900 logarithmically distributed *l* for every order of magnitude thereafter were chosen. Calculations were performed on our world wide Correlizer@home BOINC grid (<u>http://svahesrv2.bioquant.uni-heidelberg.de/correlizer/</u>), the German D-Grid, the European Grid Initiative EGEE, as well as the Erasmus Computing Grid), the Almere Grid, and all the unnamed computing grids we have access through via these. Therefore the analyses were split into jobs of a few minutes computing a small number of windows each, thus being an extremely efficient ‘gap-filler’ on all these machines.

*<u>Random sequences:</u>* To investigate the error behaviour and to determine the origin of various correlation properties, artificial sequences based on different assumptions about their composition were constructed: They were constructed from a uniform distribution of base pairs using a R250 random number generator based on 16 parallel copies of a linear shift register with a period of 2^250^ – 1 (Kirkpatrick and Stoll, 1981). This is a far greater period compared to the linear congruent generator used normally and thus produces series with no structure resulting from the random number generator. The R250 generator is computationally faster as well (Maier, 1991). The base pair composition was biased by the human base pair distribution (A: 30 %, C: 20 %, G: 20 %, T: 30 %). Other biases were not chosen here, since a simple base pair bias does not result in different general, multi-scaling or fine-structure correlation behaviours (for further details and more complicated random sequences see Knoch et al. 2002; Knoch, 2002; Knoch et al., 2009).

## Supplemental Tables

**STable 1:** Comparison between different chromosome interaction capture methods, showing their different application potential in respect to scientific aims and their signal-to-noise ratio which could function as an intrinsic quality statement (O: one; M: many; A: all; O<->O: oneto-one; P: primer; PCR: polymerase chain reaction; RE: restriction enzyme; Sel.PCR: selection with PCR; Seq: sequencing).

| Method | Interact-<br>ion<br>Types | Interact-<br>ion<br>Selection | 1<sup>st</sup><br>RE | Re-Lig. | 2<sup>nd</sup><br>RE | Ampli-<br>fication | Noise<br>Reduction | Readout | Signal-Noise<br>Ratio |
| --- | --- | --- | --- | --- | --- | --- | --- | --- | --- |
| **3C** | O<->O | P/PCR | Y | Y | N | while<br>sel. PCR | none | gel | + |
| **3C-PCR** | O<->O | P/PCR | Y | Y | N | while<br>sel. PCR | none | qPCR | + |
| **4C** | O/M->A | P/PCR | Y | Y | Y/N | while<br>sel. PCR | none | fluoresc.<br>chip | + |
| **3C/4C-seq** | O/M->A |  | Y | Y | Y | Wwhile<br>sel.PCR<br>and<br>for seq. | none | sequencing | ++ |
| **5C** | M<->M | P<br>com/PCR | Y | Y | Y/N | while<br>sel. PCR | none | sequencing | ++ |
| **Hi-C** | A<->A | none | Y | Y | Y | for<br>seq. | re-ligation<br>bead<br>selection | sequencing | ++ |
| **T2C** | arbitrary<br>combinat<br>ion | oligo<br>capture | Y | Y | Y | for<br>seq | oligo<br>placement | sequencing | +++ |

**STable 2:** The quality and multiplexability of *T2C* is shown by a detailed overview of the regions investigated (grouped) on one capture array, of the homo sapiens (HS) and mus musculus (MM) genomes, with their chromosome and chromosomal position and size, the use of which first and second restriction enzyme or in case of very high resolution sonication, as well as the average fragment size calculated from < *L_Fragment_* > = *L_Region_*/*F_Fragment_*, and the number of oligos per region. The “name” of the region gives the borders with respect to the ideogram bands.

| Capture Array Grouping | Region “Name” | Chromosome [N] | Chromosomal Position | | Region Size $L_{Region}$ [bp] | Restriction Enzymes or Sonication | | Number of Fragments $N_{Fragment}$ [N] | Average Fragment Size $L_{Fragment}$ [bp] | Number of Capturing Oligos $N_{Oligos}$ [N] |
| --- | --- | --- | --- | --- | --- | --- | --- | --- | --- | --- |
|  |  |  | Start [bp] | End [bp] |  | 1st RE | 2nd RE/ Sonication |  |  |  |
| single capture array | 11p 15.4-15.5 | HS 11 | 1,110,650 | 3,216,350 | 2,105,700 | BglIII | NlaIII | 344 | 6121 | 525 |
| single capture array | 7q E3-F1 | MM 7 | 109,876,350 | 111,966,600 | 2,090,250 | HindIII | NlaIII | 719 | 2915 | 800 |
| single capture array | 3q A3-B | MM 3 | 32,548,927 | 36,548,927 | 4,000,000 | ApoI | sonication | 7,079 | 565 | 10,827 |
|  | 3q F1-F2.2 | MM 3 | 92,363,599 | 97,863,599 | 5,500,000 |  |  | 9,045 | 608 | 12,353 |
|  | 4q B1-B3 | MM 4 | 50,792,682 | 57,292,682 | 6,500,000 |  |  | 12,296 | 528 | 18,776 |
|  | 6q C1 | MM 6 | 64,999,977 | 70,999,977 | 6,000,000 |  |  | 13,466 | 445 | 13,737 |
|  | 6q F1-F3 | MM 6 | 119,932,440 | 125,432,440 | 5,500,000 |  |  | 9,292 | 591 | 11,780 |
|  | 7q C | MM 7 | 62,627,388 | 69,127,388 | 6,500,000 |  |  | 14,438 | 450 | 18,746 |
|  | 7q E3-F1 | MM 7 | 106,961,038 | 117,461,038 | 10,500,000 |  |  | 19,405 | 541 | 26,024 |
|  | 10q A2-A4 | MM 10 | 17,344,736 | 23,844,736 | 6,500,000 |  |  | 11,613 | 559 | 17,360 |
|  | 10q B4-B5.3 | MM 10 | 63,148,728 | 69,148,728 | 6,000,000 |  |  | 11,135 | 538 | 17,382 |
|  | 10q D3 | MM 10 | 125,635,645 | 130,135,645 | 4,500,000 |  |  | 7,437 | 605 | 10,773 |
|  | 11q A3.3-A5 | MM 11 | 29,683,511 | 37,183,511 | 7,500,000 |  |  | 11,726 | 639 | 18,387 |
|  | 12q F1-F2 | MM 12 | 110,657,161 | 118,657,161 | 8,000,000 |  |  | 12,108 | 660 | 16,156 |
|  | 14q B-C2 | MM 14 | 43,002,529 | 52,502,529 | 9,500,000 |  |  | 17,216 | 551 | 19,788 |
|  | 16q A2-B1 | MM 16 | 16,061,845 | 22,061,845 | 5,000,000 |  |  | 9,377 | 533 | 13,444 |
|  | 17q B1-B2 | MM 17 | 33,143,007 | 41,143,007 | 8,000,000 |  |  | 15,621 | 512 | 19,308 |
|  |  |  |  |  | total<br>99,500,00 |  |  | total<br>181,254 | average<br>549 | total<br>244,841 |

**STable 3:**
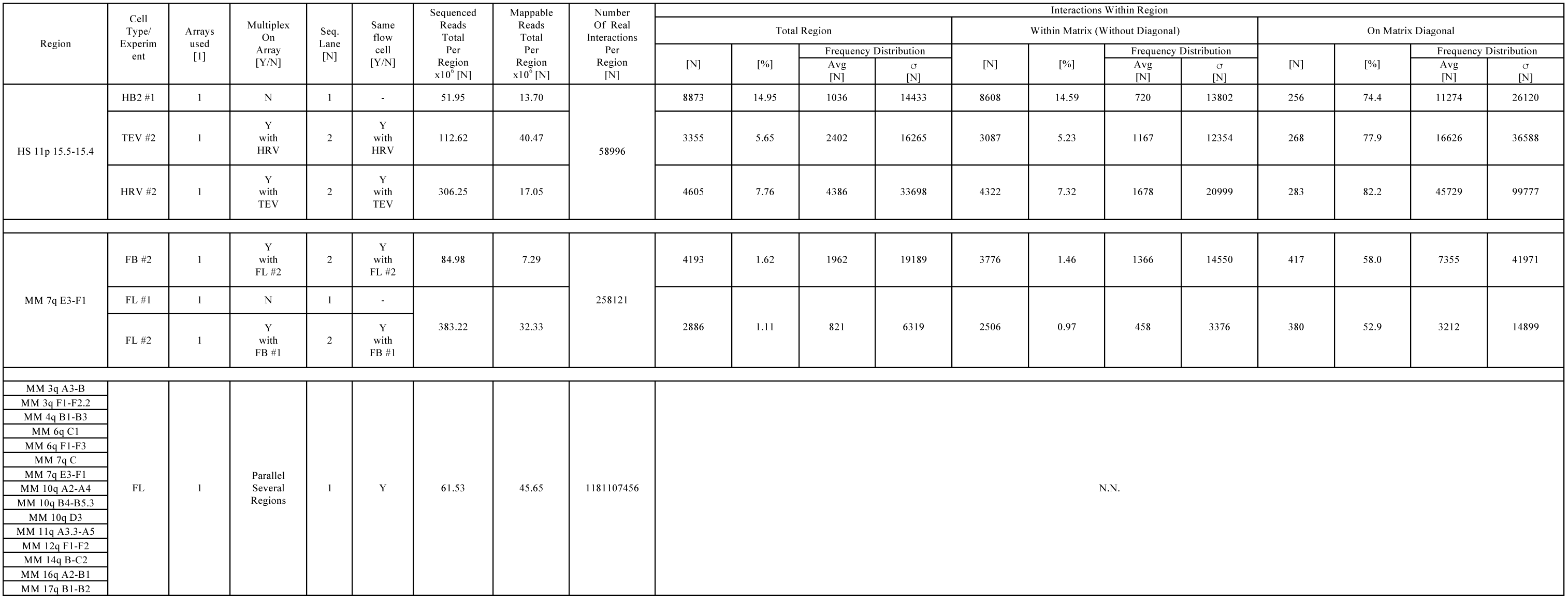
Sequencing and interaction statistics of the experiments done with *T2C* for the regions investigated (grouped) for the Homo sapiens (HS) and Mus musculus (MM) genomes, in respect to the number of capture arrays used, whether and how the multiplexing was done, results in sequenced reads of which sub-fraction could be uniquely mapped, and finally sorted into square interaction matrices (notably, the matrix is mirrored at the diagonal), which can be analysed in total or according to whether the interactions are within the matrix or on the matrix diagonal concerning the number of existent interactions, their fraction of in total possible interactions, and the frequency distribution of the frequency of the interactions. For the high-resolved regions using ApoI as restriction enzyme and sonication as second “restriction” due to the low number of sequence reads in respect to the total number of interactions no analysis concerning the interactions within the region was performed.

| Region | Cell Type/<br>Experiment | Arrays used<br>[1] | Multiplex On Array<br>[Y/N] | Seq. Lane<br>[N] | Same flow cell<br>[Y/N] | Sequenced Reads Total Per Region<br>x10<sup>6</sup> [N] | Mappable Reads Total Per Region<br>x10<sup>6</sup> [N] | Number Of Real Interactions Per Region<br>[N] | Interactions Within Region |  |  |  |  |  |  |  |  |  |  |  |
| --- | --- | --- | --- | --- | --- | --- | --- | --- | --- | --- | --- | --- | --- | --- | --- | --- | --- | --- | --- | --- |
|  |  |  |  |  |  |  |  |  | Total Region |  |  |  | Within Matrix (Without Diagonal) |  |  |  | On Matrix Diagonal |  |  |  |
|  |  |  |  |  |  |  |  |  | [N] | [%] | Frequency Distribution |  | [N] | [%] | Frequency Distribution |  | [N] | [%] | Frequency Distribution |  |
|  |  |  |  |  |  |  |  |  |  |  | Avg [N] | σ [N] |  |  | Avg [N] | σ [N] |  |  | Avg [N] | σ [N] |
| HS 11p 15.5-15.4 | HB2 #1 | 1 | N | 1 | - | 51.95 | 13.70 | 58996 | 8873 | 14.95 | 1036 | 14433 | 8608 | 14.59 | 720 | 13802 | 256 | 74.4 | 11274 | 26120 |
|  | TEV #2 | 1 | Y with HRV | 2 | Y with HRV | 112.62 | 40.47 |  | 3355 | 5.65 | 2402 | 16265 | 3087 | 5.23 | 1167 | 12354 | 268 | 77.9 | 16626 | 36588 |
|  | HRV #2 | 1 | Y with TEV | 2 | Y with TEV | 306.25 | 17.05 |  | 4605 | 7.76 | 4386 | 33698 | 4322 | 7.32 | 1678 | 20999 | 283 | 82.2 | 45729 | 99777 |
| MM 7q E3-F1 | FB #2 | 1 | Y with FL #2 | 2 | Y with FL #2 | 84.98 | 7.29 | 258121 | 4193 | 1.62 | 1962 | 19189 | 3776 | 1.46 | 1366 | 14550 | 417 | 58.0 | 7355 | 41971 |
|  | FL #1 | 1 | N | 1 | - | 383.22 | 32.33 |  | 2886 | 1.11 | 821 | 6319 | 2506 | 0.97 | 458 | 3376 | 380 | 52.9 | 3212 | 14899 |
|  | FL #2 | 1 | Y with FB #1 | 2 | Y with FB #1 |  |  |  |  |  |  |  |  |  |  |  |  |  |  |  |
| MM 3q A3-B | FL | 1 | Parallel Several Regions | 1 | Y | 61.53 | 45.65 | 1181107456 | N.N. |  |  |  |  |  |  |  |  |  |  |  |
| MM 3q F1-F2.2 |  |  |  |  |  |  |  |  |  |  |  |  |  |  |  |  |  |  |  |  |
| MM 4q B1-B3 |  |  |  |  |  |  |  |  |  |  |  |  |  |  |  |  |  |  |  |  |
| MM 6q C1 |  |  |  |  |  |  |  |  |  |  |  |  |  |  |  |  |  |  |  |  |
| MM 6q F1-F3 |  |  |  |  |  |  |  |  |  |  |  |  |  |  |  |  |  |  |  |  |
| MM 7q C |  |  |  |  |  |  |  |  |  |  |  |  |  |  |  |  |  |  |  |  |
| MM 7q E3-F1 |  |  |  |  |  |  |  |  |  |  |  |  |  |  |  |  |  |  |  |  |
| MM 10q A2-A4 |  |  |  |  |  |  |  |  |  |  |  |  |  |  |  |  |  |  |  |  |
| MM 10q B4-B5.3 |  |  |  |  |  |  |  |  |  |  |  |  |  |  |  |  |  |  |  |  |
| MM 10q D3 |  |  |  |  |  |  |  |  |  |  |  |  |  |  |  |  |  |  |  |  |
| MM 11q A3.3-A5 |  |  |  |  |  |  |  |  |  |  |  |  |  |  |  |  |  |  |  |  |
| MM 12q F1-F2 |  |  |  |  |  |  |  |  |  |  |  |  |  |  |  |  |  |  |  |  |
| MM 14q B-C2 |  |  |  |  |  |  |  |  |  |  |  |  |  |  |  |  |  |  |  |  |
| MM 16q A2-B1 |  |  |  |  |  |  |  |  |  |  |  |  |  |  |  |  |  |  |  |  |
| MM 17q B1-B2 |  |  |  |  |  |  |  |  |  |  |  |  |  |  |  |  |  |  |  |  |

**STable 4:** General consensus loop sizes and thus position relative to the start of the first loop at the first loop base determined for human HB2, as well as HEK293T TEV (intact cohesin) and HRV (cleaved cohesin) cells of the IGF/H19 region at HS 11p 15.5-15.4. The subchromosomal domain size is calculated for domains with defined borders only from the sum of the loop sizes present.

| Loop<br>[#] | Loop Size<br>[kbp] | Domain/Linker<br>[#] |
| --- | --- | --- |
| 1 | 15.0 | Domain 1 |
| 2 | 45.9 |  |
| 3 | 56.0 |  |
| 4 | 43.0 |  |
| 5 | 56.7 |  |
| 6 | 48.1 |  |
| 7 | 63.2 | Linker 2 |
| 8 | 45.9 | Domain 2<br>728.5 |
| 9 | 59.6 |  |
| 10 | 51.7 |  |
| 11 | 37.3 |  |
| 12 | 76.1 |  |
| 13 | 28.7 |  |
| 14 | 46.7 |  |
| 15 | 25.1 |  |
| 16 | 76.8 |  |
| 17 | 25.8 |  |
| 18 | 59.6 |  |
| 19 | 13.6 |  |
| 20 | 42.3 |  |
| 21 | 47.4 |  |
| 22 | 51.7 |  |
| 23 | 40.2 |  |
| 24 | 33.7 | Linker 2 |
| 25 | 48.1 | Domain 3<br>403.4 |
| 26 | 39.5 |  |
| 27 | 44.5 |  |
| 28 | 63.2 |  |
| 29 | 46.7 |  |
| 30 | 53.8 |  |
| 31 | 48.8 |  |
| 32 | 25.1 |  |
| 33 | 33.7 |  |
| 34 | 43.1 | Linker 3 |
| 35 | 46.7 | Domain 4 |
| 36 | 73.2 |  |
| 37 | 43.8 |  |
| 38 | 63.2 |  |
| 39 | 62.4 |  |
| Average<br>StdDev<br>StdErr | 48.6±14.5±2.4<br>46.7±15.1±8.7 | Loops<br>Linker |

**STable 5:** General consensus loop sizes and thus position relative to the start of the first loop at the first loop base determined for mouse fetal brain (FB; inactive β-globin) and fetal liver (FL; active β-globin) cells of the β-globin locus at MM 7q E3-F1. The subchromosomal domain size is calculated for domains with defined borders only from the sum of the loop sizes present.

| *Loop*<br>*[#]* | *Loop Size*<br>*[kbp]* | *Domain/Linker*<br>*[#]* |
| --- | --- | --- |
| 1 | 40.6 | Domain 1 |
| 2 | 37.7 |  |
| 3 | 55.6 | Linker 1 |
| 4 | 52.7 | Domain 2<br>1343.6 |
| 5 | 39.9 |  |
| 6 | 84.1 |  |
| 7 | 50.6 |  |
| 8 | 78.4 |  |
| 9 | 47.3 |  |
| 10 | 88.3 |  |
| 11 | 51.3 |  |
| 12 | 53.4 |  |
| 13 | 59.8 |  |
| 14 | 47.4 |  |
| 15 | 47.7 |  |
| 16 | 57.7 |  |
| 17 | 86.9 |  |
| 18 | 53.4 |  |
| 19 | 72.0 |  |
| 20 | 44.9 |  |
| 21 | 54.2 |  |
| 22 | 93.3 |  |
| 23 | 77.7 |  |
| 24 | 52.7 |  |
| 25 | 49.9 |  |
| 26 | 82.7 | Linker 2 |
| 27 | 48.5 | Domain 3 |
| 28 | 37.8 |  |
| 29 | 45.0 |  |
| 30 | 83.4 |  |
| 31 | 42.7 |  |
| 32 | 62.0 |  |
| 33 | 48.5 |  |
| 34 | 60.6 |  |
| Average<br>StdDev<br>StdErr | 57.8±16.2±2.9<br>69.2±19.2±13.6 | Loops<br>Linker |

**STable 6:** General consensus loop sizes and thus position relative to the start of the first loop at the first loop base determined for mouse MEL cells of one part of the IGH locus at MM 12q F1-F2. The subchromosomal domain size is calculated for domains with defined borders only from the sum of the loop sizes present.

| *Loop*<br>*[#]* | *Loop Size*<br>*[kbp]* | *Domain/Linker*<br>*[#] [kbp]* |
| --- | --- | --- |
| 1 | 33.8 | Domain 1<br>333.3 |
| 2 | 47.5 |  |
| 3 | 52.7 |  |
| 4 | 38.7 |  |
| 5 | 22.0 |  |
| 6 | 23.4 |  |
| 7 | 38.6 |  |
| 8 | 38.7 |  |
| 9 | 37.8 |  |
| Average<br>StdDev<br>StdErr | 37.0±9.9±3.3 | Loops |

**STable 7:**
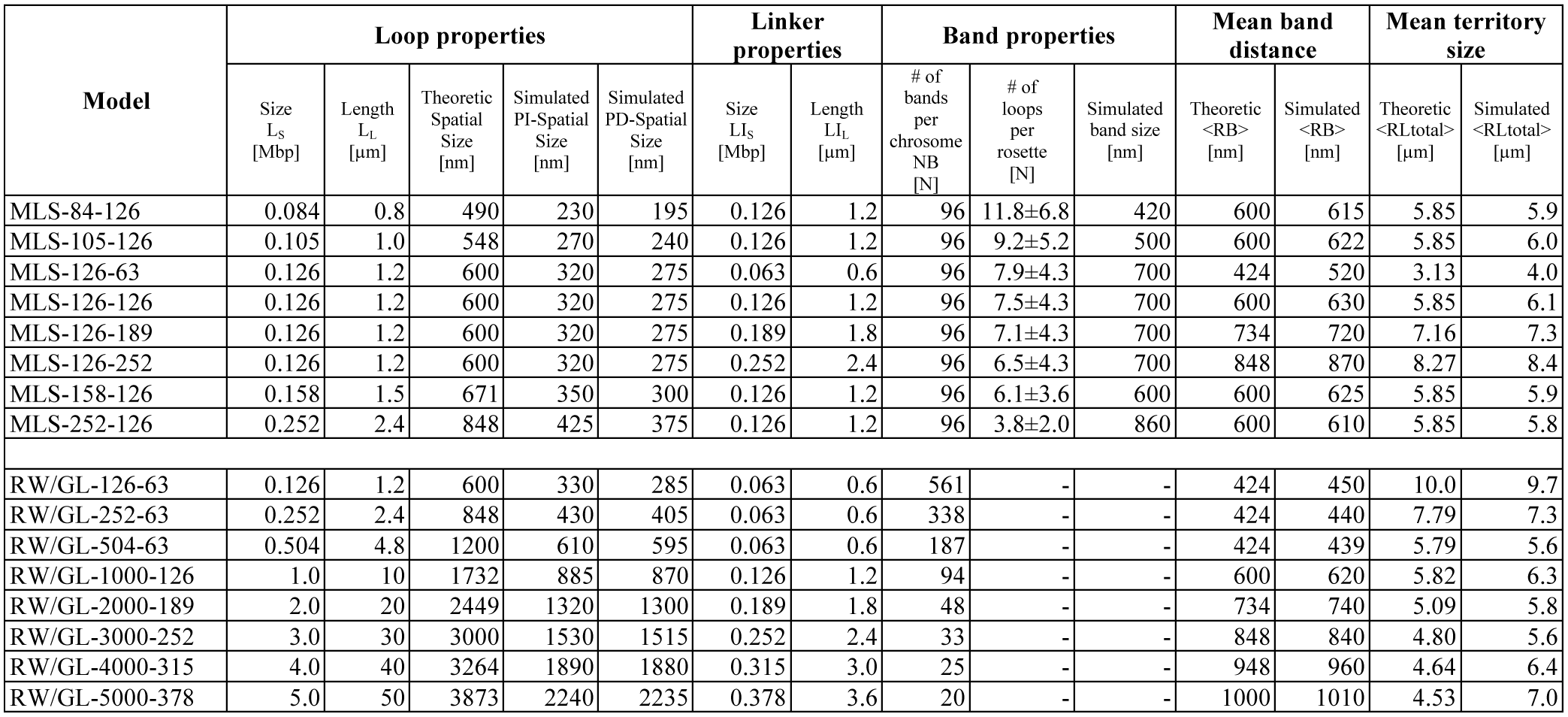
Simulated chromosome models with their physical properties (as in detail described Knoch, 1998; Knoch 2002; Knoch, submitted; Knoch submitted): The band number is the number of subcompartments or loops per chromosome. The average theoretic loop size is 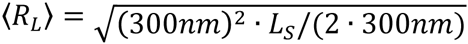 and was determined from simulated position dependent (PD) and independent (PI) spatial distances for genomic separations at half the loop size. The average simulated band size is the average extension of the mass distribution. The average theoretic band distance is 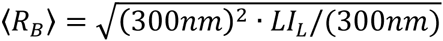 and was simulated from the average spatial distance averages between succeeding subcompartments. The average theoretic territory size is 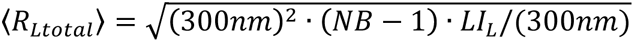 and was simulated from the average mass distribution extension. Naming of models: [model name]-[loop length]-[linker length].

## Supplemental Movies

*SMovie 1:*

Brownian Dynamics simulated decondensation from a metaphase starting configuration of a simulated Multi-Loop-Subcompartment model with 126 kbp loops and linkers with segment length of 50 nm (∼5.2 kb). The whole movie is 750 ms long and shows how abruptly the metaphase chromosome expands explosively due to its high density while opening the linker which is constrained to a loop in metaphase. Nevertheless, the rosettes form distinct chromatin territories in which the loops do not intermingle freely (see also Fig. 10) in contrast to other models such as the RW/GL model. The final shape and form in a whole nucleus would be determined by the limitations the other adjacent chromosomes provide. The difference densities during decondensation also resemble nicely the conditions of shorter linkers, general genome regions with higher densities, or also the variation of nuclear volumes. Notably, the intrinsic movement of the chromatin fibre is clearly taking place on the milli-second scale, and hence, obviously a topological preformed architecture would dissolve within seconds if it would not be stable.

*SMovie 2:*

Brownian Dynamics simulation of the consensus architecture of the of the IGF/H19 region at HS 11p 15.5-15.4 region, with a segment length of 20 nm (∼ 2.0 kbp; colours of loops like in Fig. 1B middle, with additional linkers at the beginning and end of the region in red.). The whole movie encompasses 146 ms and shows the high intrinsic dynamics of the loops and the loop aggregate/rosette. Obviously, the single subchromosomal domains are constrained by the subsequent subchromosomal domains as for the β-globin locus (Mov. S3) and in contrast to the IGH locus (Mov. S4), which are connected directly by the linker. Hence, obviously a topological preformed architecture would dissolve within seconds if it would not be stable. Nevertheless, the loop aggregates/rosettes form distinct subchromosomal domains in which the loops do not intermingle freely (see also Fig. S10) in contrast to other models such as the RW/GL model. The final shape and form in a whole nucleus would be determined by the limitations adjacent chromosomes provide.

*SMovie 3:*

Brownian Dynamics simulation of the consensus architecture of the of the β-globin locus at MM 7q E3-F1 region, with a segment length of 20 nm (∼ 2.0 kbp; colours of loops like in Fig. 1C middle, with additional linkers at the beginning and end of the region in red.). The whole movie encompasses 146 ms and shows the high intrinsic dynamics of the loops and the loop aggregate/rosette. Obviously, the single subchromosomal domains are constrained by the subsequent subchromosomal domains, which are connected directly by the linker. Hence, obviously a topological preformed architecture would dissolve within seconds if it would not be stable. Besides, the constrains resulting from the relatively large loop number of the relatively big subchromosomal domain are also obvious compared to the IGF/H19 region (Mov. S2) or the single subchromosomal domain of the IGH locus. Nevertheless, the loop aggregates/rosettes form distinct subchromosomal domains in which the loops do not intermingle freely (see also Fig. S10) in contrast to other models such as the RW/GL model. The final shape and form in a whole nucleus would be determined by the limitations adjacent chromosomes provide.

*SMovie 4:*

Brownian Dynamics simulation of the consensus architecture of the IGH locus at MM 12q F1-F2 locus with a segment length of 20 nm (∼ 2.0 kbp; colours of loops like in Fig. 1D middle, with an additional linker in red.). The whole movie encompasses 146 ms and shows the high intrinsic dynamics of the loops and the loop aggregate/rosette is including the fact that this single subchromosomal domain can freely rotated since it is now not constrained by other subchromosomal domains compared to the β-globin locus (Mov. S2) or the the IGF/H19 region (Mov. S3). Hence, obviously a topological preformed architecture would dissolve within seconds if it would not be stable. Nevertheless, the loop aggregates/rosettes form distinct subchromosomal domains in which the loops do not intermingle freely (see also Fig. S10) in contrast to other models such as the RW/GL model. The final shape and form in a whole nucleus would be determined by the limitations adjacent chromosomes provide.

## Supplemental Figures

*SFigure 1:*

The high-quality optimization of *T2C* in the first restriction, ligation, and second restriction of the human HB2, HEK293T TEV, and HEK293T HRV samples: A, Agarose gel (0.6% wt/vol) showing the primary enzyme restriction by BglII (six-cutter enzyme) for the H2B samples, which typically produces a smear of DNA fragments between 0.4-12 kbp (two replicates, HB2-1 and HB2-2 are shown). B, Agarose gel (1.5% wt/vol) showing that after ligation for the H2B samples, the DNA smear has returned to a sharp band around 12 kbp (two replicates, HB2-1 and HB2-2 are shown). Ligated samples were loaded undiluted and diluted 1:10. C, Agarose gel (1.5% wt/vol) showing the secondary enzyme restriction by NlaIII (four-cutter enzyme) for the H2B samples, which results in a DNA smear of 0.1-2 kbp (the first replicate HB2-1 was used for the array). D, Agarose gel (0.6% wt/vol) showing the primary enzyme restriction by BglII (six-cutter enzyme) for the TEV and HRV samples, which typically produces a smear of DNA fragments between 0.4-12 kbp. E, Agarose gel (1.5% wt/vol) showing that after ligation for the TEV and HRV samples, the DNA smear has returned to a sharp band around 12 kbp. Ligated samples were loaded undiluted. F, Agarose gel (1.5% wt/vol) showing the secondary enzyme restriction by NlaIII (four-cutter enzyme) for the TEV and HRV samples, which results in a DNA smear of 0.1-2 kbp.

*SFigure 2:*

The high-quality optimization of *T2C* in the first restriction, ligation, and second restriction of the mouse fetal liver (FL) and fetal brain (FB) using HindIII as first and NlaIII as second restriction enzyme, as well as fetal liver using ApoI as first restriction enzyme and sonication: A, Agarose gel (0.6% wt/vol) showing the primary enzyme restriction by HindIII (six-cutter enzyme) for the fetal liver and brain samples, which typically produces a smear of DNA fragments between 0.4-12 kbp (two replicates are shown). B, Agarose gel (1.5% wt/vol) showing that after ligation for the fetal liver and brain samples, the DNA smear has returned to a sharp band around 12 kbp for different amounts of DNA. C, Agarose gel (1.5% wt/vol) showing the secondary enzyme digestion by NlaIII (four-cutter enzyme) for the fetal liver and brain samples, which results in a DNA smear of 0.1-2 kbp. D, Agarose gel (0.6% wt/vol) showing the primary enzyme digestion by ApoI (five-cutter enzyme) for a fetal liver sample, which typically produces a smear of DNA fragments between 0.2-5 kbp. E, Agarose gel (1.5% wt/vol) showing that after ligation for a fetal liver sample, that the DNA smear has returned to a sharp band around 12 kbp. F, Agarose gel (1.5% wt/vol) showing the sonication efficiency of the ligated material for the a fetal liver sample for different amounts of DNA (1-4 µl of DNA).

*SFigure 3:*

The normalized frequency distribution of the fragment sizes for Bgl II, Hind III, and ApoI, shows the high resolution with many fragments being at the limit of what can be captured, i.e. ∼50 bp. Thus, the resolution reached for many fragments even with relatively infrequent restriction enzymes as BglII and HindIII, is near or at the fundamental limits of cross-link techniques (persistence length of free DNA on average ∼50nm or ∼140 bp; typical protein/nucleosome binding region ∼200-500 bp). Additionally the normalized frequency distributions within the region are a very good representation of the general restriction distribution of the enzymes, with minor local variances. In the case of Apo I, due to the many regions and their size, i.e. the high degree of representation (∼1/30 of the entire genome) no difference was found here, so that only the ApoI frequency of the regions is shown.

*SFigure 4:*

Reproducibility, variation and statistical limit in *T2C*: Whereas in the case of the human IGF/H19 11p 15.5-15.4 region in HB2 cell (A; HB2 1.1) only one experiment in one sequencing lane was done, in the HEK293T TEV (B) and HEK293T HRV (C) cell system one capture array was used and two lanes were sequenced (TEV 2.1 & 2.2; HRV 2.1 & 2.2), In the case of the mouse β-globin region 7qE3-F1 for fetal brain (D) and fetal liver (E) cells, one capture array was used and two lanes were sequenced (FB 2.1 & 2.2; FL 2.1 & 2.2) with an additional single capture array and one sequenced lane for fetal liver cells (E; FL 1.1). The matrices clearly show the reproducibility in different sequence lanes (B-E) as well as between different arrays (E) and that in different cell types the basic structural organization is the same (A-C; D-E). The variation between different lanes and arrays is extremely small and shows, that summing up the different lanes only leads to minor additional interactions (compare totals to individuals), thus in principle the statistical limit is reached. Note: the images have a logarithmic frequency range with rainbow coloured visualization and the matrices were normalized for comparability.

*SFigure 5:*

The interaction network of the IGF/H19 region HS 11p 15.5-15.4 resulting from *T2C* shows not only its high quality but also the agreement and differences between different capture techniques: Obviously, *T2C* leads to much more detailed and clearer results (B, C, E) compared to HiC for IMR90 cells (A; data from Dixon et al., 2012), and shows interactions of loops at the domain borders or possibly hint at two domain borders (B), suggesting that either i) neighbouring loops interact at domain borders, i.e. there are two larger interaction domains whose borders interact with the entire other interaction domain, ii) there are two sets of interaction domains with a different border either due to the two alleles always present or any other subpopulation of cells, or iii) there is one large interaction domain followed by a very small interaction domain, which is followed again by a larger interaction domain. On the fragment level *T2C* shows not only a clearer dedicated pattern of complex interaction networks (B), but also detailed visualization of *T2C* interactions from one viewpoint (linear frequency range: C; logarithmic frequency range: E) shows these interaction networks not only in much more detail and clearer compared to 4C-seq (D, F) but also identifies more novel, i.e. previously unknown interactions. Also immediately the advantage of a logarithmic frequency range with a rainbow coloured visualization becomes clear as well as the fact that a squared matrix representation is much easier to understand in terms of relating interactions either with structural or annotational features due to the perception of the human visual cortex trained to horizontal and vertical analysis.

*SFigure 6:*

*T2C* interaction network of the β-globin region MM 7qE3-F1: Again *T2C* leads to interaction matrices with high-resolution, a high-frequency range, and unseen quality (fetal brain: A; fetal liver: B; again the matrices are normalized to each other for comparability), with all the known interactions, as e.g. between the β-globin promoter and the local control region (LCR) and between the LCR-3’HS1 sites, and the increased interaction degree of the active β-globin gene. These *T2C* data can be further annotated by other data (bottom), e.g. restriction enzyme sites, transcription factor binding sites, histone modifications and other data, where again the high-resolution and quality of *T2C* will allow for the first time to make sound statements. And again immediately the advantage of a logarithmic frequency range with a rainbow coloured visualization becomes clear as well as the fact that a squared matrix representation is much easier to understand in terms of relating interactions either with structural or annotational eatures due to the perception of the human visual cortex trained to horizontal and vertical analysis.

*SFigure 7:*

The similarity of *T2C* combined maximum/average projections of the IGF/H19 region HS 11p 15.5-15.4 HB2, HEK293T TEV (intact cohesin), and HEK293T HRV (cleaved cohesin), cell types and functional states shows that i) clearly dedicated architectural loops exist, whose ii) locations show only minor differences between the cell type or functional state, despite varying interaction frequencies whose origin (e.g. due to different crosslink-influencing binding of proteins, which might not have even a structural relevance, but influence the experimental crosslinking) and functional relevance, e.g. in terms of loop formation e.g. due to an enhancer-gene looping interaction remain still unclear. Consequently, the genome has a clear basic structural consensus architecture between different cell types or states, which is functionally altered or fine-tuned. The combination also shows, in comparison with the interaction matrices (Fig. 1B-D; Fig. S4-6), that construction of an automatic loop detection algorithm is highly dependent on local conditions, loop architecture and inter- and intra-loop interactions as thus presumably also on the local chromatin quasi-fibre compaction and architecture – all including their dynamics and functional alterations/dependencies. Thus, a one-for-all algorithm with a one-for-all parameter set to detect the loops might not exist for a genome wide analysis.

*SFigure 8:*

Experimental interaction scaling curves derived from *T2C* for the human IGF/H19 11p 15.5-15.4 region in HB2, HEK293T TEV, and HEK293T HRV (A-C) cell systems, as well as the mouse β-globin locus 7qE3-F1 for fetal brain and liver cells using a 3 bp average of the raw data from 1 to 200 bp (red) and thereafter in the polymer regime a combination of a grouping with a 1% resolution per order of magnitude and an added running window smoothing average to get more general characteristics for raw (blue) as well as “secured” (i.e. only 100% non-neighbouring or restricted and re-ligated interactions are used; light blue). The values < 10 bp are in principle due to the algorithm used and for transparency not discarded since they nevertheless show the extrapolation from existing values > 10 bp. In all cases the interactions show long-rang scaling with multi-scaling on different scales as well as a fine structure on top (Fig. 2A). From 1-195 bp the behaviour and fine structure is associated with the nucleosomal winding of the DNA around the nucleosome, i.e. e.g. the 1.7 and 145 bp (the peak of interactions) winding of the DNA double helix around nucleosomal core and the nucleosomal repeat length of 195 bp (Fig. 2D; Fig. S9) and its dedicated fine structure as seen also in the DNA sequence correlation multi-scaling power-law correlations with its fine structure (Fig. 1E; Fig. S16-21). This is despite the used resolution of the here used restriction enzymes visible in the unsecured data and also a dependent on the used restriction enzyme. Thereafter, i.e. for scales > 195 bp there is a clear plateau like increase up to a “peak” at ∼10^3^ bp, which is clearly indicating that the nucleosomes interact equally or even more up to a scale of ∼10^3^ bp, i.e. that on average a chromatin quasi fibre is formed with 5±1 nucleosomes per 11 nm, since a nucleosome has a dimension on that scale and we see a clear decrease of interactions after a scale of 6 nucleosomes, i.e. ∼1.0-1.2 bp. Whether the fluctuations within this regime can be associated with the order of nucleosomes within the fibre conformation is yet hard to say. In the case of “secured” interactions the plateau and increase to the peak are seen, thus this is a real effect and not e.g. due to neighbouring or unrestricted DNA fragments. Thereafter, there is a multi-scaling behaviour with three regimes: i) the first regime until ∼10^4^ bp still shows the chromatin quasi fibre formation before it goes over into the ii) regime of chromatin fibre and chromatin loops with a slightly higher decrease in interactions, which then transits iii) to a nearly flat plateau indicating the formation of an aggregated state of rosettes in agreement with polymer simulations (Fig. 2B; Fig. S10-15). Thereafter, we see a sharp decrease due to the limits of the regions for which the experiments were done, i.e. ∼2.1 Mbp. This entire multi-scaling behaviour is not only in agreement with polymer models (Fig. 2B; Fig. S10-15), but furthermore there is a fine structure which can be associated with the loop sizes already determined in the interaction matrices (of course, the loops are not as clearly visible as in the simulations due to the variation of the loop sizes, but clear peaks around the average experimental loop sizes is clearly visible). Even beyond that, the entire behaviour is also in agreement with the fine-structured multi-scaling long-range power-law correlations of the DNA sequence itself (Fig. 2E; Fig. S16-21).

*SFigure 9:*

Experimental interaction scaling curves derived from *T2C* for the high-resolution data derived from mouse MEL cells for 15 loci covering in total ∼99 Mbp with a subnucleosomal fragment resolution (Tab. S2) show clearly a fine-structure associated with the nucleosome as shown for the average over all chromosomes (A-C): In general there is an increase of the interaction until a plateau is reached from ∼50 bp to ∼100 bp (A, B), thereafter there is a sharp peak which is ∼1.5 orders of magnitude higher from ∼110 to 195 bp with a width of ∼85 bp, followed by a slight decent to a second “dip/plateau” from ∼230 bp up to the transit to a new descent at ∼10^3^ bp which then obviously transits at ∼10^4^ bp to the known multi-scaling behaviour seen in the lower resolved human and mouse cases (Fig. S8). On this behaviour there is a fine structure (A-C) which can be associated with the binding of the DNA double helix to the nucleosomal core (Fig. S8): in the first plateau many peaks are agreeing with those (Fig. S21) of the fine structure found in correlations of the DNA sequence itself, on top of the peak a clear fine structure at 145 bp can be found and again many of the features agree with correlations of the DNA sequence itself (Fig. S21; and Knoch, 2002; Knoch, 2003; Knoch et al., 2009). Additionally, the plateau from 195 bp to 1200 shows also characteristic features, e.g. at 290 bp as well as at 385 bp the peaks are exactly where di-nucleosomal features are expected. This is logical since neighbouring nucleosomes might see each other most likely/often. Astonishingly, the plateau itself only decreases by ∼10%, i.e. nucleosomes 4-6 see the first nucleosome with nearly the exact same probability, which suggests an average quasi fibre, with a packing density of on average 5±1 nucleosomes per 11nm, since the nearest proximity of the 4-6th or any other subsequent nucleosome cannot be smaller than the nucleosome core itself. Since there is a slight dependency in respect to the used (Fig. S8) restriction enzyme with lower resolution, there might be also dependencies here, which might, however, be much smaller due to the high resolution achieved here.

*SFigure 10:*

Simulated chromatin models description and relation/evaluation of spatial distances between genomic markers, in the Immunoglobulin Heavy Chain locus and the Prader-Willi/Angelmann Syndrome region (Knoch, 1998; Knoch et al., 1998; Knoch, 2002; Knoch, 2003; Jhunjhunwhala et. al., 2008; Rauch et al., 2008): A, Volume rendered images of simulated Random-Walk/Giant-Loop and Multi-Loop-Subcompartment models. As a starting conformation with the form and size of a metaphase chromosome (top), rosettes were stacked (α). From such a starting configuration, interphase chromosomes in thermodynamic equilibrium, were decondensed by Monte-Carlo and relaxing Brownian Dynamics steps. A volume rendered image of the simulated Random-Walk/Giant-Loop model containing large loops (5 Mbp) is shown (left; β). Note that the large loops do not form distinct structures but intermingle freely (left; β). In contrast, in a volume rendered image of the simulated Multi-Loop-Subcompartment model, containing 126 kbp sized loops and linkers, the rosettes form distinct chromatin territories in which the loops do not intermingle freely (middle; γ). In an image of the simulated RW/GL model containing 126 kbp loops and 63 kbp linkers, again distinct chromatin territories are formed but in contrast to the MLS model no subcompartments form (right; δ). B, Strategy for position-dependent and position-independent virtual spatial distance measurements in the simulations: For position-dependent virtual distance measurements, the first marker was placed close to the base of the loop (marker 1). The virtual spatial distances were measured from this “viewpoint” to other makers on the chromatin fibre, e.g. in the rosette (1-7) and to a linker (8-10). For position-independent measurements a set of markers separated by the same genomic distance were randomly positioned (x, y,z). C, Comparison between simulated position-dependent (dotted lines) and position-independent (solid lines) spatial distances. The curves (A-D) indicate simulated MLS models with 126 kbp loops and different linker sizes. RW/GL is shown for comparison (A). Position-dependent distances (dotted lines) show a stepwise increase in the region where a linker is connecting two chromatin sub-compartments, while position-independent distances (solid lines) do not show the stepwise increase in spatial distances as a function of genomic separation. D, Random-Walk Giant Loop and Multi-Loop-Subcompartment Models: a indicates the RW/GL model in which large loops are attached to a non-DNA backbone. b shows the simulated model containing a chromatin linker between loops. MLS model is shown containing 126 kbp loops and linkers with individual rosettes spanning 1-2 Mbp. E, The simulated spatial distances as a function of genomic separation are shown for a fixed loop structure. The simulated loop size was 126 kbp. Two virtual genomic markers were chosen that were separated by 252 kbp. The coloured spatial distance map indicates the frequency distribution of simulated spatial distances. F and G, Comparison between experimental data and computer simulated data obtained from spatial distance measurements in the Igh locus (Jhunjhunwhala et. al., 2008) and the Prader-Willi/Angelmann Syndrome region (Rauch et al., 2008) as a function of genomic separation (resolution of the simulation model is 5.2 kbp, i.e. the base-pair size of the polymer segments from which the simulations are setup). Nomenclature is loop size [kbp]-linker size [kbp]-topology. Experimental spatial distance measurements [mm] were plotted as a function of genomic separation [Mbp] in the Igh locus for pre-pro-B cells (blue dots and green circles) and pro-B cells (red squares and pink triangles), and in the Prader-Willi/Angelmann Syndrome region for fibroblasts for either structure preserving para-formaldehyde fixation (FAA; black full circles and triangles) or structure destroying methanol acetic acid fixation (MAA; black open circles and triangles) using λ-probes (black full circle and triangles) and BAC-probes (black open circle and triangles). The comparison shows clearly, best agreement for a multi-loop aggregate/rosette like model, and even clearly structure destroying MAA fixation (see the λ-probes data for very low genomic separation), even for genomic separations at 650 kbp can only anticipate RW-GL models with loops smaller than ∼500 kbp.

*SFigure 11:*

Simulated Multi-Loop-Subcompartment model averaged spatial distance maps with exact spatial distance 〈*R_S_*〉 (first column left) and on the diagonal normalized interaction maps (all other columns) for interaction radii 〈*d_i_*〉 of 30 nm, 50 nm, 70 nm, 90 nm, 110 nm, 130 nm, and 150 nm, for different model parameters ([model name]-[loop length]-[linker length]; Knoch, 1998; Knoch et al., 1998; Knoch, 2002; Knoch, 2003; Jhunjhunwhala et. al., 2008; Rauch et al., 2008; Knoch, submitted; Knoch, submitted) with a resolution of ∼520 bp. Visual inspection immediately reveals on a large scale clearly the formation of distinct subchromosomal domains with a clear edge and inter domain interactions, as well as on intermediate scales the loop and rosette like structure of the MLS model in agreement with the experiment. Again the low overlap of chromosome territories and subchromosomal domains can be seen as in general one of intrinsic MLS model property (Knoch, 1998; Knoch et al., 1998; Knoch, 2002; Knoch, 2003). Thus, already all the effects seen in experimental interaction maps are in agreement with the simulations, and additionally the interactions are a function of all model parameters even in slight details considering that no nucleosomes where modelled here: i) in general the interaction degree depends on the interaction and crosslink probability, ii) the domain size, domain separation, and spacing of loops are proportional to their size (A-H), iii) the interactions between the domains depend on the linker size, the size and number of loops, i.e. density of the rosettes (A-H). Thus, the subtle combination of density of rosettes due to loop size, loop number, chromatin fibre persistence and the thus resulting exclusion effects leading eventually for high numbers to spread out and shielding effects of rosettes, as well as the subtle influence on the interaction pattern between entire domains. The linker between domains and its proportionality to inter-domain interactions is as clearly visible as well as non-equilibration effects, which we deliberately show here to create an understanding of the interactions of loops at aggregate/rosette borders and similar effects. The in general large emptiness of experimental interaction maps depends on the interaction radii and thus also interaction and cross-link frequency. Since the simulations have no nucleosomal resolution, but instead a preset homogeneous chromatin fibre compaction and thus density it becomes also clear that a random-walk of nucleosomes cannot generate such a pattern as the distinct loops would not be smeared out and at least be not as clear (Fig. 1; Fig. S4-7). Thus, this also proves that the experimental crosslink probability, radius, and frequency can be estimated to be relatively low although since the relation contains a too complex parameter set not unambiguously fittable. Furthermore, by zooming in, one can see clearly the loop base structure within a rosette and the pattern created there in agreement with experimental findings at highest resolution.

*SFigure 12:*

Simulated Random-Walk/Giant-Loop model averaged spatial distance maps with exact spatial distance 〈*R_S_*〉 (first column left) and on the diagonal normalized interaction maps (all other columns) for interaction radii 〈*d_i_*〉 of 30 nm, 50 nm, 70 nm, 90 nm, 110 nm, 130 nm, and 150 nm, for different model parameters ([model name]-[loop length]-[linker length]; Knoch, 1998; Knoch et al., 1998; Knoch, 2002; Knoch, 2003; Jhunjhunwhala et. al., 2008; Rauch et al., 2008; Knoch, submitted; Knoch, submitted) with a resolution of ∼520 bp. In the RW/GL model the large loops of several mega base pairs do not form distinct structures but intermingle freely in contrast to the Multi-Loop-Subcompartment model and in disagreement with the experiments (Fig. 1; Fig. S4-7). This is even the case for 126 kbp loops and 63 kbp linkers although there distinct low-overlapping chromatin territories are formed. Again the general properties are found: i) in general the interaction degree depends on the interaction and crosslink probability, ii) the spacing of loops are proportional to their size (A-H), iii) the interactions between loops depend on the loop size, linker size, and the number of loops in proximity (A-H). Subtle combinations of loop size, linker size and thus resulting exclusion effects leading eventually for high numbers to spread out and shielding effects of loops, as well as the subtle influence on the interaction pattern between entire loops appears or increases with at loop sizes smaller than 508 kbp. The linker between loops and its proportionality to inter-loop interactions is as clearly visible as well as non equilibration effects, which we deliberately show here to create an understanding of the interactions of loops and similar effects. The in general large emptiness of experimental interaction maps depends on the interaction radii and thus also interaction and cross-link frequency. The general influence of homogenizing random-walk topologies becomes also obvious and again although due to the non-nucleosomal resolution, but instead the preset homogeneous chromatin fibre compaction and thus density it becomes also clear that a random-walk of nucleosomes cannot generate such a pattern as originating from the loops. Again the simulations prove that the experimental crosslink probability, radius, and frequency can be estimated to be relatively low although since the relation contains a too complex parameter set not unambiguously fittable. Besides, by zooming in, one can see clearly that the loop base structure of large loops is not as defined as within a rosette, since the interaction probability in a loop aggregate/rosette is reduced and thus more defined (Fig. S11)

*SFigure 13:*

Description of the measurement process (A), and spatial-distance *D_SD_* and yard-stick dimensions *D_Y_* of simulated single chromosomes (B-E). The spatial-distance dimension was determined from position independent spatial distances as function of randomly positioned genetic markers with a genomic/curvature separation *c*_SD_. Thus, markers could reside both on the same loop (**a-b**), on different loops (**a-c**), on a loop and in a linker (**b-d**), both in the linker (**d-e**) or on loops belonging to different rosettes (**b-f**). The exact yard-stick dimension *D_Y_* was calculated by walking along the fibre using a yard-stick *l_Y_*. Thus, the start and end of a small *l_Y_* mostly reside in the same loop (**1-16**) in contrast to large *l_Y_* often lying in different loops (**1-6**) or rosettes (**1-3**). The end-point of *l_Y_* was determined exactly by finding the chain segment, where *L*_1_ < *l_Y_* < *L*_2_, before solving the corresponding vector equation. The spatial-distance function *R_SD_* (*c_SD_*) (B, C) and exact yard-stick curve length function *C_SD_* (*l_SD_*) (D, E) shows power-law behaviour as expected for fractal self-similar polymer foldings. The slopes are the spatial-distance dimension *D_SD_* and the exact yard-stick dimension *D_Y_*. The finite size of chromosomes generates a cut-off > ∼80 Mbp or > ∼8 μm after which the power-law behaviour breaks down. Additionally, non-trivial power law behaviour due to the deviation of *D_SD_* and *D_Y_* from 1.0 (a stiff linear segment) or ∼2.0 (a random-walk), four major scaling regions exist. The detailed dimension behaviour is given by the local dimensions *D_SD_*(*C_SD_*) and *D_Y_*(*l_y_*) (C, E) with fluctuations the bigger the closer to the cut-off. The general multi-scaling behaviour of *D_SD_* and *D_Y_* is characterized by an increase from an initial 1.0 for small *c_SD_* and *l_Y_* and characterizing the stiff chromatin segments, over values ∼2.0 as for the random-walk of the segments to a maximum of 3.0 stating the ring-shaped loops of both the MLS and the RW/GL model and globular state of the rosettes of the MLS models according to the *c_SD_* and *l_Y_*. In the MLS model thereafter again local dimensions ∼2.0 are reached describing the random organization of the rosettes relative to each other. Within the general behaviour a fine structure attributable to the loops aggregated in rosettes is present for MLS models, better measured with *D_SD_*. The maximum position and height is proportional to the loop and linker size in the MLS and the RW/GL model, and all the inherent model properties are visible. Naming of models: [model name]-[loop length]-[linker length].

*SFigure 14:*

Simulated Multi-Loop-Subcompartment interaction scaling curves *I*(*s*) shown here for clearness only for interaction radii 〈*d_i_*〉 of 30 nm, 70 nm, 110 nm, and 150 nm for different model parameters ([model name]-[loop length]-[linker length]; Knoch, 1998; Knoch et al., 1998; Knoch, 2002; Knoch, 2003; Jhunjhunwhala et. al., 2008; Rauch et al., 2008; Knoch, submitted; Knoch, submitted) with a resolution of ∼520 bp. The slopes would be the interaction coefficient ι(*s*). Due to the high resolution the general behaviour and fine-structural features of long-range interaction scaling as a function of the genetic separation *s* becomes, however, immediately clear in contrast to the spatial-distance function *R_SD_*(*c_SD_*) (B, C) and exact yard-stick curve length function *C_SD_*(*l_SD_*) (Fig. S13). *I*(*s*) shows long-range power-law behaviour as expected for fractal self-similar polymer foldings. The finite size of chromosomes generates a cut-off > ∼1 to ∼10 Mbp depending on the interaction radius after which the power-law behaviour breaks down. The detailed power-law behaviour is characterized by different behaviours on different scales attributable to the rosette like subcompartments in the MLS model and the scale above where the arrangement of subcompartments into the chromosome territory by a random linker walk is visible. Within the general behaviour the fine structure attributable to the loops aggregated in rosettes is clearly visible in detail for MLS models. The pronouncedness of the fine structure is averaged out by higher interaction radii, and thus shows also what happens if experimental fragment sizes are in- or decreased. Thus, already all the effects seen in simulated interaction maps are here again a function of all model parameters even in slight details considering that no nucleosomes where modelled here: i) in general the scaling degree depends on the interaction and crosslink probability, ii) the domain size, domain separation, and spacing of loops are proportional to their size, iii) the interactions between the domains depend on the linker size, the size and number of loops, i.e. density of the rosettes.

*SFigure 15:*

Simulated Random-Walk/Giant-Loop interaction scaling curves *I*(*s*) shown here for clearness only for interaction radii 〈*d_i_*〉 of 30 nm, 70 nm, 110 nm, and 150 nm for different model parameters ([model name]-[loop length]-[linker length]; Knoch, 1998; Knoch et al., 1998; Knoch, 2002; Knoch, 2003; Jhunjhunwhala et. al., 2008; Rauch et al., 2008; Knoch, submitted; Knoch, submitted) with a resolution of ∼520 bp. The slopes would be the interaction coefficient ι(*s*). Due to the high resolution the general behaviour and fine-structural features of long-range interaction scaling as a function of the genetic separation *s* becomes, however, immediately clear in contrast to the spatial-distance function *R_SD_*(*c_SD_*) (B, C) and exact yard-stick curve length function *C_SD_*(*l_SD_*) (Fig. S13). *I*(*s*) shows long-range power-law behaviour as expected for fractal self-similar polymer foldings, with mainly one general behaviour due to the arrangement of loops into the chromosome territory by a random linker walk. The finite size of chromosomes generates a cut-off > ∼1 to ∼ 10 Mbp depending on the interaction radius after which the power-law behaviour breaks down. Within the general behaviour the fine structure attributable to the loops which are clearly not clustered in rosettes is clearly visible in detail for RWGL models. Thus, again already all the effects seen in simulated interaction maps are here again a function of all model parameters even in slight details considering that no nucleosomes where modelled here: i) in general the scaling degree depends on the interaction and crosslink probability, ii) the domain size, domain separation, and spacing of loops are proportional to their size, iii) the interactions between the domains depend on the linker size, the size and number of loops, i.e. density of the rosettes.

*SFigure 16:*

Introduction to the correlation function *C*(*l*) and the correlation coefficient δ(*l*) for the averages over all chromosomes for each “strain-specific” sequencing of *Homo sapiens, Homo sapiens GRCH37, Mus musculus*, and *Mus musculus C57BL6j* (from the http://www.ebi.ac.uk/genomes/eukaryota.html genome list; Knoch, 1998; Knoch et al., 1998; Knoch et al., 1998; Knoch et al., 2002; Knoch, 2002; Knoch, 2003): A: The correlation function *C*(*l*) of random sequences shows power-law behaviour as expected for a fractal self-similar sequence. All results are numerically exact. The slope is the correlation coefficient δ(*l*) whose value in the linear region is −0.5 (yellow line), which resembles the theoretic value and thus indicating random correlations. The finite sequence length generates a cut-off after which the power-law behaviour breaks down, thus concatenation of two sequences creates a double cut-off. Sequences of *Homo sapiens, Homo sapiens GRCH37, Mus musculus*, and *Mus musculus C57BL6j* exhibit even after averaging over all chromosomes for each strain not only a positively correlated power law behaviour due to a δ(*l*) bigger than - 0.5 (B), but also four regions (numbers 1-4) with different degree of correlation for 10^8^ bp. B: The detailed correlation behaviour is given by the local correlation coefficient δ(*l*), which fluctuates around −0.5 for random sequences. The fluctuations become bigger as the window size approaches the cut-off. *Homo sapiens, Homo sapiens GRCH37, Mus musculus*, and *Mus musculus C57BL6j* again for the averages over all chromosomes per strain, reveal a distinct positively correlated pattern with less fluctuations. In general, δ(*l*) increases from a starting value until a plateaued maximum, before a decrease and a second statistically significant maximum. Finally, δ(*l*) decreases to values characteristic for random sequences and enters the region of fluctuation. Within this general behaviour, a distinct fine-structure is visible more dominantly in the human compared to the mouse case, which survives averaging over different locations within the chromosomal sequences and even over all chromosomes of each strain. The very pronounced local maximum at 11 bp is related to the double helical pitch, whereas the local minima and maxima are related to the nucleosome, which is obvious for 146 bp, but less obvious for other positions e.g. at 172 bp, 205 bp, 228 bp and 248 bp (Fig. S21). The second maximum around 10^5^ is related to chromatin loops of the three-dimensional genome organization and its grouping in aggregates/rosettes. Thus, the 4 regions in *C(l)* (A) can be associated with i) the nucleosome, ii) the compaction of the nucleosome chain into a compacted fibre, iii) the formation of loops of the chromatin fibre, and iv) the arrangement of sub-chromosomal domains in the entire chromosomes. The differences between mouse and human are mainly, an earlier ascent in the case of mouse and a lower plateau in comparison to human sequences (A, B). The differences between the strains within one species are mainly due to differences in the quality of sequences, e.g. unfinished/partial sequencing of the Y-chromosome, and also, but to a less degree, due to real sequence differences as genome rearrangements. C, D: To distinguish real from statistical correlations, the standard deviation was computed from 20 random sequences with similar base pair distribution as in *Homo sapiens* for *C(l)* (c) and δ(*l*) (D). The standard deviation of δ(*l*) shifts only to higher window sizes depending on the sequence length. E, F: For the real sequences the standard deviation for C(*l*) (E, in absolute (thin lines) and relative (thick lines) terms, i.e. dividing the standard deviation *StDev C*(*l*) by the average over all chromosomes of a strain < *C*(*l*) > according to *StDev C*(*l*)_relative_ = *StDev C*(*l*)/< *C*(*l*) >), and δ(*l*) (f) does not increase so much with growing window sizes as in the case of random sequences due to the fact that real genomes have never an entire random sequence organization due to their evolutionary construction.

*SFigure 17:*

A-H: General scaling behaviour of the correlation function *C*(*l*) in *Homo sapiens* and *Homo sapiens GRCH37* strains shows clear power-law behaviour with four clearly distinct regions (Fig. 1S3) of different correlation degree, while approaching the finite sequence length generates a cut-off after which the power-law behaviour breaks down. In most cases differences between chromosomes is bigger than between strains, despite the obvious differences in length of several chromosomes (e.g. *Homo sapiens GRCH37* Y-chromosome) mainly due to differences in the quality of sequences, e.g. unfinished/partial sequencing of the Y-chromosome, and also, but to a less degree, due to real sequence differences as genome rearrangements. Apparently, the differences grow with growing window size *l* (and thus the scale), but appear mainly for *l* >10^65^-10^7^ bp due to approaching the upper cut-off, with the exception of chromosomes 22, X, and Y, where the differences appearing at *l* already >10^3^ bp pointing to a general bigger difference in the general sequence organization in respect to the other chromosomes. This special behaviour as well as the general scaling behaviour is nearly identical with the two mouse strains *Mus musculus*, and *Mus musculus C57BL6j*, thus this is a general feature of those chromosomes across species.

*SFigure 18:*

Detailed multi-scaling behaviour of the correlation coefficient δ(*l*) and its fine-structural features for *Homo sapiens* and *Homo sapiens GRCH37:* The correlation coefficient δ(*l*) shows strong positive correlations for human chromosomes (A-H). In general, δ(*l*) increases from a starting value until a plateaued maximum from 10^2^-10^36^ bp, before a decrease and a second statistically significant maximum at ∼10^5^ bp for all chromosomes of both strains. Finally, δ(*l*) decreases to values characteristic for random sequences and enters the region of fluctuation. The differences between the strains within one species are mainly due to differences in the quality of sequences, e.g. unfinished/partial sequencing of the Y-chromosome, and also, but to a less degree, due to real sequence differences as genome rearrangements. Within this general behaviour, a distinct fine-structure visible more dominantly in the human compared to the mouse case is present in all chromosomes and survives averaging (Fig. S16B, S21). The very pronounced local maximum at 11 bp is related to the double helical pitch, whereas the local minima and maxima are related to the nucleosome, which is obvious for 146 bp, but less obvious for other positions e.g. at 172 bp, 205 bp, 228 bp and 248 bp. The second maximum around 10^5^, shows also a fine-structure which is due to the individual chromatin loops of the three-dimensional genome organization and their grouping in aggregates/rosettes.

*SFigure 19:*

A-G, General scaling behaviour of the correlation function *C*(*l*) in *Mus musculus* and *Mus musculus C57BL6j* shows clear power-law behaviour with four clearly distinct regions (Fig. S16) of different correlation degree, while approaching the finite sequence length generates a cut-off after which the power-law behaviour breaks down. In most cases differences between chromosomes is bigger than between strains, despite the obvious differences in length of several chromosomes (e.g. *Mus musculus C57BL6j* Y-chromosome) mainly due to differences in the quality of sequences, e.g. unfinished/partial sequencing of the Y-chromosome, and also, but to a less degree, due to real sequence differences as genome rearrangements. Apparently, the differences grow with growing window size *l* (and thus the scale), but appear mainly for *l*>10^65^-10^7^ bp due to approaching the upper cut-off, with the exception of chromosomes 22, X, and Y, where the differences appearing at *l* already >10^3^ bp pointing to a general bigger difference in the general sequence organization in respect to the other chromosomes. This special behaviour as well as the general scaling behaviour is nearly identical for the two human strains *Homo sapiens* and *Homo sapiens GRCH37,* thus this is a general feature of those chromosomes across species.

*SFigure 20:*

Detailed multi-scaling behaviour of the correlation coefficient δ(*l*) and its fine-structural features for *Mus musculus* and *Mus musculus C57BL6j:* The correlation coefficient δ(*l*) shows strong positive correlations for human chromosomes (A-G). In general, δ(*l*) increases from a starting value until a plateaued maximum from 10^2^-10^36^ bp, before a decrease and a second statistically significant maximum at ∼10^5^ bp for all chromosomes of both strains. Finally, δ(*l*) decreases to values characteristic for random sequences and enters the region of fluctuation. The differences between the strains within one species are mainly due to differences in the quality of sequences, e.g. unfinished/partial sequencing of the Y-chromosome, and also, but to a less degree, due to real sequence differences as genome rearrangements. Within this general behaviour, a distinct fine-structure visible less dominantly in the human compared to the mouse case is present in all chromosomes and survives averaging (Fig. S16B, S21). The very pronounced local maximum at 11 bp is related to the double helical pitch, whereas the local minima and maxima are related to the nucleosome, which is obvious for 146 bp, but less obvious for other positions. The second maximum around 10^5^, shows also a fine-structure which is due to the individual chromatin loops of the three-dimensional genome organization and their grouping in aggregates/rosettes.

*SFigure 21:*

The fine-structural features of *Homo sapiens* and *Homo sapiens GRCH37* as well as *Mus musculus* and *Mus musculus C57BL6j*, survives averaging over all chromosomes (as previously shown (Fig. S16; and Knoch, 1998; Knoch et al., 1998; Knoch et al., 2002; Knoch, 2002; Knoch, 2003). The very pronounced local maximum at 11 bp is related to the double helical pitch in both species, whereas the local minima and maxima are very clearly related to the nucleosome in the much more pronounced human case, which is e.g. obvious for 146 bp, but less obvious for 172 bp, 205 bp, 228 bp and 248 bp. This fine-structure present in all human sequences is in agreement with the pattern found in simulations using a consensus nucleosomal binding sequence organized in a block/gene fashion and the positions of the local maxima are mostly the same as in the human genome, whereas the similarity of the position of the local minima is difficult to compare as they smear out in the human sequence due to the block structure of genomes (Knoch 1998; Knoch 2002). In mouse, the general behaviour of the fine-structure is different and not as pronounced compared to the human case, a close inspection reveals that many of the fine-structural peaks although small are also present, at least in individual chromosomal sequences. Thus consequently, the concentration of nucleosomal binding sites seem to be less and differently distributed in mouse compared to human sequences and also might have an evolutionary different survival time within the DNA sequence (Knoch et al., 1998; Knoch et al., 1998; Knoch et al., 2002; Knoch, 2002; Knoch, 2003).

## References

Barbieri, M., Chotalia, M., Fraser, J., Lavitas, L.-M., Dostie, J., Pombo, A., and Nicodemi, M. (2012). Complexity of chromatin folding is captured by the strings and binders switch model. Proc Natl Acad Sci USA 109, 16173–16178.

Baudy, P. and Bram, S. (1978) Chromatin fiber dimensions and nucleosome orientation: a neutron scattering investigation. Nucleic. Acids. Res. 5 (10), 3697–3714.

Baudy, P. and Bram, S. (1979) Neutron scattering on nuclei. Nucleic. Acids. Res. 6 (4): 1721–1729.

Baum, M., Erdel, F., Wachsmuth, M. and Rippe, K. (2014) Retrieving the intracellular topology from multi-scale protein mapping in living cells. Nat. Commun. 5, 4494–4506.

Belmont, A. S. and Bruce, K. (1994) Visualization of G1 Chromosomes: A Folded, Twisted, Supercoiled Chromonema Model of Interphase Chromatid Structure. J. Cell Biol. 127 (2), 287–302.

Belmont, A.S. (2001). Visualizing chromosome dynamics with GFP. Trends Cell Biol 11, 250–257.

Belmont, A.S. (2014). Large-scale chromatin organization: the good, the surprising, and the still perplexing. Curr Opin Cell Biol 26, 69–78.

Berezney, R., Dubey, D. D. and Huberman, J. A. (2000) Heterogeneity of eukaryotic replicons, replicon clusters, and replication foci. Chromosoma 108(8), 471–484.

Bickmore, W.A. (2013). The spatial organization of the human genome. Annu Rev Genom Hum Genet 14, 67–84.

Boveri, T. (1909) Die Blastomerenkerne von Ascaris meglocephala und die Theorie der Chromosomenindiviualität. Archiv für Zellforschung 3, 181–268.

Bulger, M. and Groudine, M. (2011) Functional and mechanistic diversity of distal transcription enhancers. Cell 144, 327–339.

Capoulade, J., Wachsmuth, M., Hufnagel, L. and Knop, M. (2011) Quantitative fluorescence imaging of protein diffusion and interaction in living cells. Nat. Biotechnol. 29(9), 835–839.

Carey, M., Leatherwood, J. and Ptashne, M. (1990) A potent GAL4 derivative activates transcription at a distance in vitro. Science 247(4943), 710–712.

Comings, D. E. (1978) Mechanisms of chromosome banding and implications for chromosome structure. Annu. Rev. Genet. 20, 440–460.

Comings, D. E. (1968) The rationale for an ordered arrangement of chromatin in the interphase nucleus. Am. J. Hum. Genet. 20, 440–460.

Cremer, C., Zorn, C. and Cremer, T. (1974) An ultraviolet microbeam for 257nm. Microskopy Acta 75, 331–337.

Cremer, T. and Cremer, C. (2001) Chromosome territories, nuclear architecture and gene regulation in mammalian cells. Nat. Rev. Genet. 2, 292–301.

Cremer, T. and Cremer, M. (2010) Chromosome territories. Cold Spring Harb Perspect Bio.

Cremer, T., Cremer, C., Baumann, H., Luedtke, E. K., Sperling, K., Teubner, V. and Zorn, C. (1982) Rabl’s model of the interphase chromosome arrangement, tested in chinese hamster cells by premature chromosome condensation and Laser-UV-Microbeam experiments. Human Genetics 60, 46–46.

Dekker, J., Rippe, K., Dekker, M. and Kleckner, N. (2002) Capturing chromosome conformation. Science 295, 1306–1311.

Dixon, J.R., Selvaraj, S., Yue, F., Kim, A., Li, Y., Shen, Y., Hu, M., Liu, J.S., and Ren, B. (2012). Topological domains in mammalian genomes identified by analysis of chromatin interactions. Nature 485, 376–380.

Dostie, J. and Dekker, J. (2007) Mapping networks of physical interactions between genomic elements using 5C technology. Nat Protoc 2, 988–1002.

Dross, N., Spriet, C., Zwerger, M., Muller, G., Waldeck, W., and Langowski, J. (2009). Mapping eGFP oligomer mobility in living cell nuclei. PLoS ONE 4, e5041.

Dubochet, J. (2012) Cryo-EM – the first thirty years. J. Microsc. 245(3), 221–224.

Eigen, M. and Winkler-Oswatitsch, R. Transfer-RNA: The early adapter. Naturwissenschaften 68, 217–228, 1981.

Eigen, M. and Winkler-Oswatitsch, R. Transfer-RNA, an early gene? Naturwissenschaften 68, 217–228, 1981.

Eltsov, M., Maclellan, K. M., Maeshima, K., Frangakis, A. S. and Dubochet, J. (2008) Analysis of cyro-electron microscopy images does not support the existence of 30-nm chromatin fibers in mitotic chromosomes in situ. PNAS 105(50), 19732–19737.

Erenpreisa, J. (1989) Large rossettes - the element of the suprachromonemal organisation of interphase cell nucleus. Proc. Latv. Acad. Sci. Ser. B 7, 68–71, (Russ).

Fejes-Toth, K., Knoch, T. A., Wachsmuth, M., Frank-Stöhr, M., Stöhr, M., Bacher, C. P., Müller, G. and Rippe, K. (2004) Trichostatin A induced histone acetylation causes decondensation of interphase chromatin. J. Cell Sci. 117(18), 4277–4287.

Finch, J. T. and Klug, A. (1976) Solenoidal model for the superstructure in chromatin. Proc. Natl. Acad. Sci. USA 73, 1897–1901.

Gerlich, D., Beaudouin, J., Kalbfuss, B., Daigle, N., Eils, R., and Ellenberg, J. (2003). Global chromosome positions are transmitted through mitosis in mammalian cells. Cell 112, 751–764.

Ghamari, A., van de Corput, M. P., Thongjuea, S., van Cappellen, W. A., van Ijcken, W. F. J., van Haren, J., Soler, E., Eick, D., Lenhard, B. and Grosveld, F. G. (2013) In vivo live imaging of RNA polymerase II transcription factories in primary cells. Genes Dev. 27(7), 767–777.

Giorgetti, L., Galupa, R., Nora, E. P., Piolot, T., Lam, F., Dekker, J., Tiana, G. and Heard, E. (2014) Predictive polymer modeling reveals coupled fluctuations in chromosome conformation and transcription. Cell 157(4), 950–963.

Görisch, S.M., Wachsmuth, M., Ittrich, C., Bacher, C.P., Rippe, K., and Lichter, P. (2004). Nuclear body movement is determined by chromatin accessibility and dynamics. Proc Natl Acad Sci USA 101, 13221–13226.

Hagège, H., Klous, P., Braem, C., Splinter, E., Dekker, J., Cathala, G., de Laat, W. and Forné, T. (2007) Quantitative analysis of chromosome conformation capture assays (3C-qPCR). Nat. Protoc. 2, 1722–1733.

Hanscombe, O., Whyatt, D., Fraser, P., Yannoutsos, N., Greaves, D., Dillon, N. and Grosveld, F. G. (1991) Importance of globin gene order for correct developmental expression. Genes Dev. 5(8), 1387–1394.

Ibel, K. (1982) Neutron diffraction of interphase nuclei. J. Mol. Biol. 160 (1), 77–85.

Iborra, F.J., Pombo, A., Jackson, D.A., and Cook, P.R. (1996) Active RNA polymerases are localized within discrete transcription “factories' in human nuclei. J Cell Sci. 109(Pt 6), 1427–1436.

Jhunjhunwala, S., van Zelm, M. C., Peak, M. M., Cutchin, S., Riblet, R., van Dongen, J. J. M., Grosveld, F. G., Knoch, T. A. and Murre, C. (2008) The 3D-structure of the Immunoglobulin Heavy Chain Locus: implications for long-range genomic interactions. Cell 133(2), 265–279.

Kepper, N., Foethke, D., Stehr, R., Wedemann, G. and Rippe, K. (2008) Nucleosome geometry and internucleosomal interactions control the chromatin fiber conformation. Biophys. J. 95(8), 3677–3691.

Knoch, T. A. (1998) Dreidimensionale Organisation von Chromosomen-Domänen in Simulation und Experiment. (Three-dimensional organization of chromosome domains in simulation and experiment.) TAK Press, Tobias A. Knoch, Mannheim, Germany, ISBN 3-00-010685-5.

Knoch, T. A., Waldeck, W., Müller, G., Alonso, A. & Langowski, J. DNA-Sequenz und Verfahren zur *in vivo* Markierung und Analyse von DNA/Chromatin in Zellen. German Patent Application 10013204.9-44 and International Patent Application PCT/DE01/01044.

Knoch, T. A., Münkel, C. and Langowski, J. (1998) Three-dimensional organization of chromosome territories and the human cell nucleus - about the structure of a self replicating nano fabrication site. Foresight Institute - Article Archive, Foresight Institute, Palo Alto, CA, USA, http://www.foresight.org, 1–6.

Knoch, T. A., Göcker, M. & Lohner, R. Methods for the analysis, classification and/or tree construction of sequences using correlation analysis. US Patent Application 60/436.056 and International Patent Application PCT/EP03/14854.

Knoch, T. A. (2002) Approaching the three-dimensional organization of the human genome: structural-, scaling- and dynamic properties in the simulation of interphase chromosomes and cell nuclei, long- range correlations in complete genomes, *in vivo* quantification of the chromatin distribution, construct conversions in simultaneous co-transfections. TAKPress, Tobias A. Knoch, Mannheim, Germany, ISBN 3-00-009959-X.

Knoch, T. A. (2003) Towards a holistic understanding of the human genome by determination and integration of its sequential and three-dimensional organization. High Performance Computing in Science and Engineering 2003, editors Krause, E., Jäger, W. & Resch, M., High-Performance Computing Center (HLRS) Stuttgart, University of Stuttgart, Springer Berlin-Heidelberg-New York, ISBN 3-540-40850-9, 421–440.

Knoch, T. A., Göcker, M., Lohner, R., Abuseiris, A. and Grosveld, F. G. (2009) Fine-structured multi-scaling long-range correlations in completely sequenced genomes - features, origin and classification. Eur. Biophys. J. 38(6), 757–779.

Knoch, T. A., Lesnussa, M., Kepper, F. N., Eussen, H. B., and Grosveld, F. G. (2009) The GLOBE 3D Genome Platform - Towards a novel system-biological paper tool to integrate the huge complexity of genome organization and function. Stud. Health. Technol. Inform. 147, 105–116.

Knoch, T. A. and Grosveld, F. G. Method for analysing the interaction of nucleotide sequences in a three-dimensional DNA structure. GB Patent Application GB20130020351 and International Patent Application WO2014IB02485 20141118.

Knoch, T. A. Simulation of different three-dimensional polymer models of interphase chromosomes compared to experiments. (submitted)

Knoch, T. A. Simulation of different three-dimensional polymer models of whole interphase nuclei compared to experiments. (submitted)

Kolovos, P., Knoch, T.A., Grosveld, F.G., Cook, P.R. and Papantonis, A. (2012) Enhancers and silencers: an integrated and simple model for their function. Epigenetics Chromatin 5, 1.

Kolovos, P., van de Werken, H.J., Kepper, N., Zuin, J., Brouwer, R.W., Kockx, C.E., Wendt, K.S., van, I.W.F., Grosveld, F., and Knoch, T.A. (2014). Targeted Chromatin Capture (T2C): a novel high resolution high throughput method to detect genomic interactions and regulatory elements. Epigenetics & chromatin 7, 10.

Kornberg, R. D. and Klug, A. (1981) The nucleosome. Scientific American 2, 28–44.

Lichter, P., Cremer, T., borden, J., Manuelidis, L. and Ward, D. C. (1988) Delineation of individual human chromosomes in metaphase and interphase cells by in situ suppression hybridization using recombinant DNA libraries. Hum. Genet. 80, 224–234.

Lieberman-Aiden, E., van Berkum, N.L., Williams, L., Imakaev, M., Ragoczy, T., Telling, A., Amit, I., Lajoie, B.R., Sabo, P.J., Dorschner, M.O., Sandstrom, R., Bernstein, B., Bender, M.A., Groudine, M., Gnirke, A., Stamatoyannopoulos, J., Mirny, L.A., Lander, E.S. and Dekker, J. (2009) Comprehensive mapping of long-range interactions reveals folding principles of the human genome. Science 326, 289–293.

Luger, C., Mäder, A. W., Richmond, R. K., Sargent, D. F. and Richmond, T. J. (1997) Crystal structure of the nucleosome core particle at 2.8 A resolution, Science 389, 251–260.

Maeshima, K., Hihara, S., and Eltsov, M. (2010). Chromatin structure: does the 30-nm fibre exist in vivo? Curr Opin Cell Biol 22, 291–297.

Misteli, T. (2007). Beyond the sequence: cellular organization of genome function. Cell 128, 787–800.

Müller-Storm, H. P., Sogo, J. M. and Schaffner, W. (1989) An enhancer stimulates transcription in trans when attached to the promoter via a protein bridge. Cell 58(4), 767–777.

Müller, O., Kepper, N., Schöpflin, R, Ettig, R., Rippe, K. and Wedemann, G. (2014) Changing chromatin fiber conformation by nucleosome repositioning. Biphys. J. 107(9), 2141–2150.

Naumova, N., Imakaev, M., Fudenberg, G., Zhan, Y., Lajoie, B. R., Mirny, L. A. and Dekker, J. (2013) Organization of the mitotic chromosome. Science 342(6161), 948–953.

Notbohm, H. (1986) Small angle scattering of cell nuclei. Eur. Biophys. J. 13 (6), 367–372.

Olins, A. L and Olins, D. E. (1974) Spheroid Chromatin Units (v Bodies). Science 183, 330–332.

Osborne, C.S., Chakalova, L., Brown, K.E., Carter, D., Horton, A., Debrand, E., Goyenechea, B., Mitchell, J.A., Lopes, S., Reik, W., and Fraser P. (2004) Active genes dynamically colocalize to shared sites of ongoing transcription. Nat Genet. 36(10), 1065–1071.

Paulson, J. R. and Laemmli, U. K. (1980) The structure of histone-depleted metaphase chromosomes. Cell 12, 817–828.

Pienta, K. J. and Coffey, D. S. (1984) A structural analysis of the role of the nuclear matrix and DNA loops in the organization of the nucleus and chromosome. J. Cell. Sci. Suppl. 1, 123–135.

Pope, B.D., Ryba, T., Dileep, V., Yue, F., Wu, W., Denas, O., Vera, D.L., Wang, Y., Hansen, R.S, Canfield, T.K., Thurman, R.E., Cheng, Y., Gülsoy, G., Dennis, J.H., Snyder, M.P., Stamatoyannopoulos, J.A., Taylor, J., Hardison, R.C., Kahveci, T., Ren, B., and Gilbert DM. (2014) Topologically associating domains are stable units of replication-timing regulation. Nature 515(7527), 402–405.

Rabl, C. (1885) Über Zellteilung. Morphologisches Jahrbuch 10, 214–330.

Rao, S. S. P., Huntley, M. H., Durand, N. C., Stamenova, E. K., Bochkov, I. D., Robinson, J. T., Sanborn, A. L., Machol, I., Omer, A. D., Lander, E. S., and Lieberman-Aiden, E. (2014) A 3D map of the human genome at kilobase resolution reveals principles of chromatin looping. Cell 159, 1–16.

Rauch, J., Knoch, T. A., Solovei, I., Teller, K. Stein, S., Buiting, K., Horsthemke, B., Langowski, J., Cremer, T., Hausmann, M. and Cremer, C. (2008) Lightoptical precision measurements of the Prader- Willi/Angelman Syndrome imprinting locus in human cell nuclei indicate maximum condensation changes in the few hundred nanometer range. Differentiation 76(1), 66–82.

Reznik N. A., Yampol, G. P., Kiseleva, E. V., Khristolyubova, N. B. and Gruzdev, A. D. (1990) Possible functional Structures in the chromomere, Nuclear Structure and Function. by Harris, J. R. & Zbarsky, I.B. (Editors), Plenum Press, New York - London, 27–29.

Sachs, R. K., van den Engh, G., Trask, B., Yokota, H. and Hearst, J. E. (1995) A random-walk/giant-loop model for interphase chromosomes. Proceedings of the Nat. Acad. Sci. USA 92, 2710–2714.

Simonis, M., Klous, P., Splinter, E., Moshkin, Y., Willemsen, R., de Wit, E., van Steensel, B. and de Laat, W. (2006) Nuclear organization of active and inactive chromatin domains uncovered by chromosome conformation capture-on-chip (4C). Nat Genet 38, 1348–1354.

Stadhouders, R., Kolovos, P., Brouwer, R., Zuin, J., van den Heuvel, A., Kockx, C., Palstra, R.J., Wendt, K.S., Grosveld, F.G., van Ijcken, W. and Soler, E. (2013) Multiplexed chromosome conformation capture sequencing for rapid genome-scale high-resolution detection of long-range chromatin interactions. Nat Protoc 8, 509–524.

Stehr, R., Schöpfling, R., Ettig, R., Kepper, N., Rippe, K. and Wedemann, G. (2010) Exploring the conformational space of chromatin fibers and their stability by numerical dynamic phase diagrams. Biophys. J. 98(6), 1028–1037.

Tolhuis, B., Palstra, R.J., Splinter, E., Grosveld, F. and de Laat, W. (2002) Looping and interaction between hypersensitive sites in the active beta-globin locus. Mol Cell 10, 1453–1465.

Verschure, P.J., van Der Kraan, I., Manders, E.M., and van Driel, R. (1999). Spatial relationship between transcription sites and chromosome territories. J Cell Biol 147, 13–24.

Vogel, F. and Schroeder, T. M. (1974) The internal order of the interphase nucleus. Humangenetik 25(4), 265–297.

Wachsmuth, M., Knoch, T. A. and Rippe, K. Dynamic properties of independent chromatin domains measured by fluorescence correlation spectroscopy in living cells. (submitted back to back with this submission).

Wachsmuth, M., Weidemann, T., Müller, G., Hoffmann-Rohrer, U. W., Knoch, T. A., Waldeck, W. and Langowski, J. (2003) Analyzing intracellular binding and diffusion with continuous fluorescence photobleaching. Biophys. J. 84(5), 3353–3563.

Weidemann, T., Wachsmuth, M., Knoch, T. A., Müller, G., Waldeck, W. and Langowski, J. (2003) Counting nucleosomes in living cells with a combination of fluorescence correlation spectroscopy and cofocal imaging. J. Mol. Biol. 334(2), 229–240.

Yokota, H., Singer, M. J., van den Engh, G. J. and Trask, B. J. (1997) Regional differences in the compaction of chromatin in human G0/G1 interphase nuclei. Chrom. Res. 5 (3), 157–166.

Yokota, H., van den Engh, G., Hearst, J., Sachs, R. K. and Trask, B. J. (1995) Evidence for the organization of chromatin in megabase pair-sized loops arranged along a random walk path in the human G0/G1 interphase nucleus. J. Cell Biol. 130 (6), 1239–1249.

Zuin, J., Dixon, J. R., van der Reijden, M. I. J. A., Ye, Z., Kolovos, P., Brouwer, R. W. W., van de Corput, M. P. C., van de Werken, H. J. G., Knoch, T. A., van Ijcken, W. F. J., Grosveld, F. G., Ren, B. and Wendt, K. S. (2014) Cohesin and CTCF differentitally affect chromatin architecture and gene expression in human cells. PNAS 111(3), 996–1001.

## Supplemental References

Bartek, J., Bartkova, J., Kyprianou, N., Lalani, E. N., Staskova, Z., Shearer, M., Chang, S. and Taylor-Papadimitriou, J. (1991) Efficient immortalization of luminal epithelial cells from human mammary gland by introduction of simian virus 40 large tumor antigen with a recombinant retrovirus. Proc. Natl. Acad. Sci. U S A 88, 3520–3524.

Bornkamm, G. W., Berens, C., Kuklik-Roos, C., Bechet, J. M., Laux, G., Bachl, J., Korndoerfer, M., Schlee, M., Hölzel, M., Malamoussi, A., Chapman, R. D., Nimmerjahn, F., Mautner, J., Hillen, W., Bujard, H. and Feuillard, J. (2005) Stringent doxycycline-dependent control of gene activities using an episomal one-vector system. Nucleic Acids Res 33(16), e137.

Borovik, A. S., Grosberg, A. Y. and Frank-Kamenetskii, M. D. (1994) Fractality of DNA Texts. J. Biomol. Structure 12(3), 655–669.

Chatzidimitriou-Dreismann, C. A. and Larhammar, D. (1993) Long-range correlations in DNA. Nature 361, 212–213.

Dixon, J. R., Selvaraj, S., Yue, F., Kim, A., Li, Y., Shen, Y., Hu, M., Liu, J. S. and Ren, B. (2012) Topological domains in mammalian genomes identified by analysis of chromatin interactions. Nature 485, 376–3.80.

Francke, U. (1994) Digitized and differentially shaded human chromosome ideograms for genomic applications. Cytogenet. Cell. Genet. 65, 206–219.

Kirkpatrick, S. and Stoll, E. (1981) Implementation of the R250 random number generator. J. Computational Physics 40, 517.

Li, W. (1997) The study of correlation structures of DNA sequences: a critical review. Comp. Chem. 21(4), 257–271.

Li, H. and Durbin, R. (2009) Fast and accurate short read alignment with Burrows-Wheeler transform. Bioinformatics (Oxford, England) 25(14), 1754–1760.

Li, H., Handsaker, B., Wysoker, A., Fennell, T., Ruan, J., Homer, N., Marth, G., Abecasis, G. and Durbin, R. (2009) The Sequence Alignment/Map format and SAMtools. Bioinformatics (Oxford, England) 25(16), 2078–2079.

Li, W., Marr, T. G. and Kaneko, K. (1994) Understanding Long-range correlations in DNA sequences. Physica D 75, 392–416.

Maier, W. L. (1991) A fast pseudo random number generator. Dr. Dobb’s Journal 176.

Mandelbrot, B. B. (1983) The Fractal Geometry of Nature, W. H. Freeman and Company, New York, ISBN 0-7167-1186-9.

Nativio, R., Wendt, K. S., Ito, Y., Huddleston, J. E., Uribe-Lewis, S., Woodfine, K., Krueger, C., Reik, W., Peters, J. M. and Murell, A. (2009) Cohesin is required for higher-order chromatin conformation at the imprinted IGF2-H19 locus. PLoS Genet 5(11), e1000739.

Peng, C. K., Buldyrev, S. V., Goldberger, A. L., Havlin, S., Sciortino, F., Simons, M. and Stanley, H. E. (1992) Long-range correlations in nucleotide sequences. Nature 356, 168–170.

Prabhu, V. V. and Claverie, J. M. (1992) Correlations in intronless DNA. Nature 359, 782.

Schockel, L., Mockel, M., Mayer, B., Boos, D. and Stemmann, O. (2011) Cleavage of cohesin rings coordinates the separation of centrioles and chromatids. Nat Cell Biol 13(8), 966–972.

Stanley, H. E., Buldyrev, S. V., Goldberger, A. L., Goldberger, Z. D., Havlin, S., Mantegna, R. N., Ossadnik, S. M., Peng, C. K. and Simons, M. (1994) Statistical mechanics in biology: how ubiquitous are long-range correlations? Physica A 205, 214–253.

Yunis, J. J. (1981) Mid-prophase human chromosomes. The attainment of 2000 bands. Hum. Genet. 56, 293–298.

